# Long metabarcoding of the eukaryotic rDNA operon to phylogenetically and taxonomically resolve environmental diversity

**DOI:** 10.1101/627828

**Authors:** Mahwash Jamy, Rachel Foster, Pierre Barbera, Lucas Czech, Alexey Kozlov, Alexandros Stamatakis, David Bass, Fabien Burki

## Abstract

High-throughput environmental DNA metabarcoding has revolutionized the analysis of microbial diversity, but this approach is generally restricted to amplicon sizes below 500 base pairs. These short regions contain limited phylogenetic signal, which makes it impractical to use environmental DNA in full phylogenetic inferences. However, new long-read sequencing technologies such as the Pacific Biosciences platform may provide sufficiently large sequence lengths to overcome the poor phylogenetic resolution of short amplicons. To test this idea, we amplified soil DNA and used PacBio Circular Consensus Sequencing (CCS) to obtain a ~4500 bp region of the eukaryotic rDNA operon spanning most of the small (18S) and large subunit (28S) ribosomal RNA genes. The CCS reads were first treated with a novel curation workflow that generated 650 high-quality OTUs containing the physically linked 18S and 28S regions of the long amplicons. In order to assign taxonomy to these OTUs, we developed a phylogeny-aware approach based on the 18S region that showed greater accuracy and sensitivity than similarity-based and phylogenetic placement-based methods using shorter reads. The taxonomically-annotated OTUs were then combined with available 18S and 28S reference sequences to infer a well-resolved phylogeny spanning all major groups of eukaryotes, allowing to accurately derive the evolutionary origin of environmental diversity. A total of 1019 sequences were included, of which a majority (58%) corresponded to the new long environmental CCS reads. Comparisons to the 18S-only region of our amplicons revealed that the combined 18S-28S genes globally increased the phylogenetic resolution, recovering specific groupings otherwise missing. The long-reads also allowed to directly investigate the relationships among environmental sequences themselves, which represents a key advantage over the placement of short reads on a reference phylogeny. Altogether, our results show that long amplicons can be treated in a full phylogenetic framework to provide greater taxonomic resolution and a robust evolutionary perspective to environmental DNA.

## Introduction

Sequencing of environmental DNA (eDNA) is a popular approach to study the diversity and ecological significance of microbial eukaryotes, including small animals, fungi, and protists. eDNA has catalyzed the discoveries of novel lineages at all taxonomic ranks from abundant to rare taxa, and revealed that most, if not all, known groups of microbes are genetically much more diverse than anticipated (de Vargas et al., 2015; Heger et al., 2018; Massana et al., 2015; Pawlowski et al., 2012). For protists, recent global molecular surveys revealed that they can account for up to 80% of the total diversity of eukaryotes in the environments (de Vargas et al., 2015; Logares et al., 2014; Massana et al., 2015; Pawlowski et al., 2012). Initially, these molecular environmental studies relied on cloning the small subunit ribosomal RNA gene (18S rDNA) followed by Sanger sequencing, thereby generating reads of sufficient length to enable reasonably accurate phylogenetic interpretation of the results (Amaral Zettler et al., 2002; Bass & Cavalier-Smith, 2004; Dawson & Pace, 2002; Diez et al., 2001; Edgcomb, Kysela, Teske, de Vera Gomez, & Sogin, 2002; Lopez-Garcia, Philippe, Gail, & Moreira, 2003; López-García, Rodríguez-Valera, Pedrós-Alió, & Moreira, 2001; Massana, Balagué, Guillou, & Pedrós-Alió, 2004; Massana, Castresana, et al., 2004; Moon-van der Staay, De Wachter, & Vaulot, 2001; Stoeck & Epstein, 2003; Stoeck, Taylor, & Epstein, 2003). Today, however, the overwhelming majority of eDNA data corresponds to much shorter reads produced by Illumina, which routinely generates several millions of reads (e.g. Bates *et al*., 2013; de Vargas *et al*., 2015; Geisen, 2016). This enables sequencing a large fraction of the species present in an environment, even the so-called ‘microbial dark matter’ of extremely rare organisms (de Vargas et al., 2015; Logares et al., 2014). The drawback of this method is that only genetic regions limited to a few hundred nucleotides (typically <500) can be sequenced at a time, for example the hypervariable V4 or V9 regions of the 18S rDNA or the internal transcribed spacer (ITS) (Mahé et al., 2015; Pawlowski et al., 2012; Stoeck et al., 2010).

Short amplicons contain relatively low phylogenetic signal, which can be a problem for identification especially when environmental reads are only distantly related to reference sequences. To address the issue of low phylogenetic signal in high-throughput data, a range of tools has been developed to provide reasonable taxonomic identification of environmental OTUs (Operational Taxonomic Units). Given the mass number of reads available, the most straightforward approach is to use pairwise sequence similarity searches against reference databases (de Vargas et al., 2015; Mahé et al., 2017). While fast, this approach is highly sensitive to the taxon sampling and annotation accuracy of the reference database. If a taxonomic group is absent or sequences are misannotated in the reference database, the corresponding queries will be only approximately annotated, remain unidentified, or worse, wrongly identified (Berger, Krompass, & Stamatakis, 2011). Recognizing the limitations of similarity-based methods, new tools have been developed that “place” short sequences into a phylogenetic context. The Evolutionary Placement Algorithm (EPA; implemented in RAxML, or more recently in EPA-ng) or pplacer are two such tools. They are becoming popular methods that use a reference tree of carefully selected (often long) sequences to successively score the optimal insertion position of every query sequence or OTU (Barbera et al., 2019; Matsen, Kodner, & Armbrust, 2010). These methods perform well, and have contributed to the discovery of novel eukaryotic lineages from environments where poor references exist (Bass et al., 2018; Mahé et al., 2017). However, the phylogenetic placement of short reads still requires the independent construction of a reference dataset, which by definition does not include the short reads themselves. Thus, methods like EPA rely on the availability of reference sequences generally produced by the less efficient and more expensive Sanger sequencing, or on genome or transcriptome sequencing projects.

To better exploit the phylogenetic signal of the rDNA operon in environmental metabarcoding studies, newer long-read sequencing technologies such as the Pacific Biosciences platform (PacBio) hold great promise. PacBio has lower throughput and higher error rates than Illumina but can produce reads that are over 20kb long at a fraction of the cost of Sanger sequencing. In the last two years, PacBio sequencing has started to be applied to metabarcoding studies, primarily on prokaryotic 16S rDNA (Mosher et al., 2014; Schloss, Jenior, Koumpouras, Westcott, & Highlander, 2016; Wagner et al., 2016) and most recently on larger amplicons also including the 23S rDNA (Martijn et al., 2017). For eukaryotes, the 18S rDNA was nearly fully sequenced for targeted microbial groups (Orr et al., 2018), whilst longer regions also spanning the ITS and the 28S gene were used to analyze fungal diversity (Heeger et al., 2018; Tedersoo, Tooming-Klunderud, & Anslan, 2018). These studies showed that in spite of the high error rates of PacBio, when applying a corrective process based on multiple sequence passes (Circular Consensus Sequences - CCS) together with rigorous quality filtering, long-amplicon sequencing is emerging as a robust approach for studying environmental diversity.

Here, we used soil eDNA samples to generate broad eukaryote amplicons of ca. 4500 bp spanning the 18S rDNA, ITS1, 5.8S, ITS2, and the 28S rDNA regions. We used PacBio-CCS to sequence these long-amplicons and applied several filtering steps to retain only high-quality sequences. We then followed a full phylogenetic workflow to accurately annotate long-sequences with taxonomy even in the absence of close references. These annotated sequences were combined with available references to infer a well-resolved global eukaryotic phylogeny from a concatenated 18S-28S alignment. Altogether, this study represents an important step forward to use the full power of phylogenetics to derive the accurate evolutionary origins of known and novel lineages present in the environment, as well as expanding rDNA sequence databases for metabarcoding of eukaryotes.

## Results

### Sequence curation

A total of 113,362 long rDNA Circular Consensus Sequences (CCS), all containing two or more passes, were generated with PacBio Sequel. These CCS reads were filtered by a series of stringent quality controls including the removal of non-specific amplicons and prokaryotic sequences, as well as chimera detection (Fig 1a; Supp. Fig 1a; see materials and methods). At the end of the curation pipeline, the amplicons had on average 9.95 CCS passes (stdev=2.9) (Supp. Fig 1b). The mean error rate was estimated to be 0.17% based on comparisons between CCS reads curated by our pipeline and known Sanger sequences of the same species of fungi (see materials and methods). OTUs were generated using a 97% similarity threshold based on the 18S region only, leading to 650 high-quality clusters after removing singletons. These OTUs ranged in length from 2501 to 5956 bp (Supp. Fig 1c). Most OTUs contained less than 10 reads, but some were much larger and likely represented the most abundant organisms in the samples; the largest OTU (6416 sequences) corresponded to *Brassica napus*, the main crop species cultivated in one of the samples, while the second largest OTU (1322 sequences) belonged to the gregarines (Apicomplexa), a paraphyletic group of parasites of various invertebrates that has been shown to be particularly abundant in some soil environments (Mahé *et al*., 2017).

**Figure 1.**
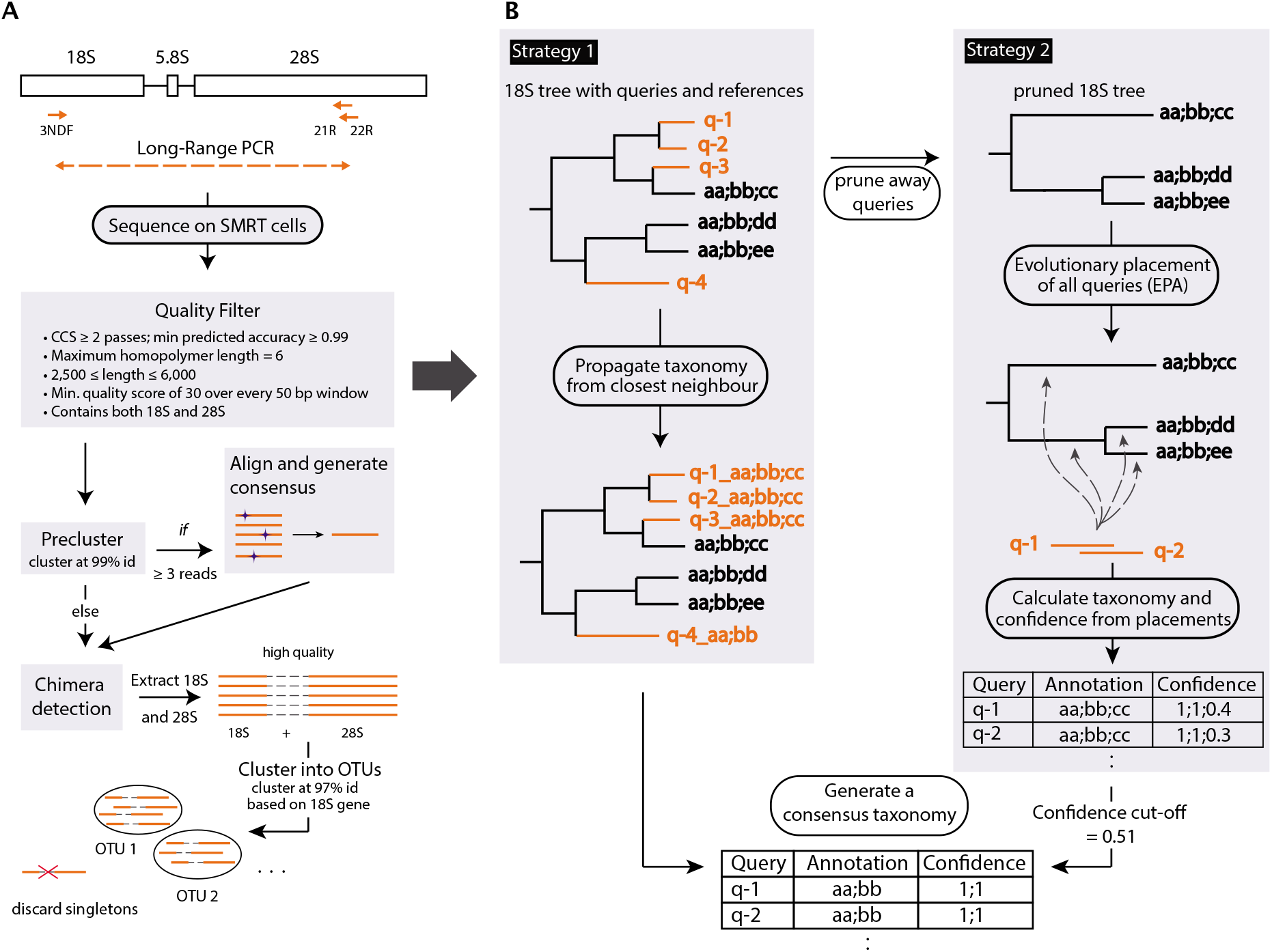
Workflow for generating long rDNA amplicons, quality filtering and taxonomic annotation. (A) General eukaryotic primers (3NDF, 21R and 22R) were used to amplify a ca. 4500 bp fragment of the rDNA operon (including 18S, ITS1, 5.8S, ITS2 and 28S) from environmental soil samples. The amplicons were sequenced on SMRT cells and subject to quality filtering before being pre-clustered at 99% similarity. Preclusters with at least 3 reads were de-noised further by aligning them and generating majority-rule consensus sequences. These were combined with the remaining reads and subject to *de-novo* chimera detection. The 18S and 28S gene regions extracted from these regions in the final step were physically linked as they originated from the same amplicon. (B) Overview of the taxonomic annotation pipeline. Reference sequences are shown in black with taxonomic ranks separated by ‘;’ (e.g. aa;bb;cc denote three taxonomic rank labels: aa, bb, and cc). Queries are shown in orange. See text for details on the pipeline.

### Phylogeny-aware taxonomic annotation

In order to annotate the environmental OTUs (hereafter referred to as queries) with taxonomy, we developed a phylogeny-aware approach that takes advantage of the increased sequence length. We used only the 18S part of the queries since the taxon sampling of 18S reference sequences is considerably denser than for 28S sequences. A phylogenetic tree was inferred using the 650 queries together with 1661 full-length references from the SILVA SSU database (i.e., a total of 2311 taxa). These references were manually selected to cover all major eukaryotic groups and, when available, included closely related sequences to the queries (Supp Fig 2). Based on this tree, a consensus taxonomy was derived using a combination of two strategies (Fig 1b). In the first strategy, the taxonomy of the most closely related reference was simply used as the taxonomy for the query. In cases where a query was most closely related to several references, the taxonomy was derived from the lowest shared taxonomic rank. In the second strategy, the queries were first removed from the tree before being placed back one at a time using EPA-ng (Barbera et al., 2019). The placements of the queries and their respective likelihoods were then used to calculate a confidence score for each taxonomic rank following the method implemented in SATIVA (Kozlov, Zhang, Yilmaz, Glöckner, & Stamatakis, 2016). The final consensus taxonomy for all queries was produced by merging the classifications from the two strategies only for the identical part of the taxonomic paths and when the SATIVA-derived confidence score for a rank was 0.51 or above (Supplementary Table1).

A key advantage of this phylogenetic approach is that it can assign taxonomy even in the absence of closely related reference sequences. This means that when queries branch off deeply in the reference tree, the taxonomic annotations are correctly derived from internal nodes — corresponding to higher taxonomic ranks — and not merely from the annotation of the most similar sequence (Supp. Fig 3b-d). Using this method, we were able to confidently assign a majority of queries (627/650, or 96.5%) to several major lineages of eukaryotes that we recognized to facilitate classification (Fig 2), including queries with similarity to references below 80%. The remaining 23 queries that were not assigned to any of the recognized major lineages were all highly-divergent; of these, 18 could be classified with confidence only to higher-rank assemblages that roughly correspond to the so-called supergroups — the most inclusive established groups of eukaryotes. For the remaining 5 queries, two were ambiguous even at the level of supergroups, thus possibly representing novel deeply-branching lineages and/or sparsely sampled taxonomic groups in the reference database, whilst three proved to be chimeras that had escaped automated filtering. Interestingly, our method also performed well for low-rank taxa, since 226 queries could be reliably annotated down to the genus and species levels.

**Figure 2.**
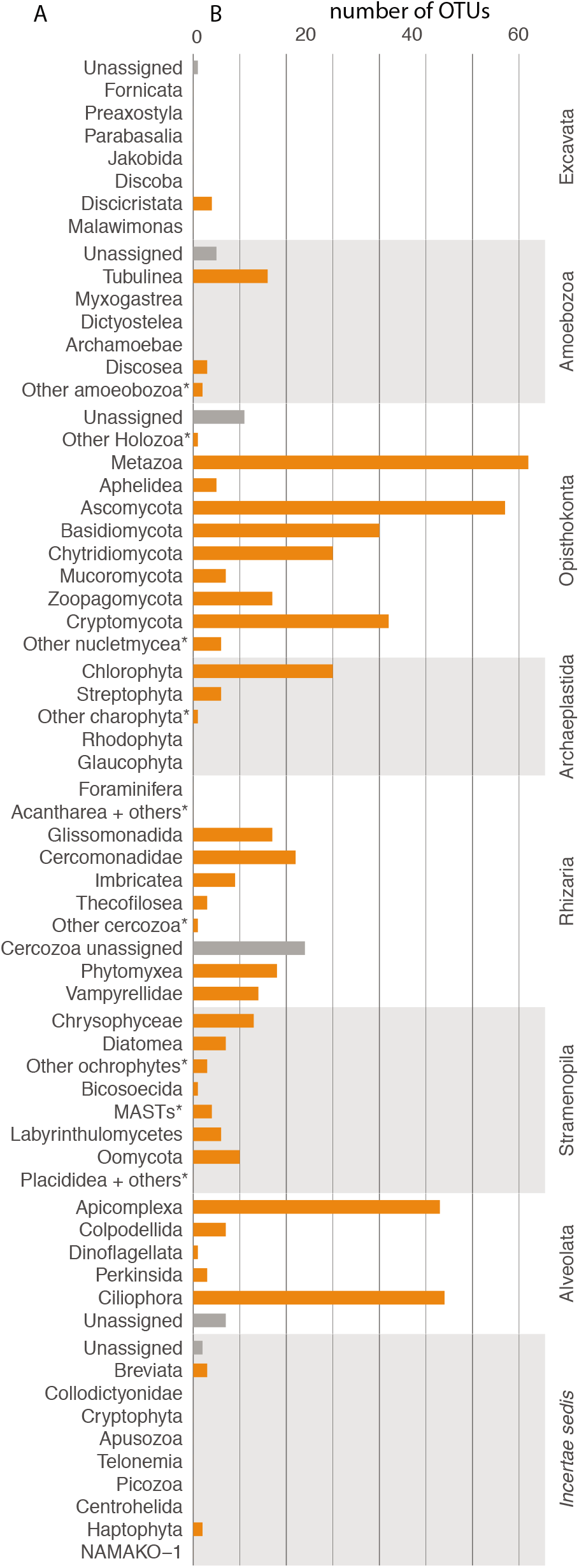
Taxonomic annotation of queries from PacBio sequencing of soil samples. (A) major eukaryotic lineages recognized to facilitate classification. (B) Number of queries assigned to each major lineage. Bars in grey highlight queries assigned to a supergroup or phylum, but not to any of the eukaryotic lineages. This may be for two reasons: (1) low confidence in assigning a sequence to a group or (2) reference sequences that are labelled only to a high taxonomic rank.

We further investigated the performance of our method by comparing it to a commonly used similarity-based taxonomic annotation tool (vsearch, Rognes *et al*., 2016), which revealed several discrepancies. Most importantly, 21 out of 48 queries (43.7%) with less than 80% similarity to references were assigned to different major lineages (those listed in Fig 2), and even to different supergroups in 4 cases. As expected, these discrepancies between major lineages became less pronounced for more similar sequences (i.e., between 80 and 90% similarity), where we observed only 9 (1.6%) conflicts, while there was no major conflict when queries were >90% similar to a reference. In general, the main differences between vsearch and our method occurred for relatively fast-evolving lineages such as Apicomplexa, or lineages with poor representation in the database such as the early-diverging fungi. One striking example illustrating the different behavior of the two methods in the absence of closely related references was for a newly proposed supergroup-level eukaryotic lineage called Hemimastigophora (Lax et al., 2018). One query with 85% sequence similarity to Streptophyta in SILVA, and thus annotated as a land plant by vsearch, was labelled as an “unidentified eukaryote” by our method. When using GenBank instead, this query was 98% similar to a newly added hemimastigote sequence, revealing that in the absence of an appropriate reference in SILVA for this group our method correctly proposed no specific annotation. We also observed other cases (37 in total) where queries were classified to one of the major lineages of eukaryotes (Fig 2) by vsearch, but our approach conservatively supported the taxonomy only to higher taxonomic levels.

Finally, we investigated the applicability of our method to short reads by truncating the long queries to the V4 region and repeating the taxonomic assignment algorithm. With the exception of one V4 query that was mis-assigned we found no conflicting taxonomic annotation between the long queries and the truncated V4 queries. However, the V4 queries were placed with overall lower confidence scores and 43 queries could not be confidently assigned to any taxonomic clade (Supp. Fig 4). This indicates, unsurprisingly, that the method performs sub-optimally when using shorter sequence lengths.

### Combined 18S-28S rDNA phylogeny of environmental DNA

The availability of long queries allows, in principle, to better resolve the origin of environmental sequences due to increased phylogenetic signal. We assembled a concatenated 18S-28S dataset including the annotated queries and reference sequences mined from various public databases. The references were selected such that it could be verified that both the 18S and 28S rDNA sequences originated from the same species (see material and method). We included representatives of all major eukaryotic lineages where possible. In addition, preliminary tree searches were used to identify long-branching taxa which were removed in downstream analyses to reduce potential long branch attraction artifacts. This yielded a final dataset of 1019 taxa, of which a majority (589 taxa = 58%) represented new environmental queries. Importantly, because the 18S sequences are physically linked to their 28S counterpart on the CCS reads, the taxonomic annotation inferred with our phylogeny-aware method could be unambiguously transferred to the combined 18S-28S reads. This provided a diverse set of taxonomically annotated environmental queries in otherwise sparsely populated reference sequences (Fig 3).

**Figure 3.**
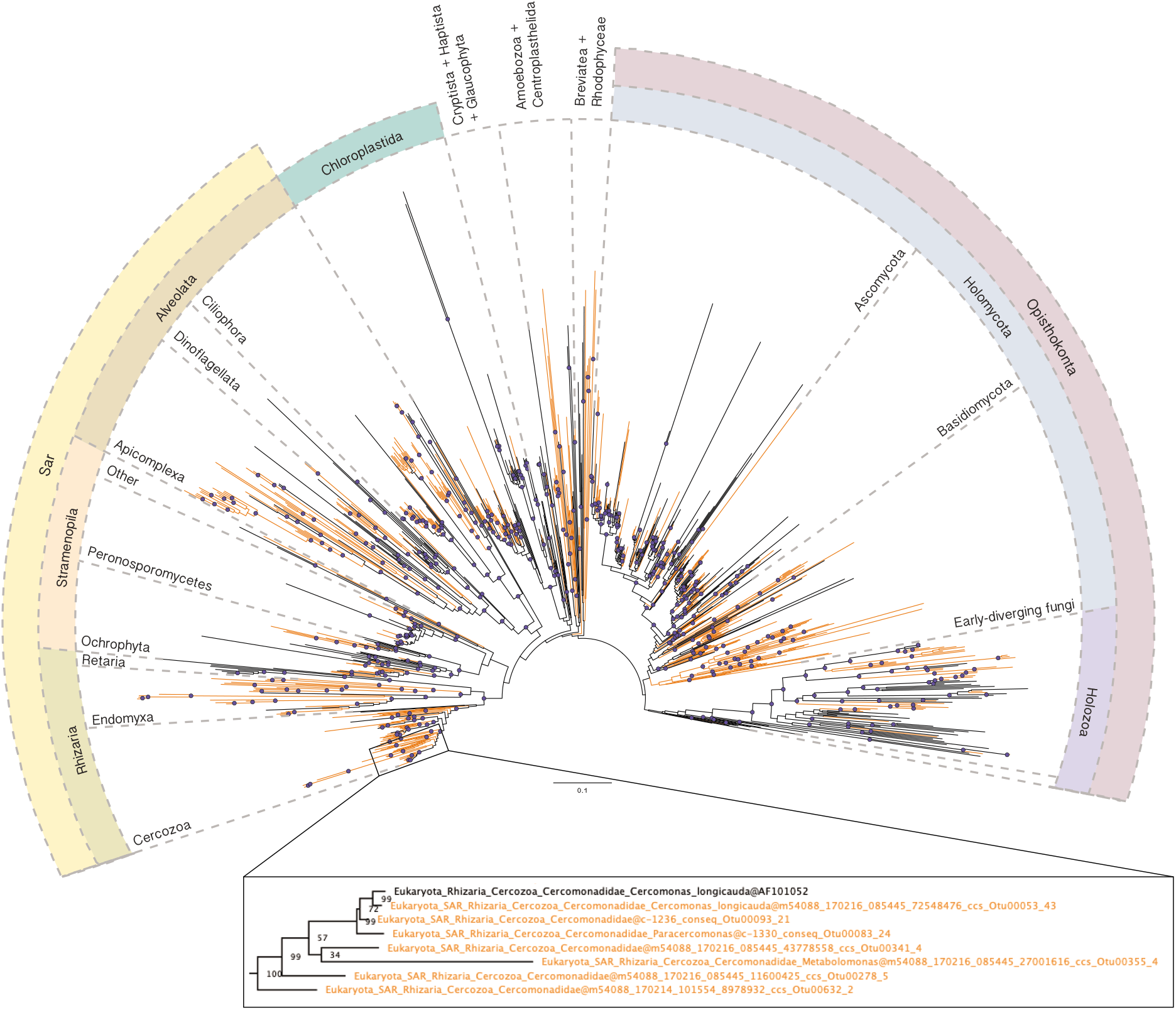
ML tree showing phylogenetic distribution of eukaryotic ribosomal diversity from soil samples. The tree corresponds to the best ML tree inferred from a concatenated alignment of 18S and 28S genes using the GTR+Gamma model. Bipartition support is derived from 300 bootstrap replicates. Purple circles represent BS ≥75%. The tree includes 589 queries from soil samples (coloured in orange) and 430 reference sequences (coloured in black) and all major eukaryotic supergroups (with the exception of Excavata which were eliminated during removal of long branches; see Methods for details). For details on taxon sampling, see Methods. The lower panel shows a subtree in Rhizaria. Zooming in illustrates how transferring the taxonomic annotation to the queries (coloured in orange) makes the phylogeny more informative, especially in clades (such as Rhizaria) where there are few references (labelled in black) with both the 18S and 28S available.

Figure 3 shows a Maximum Likelihood (ML) tree of the 1019 taxa dataset. The phylogenetic relationships were in general agreement with previous phylogenies based on the 18S and 28S (Moreira et al., 2007; Zhao et al., 2012), even recovering several well-established supergroups that were first proposed based on substantially larger concatenated protein datasets such as Sar (including the subclades Stramenopila, Alveolata, and Rhizaria) or Opisthokonta (including the subclade Holomycota and Holozoa) (Baldauf, Roger, Wenk-Siefert, & Doolittle, 2000; Burki et al., 2007). Overall, more than half of the newly sequenced diversity (345 queries; 53% of all queries) corresponded to microbial taxa other than fungi or animals (Fig 3). Members of Alveolata and Rhizaria accounted for nearly 70% of these protist queries — the most dominant lineages in decreasing number of queries were Ciliophora, Apicomplexa, Cercomonadida, Phytomyxea, Glissomonadida, and Vampyrellida. The remaining sequenced diversity was dominated by fungal lineages, accounting for 203 queries (31% of all queries) that equally represented dikarya (Ascomycota and Basidiomycota) and the so-called early-diverging fungi (EDF). Of these EDF, Cryptomycota (microsporidia) and Chytridiomycota were particularly diverse. The remaining 16% of the queries corresponded to various animal lineages as well as land plants.

### Comparison to 18S-only and V4-based phylogenetic classification of environmental DNA

The combined 18S-28S tree described above (Fig 3) provides a new solution for obtaining a taxonomically annotated and well-resolved phylogenetic framework from high-throughput environmental sequencing. To assess to which extent the added information of the 28S gene improved the phylogenetic resolution, we first compared the combined tree to the 18S-only tree constructed for the taxonomic assignment. Interestingly, both trees were largely in agreement, suggesting that the ~1000bp-fragment sequenced for the 18S gene combined with the substantially denser reference sampling available for this gene provided sufficient phylogenetic signal to recover many groupings. However, the combined tree received generally higher bootstrap support values: 54.3% of the bipartitions (552/1016) received ≥ 75% bootstrap support in the combined tree compared to 43.1% bipartitions (994/2308) in the 18S tree. The combined tree also supported (bootstrap > 75%) more specific phylogenetic position for a few queries that were taxonomically annotated only to high-rank taxa based on the 18S tree. For example, two queries labelled as Opisthokonta could be assigned more precisely to Aphelidea and as sister to nucleariid in the combined 18S-28S tree, respectively; or one deep branching eukaryote in the 18S tree in fact corresponded to a long branch within Ascomycota in the combined tree.

To investigate the benefits of long reads for phylogeny-based resolution of environmental diversity in more detail, we constructed three additional datasets with varying sequence lengths focusing on the Apicomplexa. The sequence lengths corresponded to i) the combined 18S-28S alignment ii) the full-length 18S-only alignment, and iii) an alignment of full-length 18S reference sequences but with query sequences shortened to the V4 region. The taxon-sampling across the three datasets was identical to facilitate comparison, containing 67 queries and 40 reference sequences. The inspection of the combined and the 18S trees (Supplementary Figs 5-6) revealed no major discrepancies and placed all 56 queries among gregarines. As with the full eukaryotic tree, the bootstrap values were globally higher in the combined tree but many relationships remained unsupported. However, we observed several exceptions where the increased resolution of the combined tree allowed for better interpretation. Most importantly, the monophyly of Apicomplexa was statistically supported in the combined tree (83%) whereas it was unsupported in the 18S tree (23%). The same was observed for the monophyly of other established groupings, such as haemosporidians and piroplasmids (92% vs. 55% in the combined and 18S trees, respectively). Furthermore, the combined tree (Supp. Fig 5) resolved the neogregarines and eugregarines into separate clades with moderate to strong support (99% and 75%, respectively), while they did not form separate clades in the 18S tree as previously noted based on this gene only (Rueckert & Horák, 2017; Rueckert, Simdyanov, Aleoshin, & Leander, 2011). The comparison with the phylogenetic placement of the V4 query sequences revealed that EPA-ng successfully placed all apicomplexan queries among gregarines with the exception of one query, which was placed close to *Plasmodium* instead. However, the reference-only 18S tree had a different topology than the full 18S tree, presumably because the short queries were not used for inferring the latter tree (Supplementary Fig 7). Furthermore, a close inspection showed that three queries were probably misplaced by EPA-ng on the branch leading to the cephaloidophorids (parasites of marine invertebrates; Supp. Fig 7) when instead they formed a robust monophyletic clade putatively representing novel lineages on both the 18S and combined trees (Supp Fig 5 and 6).

## Discussion

In this study, we broadly sequenced the near-complete eukaryotic rDNA operon from environmental soil samples, using PacBio sequencing. To our knowledge this is the first long amplicon environmental sequencing study that uses a full phylogenetic approach to assess the diversity of all eukaryotes. To reduce the inherently high error rate of PacBio, we combined the Circular Consensus Sequencing (CCS) approach with a series of stringent filtering steps and clustering. The final error rate of our reads has a mean of 0.17%, which is comparable with the error rate of Illumina (0.21%; Schirmer, D’Amore, Ijaz, Hall, & Quince, 2016), or in other PacBio-based studies (Schloss et al., 2016; Tedersoo et al., 2018; Wagner et al., 2016). Even though the curating pipeline discarded the majority of CCS reads, many of which might still be of high quality, the 650 OTUs that passed all filtering steps comprised a large and broad diversity of eukaryotes. Almost all major microbial lineages were sampled, from known abundant taxa in soils such as Ciliophora, Cercozoa, Apicomplexa, and fungi, to rarer lineages in soil such as the mainly aquatic Bacillariophyceae (diatoms) and Chlorophyta (green algae) (Bahram et al., 2018; Foissner & W., 1987; Stefan Geisen et al., 2018, 2015; Stephen Geisen, Cornelia, Jörg, & Michael, 2014; Mahé et al., 2017). A few main protist groups lacked new OTUs altogether, including Cryptista, Retaria, Rhodophyceae, and Glaucophyta, but these are almost exclusively aquatic and thus less likely to be recovered among soil sequences even if present in the environment at very low abundance (de Vargas et al., 2015; Stefan Geisen et al., 2018; Lallias et al., 2015). However, not all major groups typically widespread in soils were recovered with a correspondingly high sequence diversity. This was for example the case for Amoebozoa, Excavata, or Centrohelida, whose low diversity might be at least partially explained by primer bias (see materials and methods), and/or by the ecological conditions represented by our samples.

The availability of longer environmental sequences opens up the possibility to phylogenetically resolve environmental diversity with improved accuracy. Previous studies employing both the 18S and 28S genes recovered many relationships within and between major eukaryotic groups with greater resolution than that afforded by the 18S alone (Moreira et al., 2007; Zhao et al., 2012). The use of both genes was proposed to more robustly derive the origin of environmental sequences, particularly in the case of fast-evolving taxa, but this was based on Sanger sequencing of clone libraries (Marande, López-García, & Moreira, 2009). Near full-length 18S amplicons and even longer fragments including parts of the 28S have also recently been sequenced with PacBio for group-specific investigations, demonstrating that long-read high-throughput sequencing is a promising alternative to Illumina for investigating the environmental diversity of eukaryotes (Heeger et al., 2018; Orr et al., 2018). Here, we extended the approach to ~4500bp of the rDNA operon across the whole phylogenetic diversity of eukaryotes. We built a combined 18S-28S tree of eukaryotes that is globally well-resolved and can serve as a robust phylogenetic framework to describe the environmental diversity in our samples (Figure 3). Comparisons to the 18S region alone of our queries (~1200 bp) provided a similar overall topology to the combined tree, but with lower overall resolution (Supp Fig 2). Furthermore, some key groups in the apicomplexan phylogeny were either missing or not supported by the 18S-only tree. Altogether, our phylogenetic comparisons revealed that the 18S and 28S together provide increased resolution compared to the 18S alone, but the differences between single and two-gene trees vary across groups.

In order to assign taxonomy to the long environmental reads, we applied a novel phylogeny-aware approach that enables deriving robust annotation even in the absence of closely related references. Most commonly, taxonomic binning is conducted by similarity comparison to reference databases (e.g. in de Vargas *et al*., 2015; Mahé *et al*., 2017). Similarity works well when closely related references *are* available, however it requires the use of arbitrary similarity cutoffs without biological grounding below which sequences are considered of unknown origins (Bahram et al., 2018; Stoeck et al., 2010). To enable the use of phylogenetics with short environmental reads, methods such as the Evolutionary Placement Algorithm (EPA) have been recently developed and successfully applied to microbial diversity (Bass et al., 2018; Mahé et al., 2017). Whilst the need for similarity cutoffs is alleviated, the EPA still requires longer reference sequences to build a stable evolutionary framework and thus does not fully overcome the limitations of short read sequencing when references are lacking. Whilst our method relies partly on the EPA, it makes explicit use of environmental sequences to build a reference tree and computes a confidence score for each taxonomic rank. We show that it provides accurate taxonomic annotation with ranks corresponding to the phylogenetic position of queries in the reference tree—higher ranks correspond to deeper branches in the phylogeny—and that it performs better than similarity-based methods for divergent sequences (≤90% similarity). Comparison with the classical use of EPA with V4 reads revealed that whilst the overall annotations were similar, our approach utilizing long queries led to higher confidence scores. It was also more informative than placing short reads on a reference phylogeny, because the long queries directly contributed to the phylogenetic inference by filling gaps between references. Thus, the relationships between the queries themselves can be determined to reveal whether they cluster around known sequences or form entirely new clades.

One of the main benefits of our approach is that it provides both the 18S and 28S genes for the *same amplicon*. The 18S gene has long been the reference molecular marker for environmental studies of protist diversity (de Vargas et al., 2015; Diez et al., 2001; López-García et al., 2001; Massana, Balagué, et al., 2004; Massana et al., 2015; Moon-van der Staay et al., 2001). With our approach, we guarantee that each 18S sequence is paired with its 28S counterpart. As a result, we rapidly generated a massive increase of 28S sequence diversity for which the attached 18S provides a direct link to the much larger availability of 18S sequences contained in databases such as SILVA, PR2, or GenBank. As a point of comparison, the new sequences produced in this study alone represented the majority (58%) of all broad eukaryote diversity for which we could gather reference sequences for both genes. At lower taxonomic ranks, the increase in sequence diversity can be even more significant. For example, we found a total of only nine species of gregarines (Apicomplexa) that have both 18S and 28S genes in all public databases. In this study, we obtained 56 new gregarine OTUs, corresponding to a 6-fold increase in diversity for this group. Thus, we suggest that the newly generated long environmental sequences can be used in future studies as taxonomically-annotated “anchor” sequences to fill phylogenetic gaps in addition to the more traditional Sanger reference sequences.

In conclusion, we demonstrate several advantages of using high-throughput long sequence metabarcoding for environmental studies of microbial eukaryote diversity. With longer reads comes improved phylogenetic signal, and we show that it is possible to employ a full phylogenetic approach to taxonomically classify sequences and obtain a robust evolutionary framework of environmental diversity. This approach can be adapted for use with other emerging long-read technologies, e.g. Nanopore sequencing, and may prove particularly powerful in combination with higher-throughput sequencing technologies such as Illumina. Indeed, it will then be possible to map shorter but more abundant reads on a much more comprehensive reference phylogeny obtained from the same environments. The importance of eDNA studies continually grows in fields as varied as conservation biology, evolutionary biology and ecology. Long metabarcoding of the eukaryotic rDNA operon will undoubtedly play an increasingly important role in the close future.

## Materials and Methods

### Environmental samples

We used three environmental soil samples for this study: (1) soil from an alpine meadow in Tibet, China collected in summer 2011 (2) rhizosphere samples from rape seed samples from Newbald, Nuneaton, York and Morden in the UK, collected in March 2015 and, (3) pooled set-aside agricultural soils from Wellesbourne, UK, collected in September 2010. Rhizosphere samples were collected as follows: Loosely adhering soil was removed from the roots leaving no more than 2 mm rhizosphere soil. Roots were washed sequentially in 4 x 25 ml sterile distilled water to release the rhizosphere soil which was then centrifuged and the excess water drained to leave a pellet of rhizosphere soil. 250 mg of rhizosphere soil was extracted using the PowerSoil 96 Well Soil DNA Isolation Kit (MoBio Laboratories, Carlsbad, CA, USA) following the manufacturer’s recommendations, except the samples were homogenised in a TissueLyser II (Qiagen) at 20 Hz for 2 x 10 minutes with a 180° rotation of the plates between homogenisations. Set-aside agricultural soils were collected as described in (Gosling, van der Gast, & Bending, 2017) and DNA was extracted using PowerSoil DNA Isolation Kit (MoBio Laboratories) as per manufacturers protocol, using a Precellys 24 homogenizer (Bertin Technologies) for the initial mechanical lysis step.

### PCR and PacBio sequencing

We used two sets of eukaryotic universal primers to amplify a region covering the 18S, ITS1, 5.8S, ITS2, and 28S. One 18S internal forward primer, 3NDf (Cavalier-Smith *et al*., 2009) (which anneals to the conserved region adjoining the 5’ end of the V4 region, *E. coli* position 505) was used in conjunction with two 28S internal reverse primers 21R *(E. coli* position 1926) and 22R (*E. coli* position 1952) (Schwelm, Berney, Dixelius, Bass, & Neuhauser, 2016) to amplify a ca. 4500 bp region. Taxon coverage of the primers was checked *in silico* using SILVA TestProbe 3.0 (Quast et al., 2013) primers 3NDf, 21R and 22R matched against 91.5%, 88.1% and 87.2% of all eukaryotic sequences in SILVA release 132 respectively.

PCRs were carried out using the Takara PrimeSTAR GXL high fidelity DNA polymerase in 25 μl reactions with 10-20 ng of template DNA. The following cycling conditions were used: denaturation at 98 °C for 10 s, primer annealing at 60 °C for 15 s, and extension at 68 °C for 90s. A final extension time of 60 s was used after 30 cycles. PCR products were purified by polyethylene glycol and ethanol precipitation and were pooled and concentrated using Amicon 0.5ml 50K columns (Merck, Germany). Amplicon sizes were checked using TapeStation (Agilent Technologies) before SMRTbell library preparations. Three SMRT cells (one per soil sample) on the PacBio Sequel instrument with v2 chemistry were used for sequencing. Additionally, one RSII SMRT cell was used to sequence the mock community. Sequencing was carried out at Uppsala Genome Center, Science for Life Laboratory, SE-75237 Uppsala.

### Mock community

We constructed a mock community of three fungal samples: two unidentified cultures (BOR77 and BOR79) as well as one known species (*Phaeosphaeria luctuosa*). We amplified the 18S gene using two sets of primers: AU2 and AU4 for BOR77 and BOR79 (Vandenkoornhuyse, Baldauf, Leyval, Straczek, & Young, 2002), and 3NDF and 1510R (Amaral-Zettler, McCliment, Ducklow, & Huse, 2009) for *Phaeosphaeria*. All PCRs were conducted in 20 μl final volumes with 1 μl of template DNA and a final concentration of 0.5 μM of each primer, 0.4 mM dNTPs, 2.5 mM of MgCl2, 0.2 mg bovine serum albumin (BSA), 1x Promega Green Buffer and 0.5 U of Promega GoTaq. Amplicons were sequenced with Sanger sequencing to obtain reference sequences against which our PacBio sequences could be compared. To assess error rate, curated PacBio sequences were searched against the 18S reference sequences with vsearch v2.3.4 (Rognes *et al*., 2016) using

~~~
vsearch --usearch_global --id 0.9 --strand both --maxaccepts 0 \
--top_hits_only --fulldp --userfields query+target+id+mism+gaps+tl
~~~

The error rate was calculated as (mismatches + indels)/length of target sequence.

### Sequence curation and clustering pipeline

To address PacBio’s high error rate, we used a stringent sequence curation pipeline (Supp. Fig 1). Circular Consensus Sequences (CCS) were generated from raw reads by SMRT Link v4.0.0.190159 using a minimum number of two passes and Minimum Predicted Accuracy of 0.99 with all other settings set to default. The latter was shown to be the most important factor in decreasing error rate by Schloss *et al*., 2016. We pooled sequences from the three samples at this stage to analyze them together, resulting in one fastq file. A fasta file was generated using the fastq.info command (pacbio=T) in mothur v1.39.5 (Schloss et al., 2009). Sequences at this stage of the pipeline are of generally high quality but still include non-specific PCR amplicons, PCR artifacts such as chimeras and sequencing errors such as long homopolymer runs. These were filtered out using the trim.seqs command in mothur using the following settings: minlength=2500, maxlength=6000 (to discard non-specific and incomplete PCR amplicons), maxhomop=6 (to discard sequences with a homopolymer run of more 6 nucleotides), and qwindowsize=50 and qwindowaverage=30 (to trim the few sequences with a stretch of low quality sequence). The remaining non-specific PCR amplicons were filtered out by using Barrnap v0.7 (--reject 0.4 --kingdom euk) (https://github.com/tseemann/barrnap) which predicts the presence and location of 18S and 28S genes in the sequences. Reads with unexpected structure (more than one 18S, 28S, 5.8S) or incomplete/non-specific reads (missing 18S and/or 28S) were discarded. An in-house perl script was used to identify sequences represented by reverse strand (using the Barrnap output) and subsequently reverse complement them so that all sequences are in the same direction.

The sequences were then denoised by pre-clustering as described in (Martijn et al., 2017). Briefly, sequences were clustered at 99% similarity using vsearch v2.3.4 (Rognes et al., 2016) (--cluster_fast --id 0.99). For each resulting pre-cluster with three or more reads, we aligned the reads with mafft v7.271 (--auto) (Katoh & Standley, 2013) and generated a majority-rule consensus sequence using the consensus.seqs (cutoff=51) option in mothur. Gaps were removed to yield final consensus sequences. This step primarily curates indels given that these sequencing errors are randomly distributed in PacBio sequences.

The denoised sequences as well as sequences from pre-clusters of size one and two were subjected to *de novo* chimera detection using Uchime (Edgar, Haas, Clemente, Quince, & Knight, 2011) (as implemented in mothur) (chunks=40, abskew=1; abundance of the denoised sequences was taken as the number of sequences in their respective pre-clusters). Our PCR primers amplified a few archaea ribosomal genes, and these were filtered out by removing sequences which BLASTed (Altschul, Gish, Miller, Myers, & Lipman, 1990) to prokaryotic sequences in the SILVA SSU Ref NR 99 database v132 (Quast et al., 2013).

Finally, we used in-house perl scripts to extract the 18S and 28S sequences from our cleaned reads, and aligned them with mafft-auto v7.271 (Katoh & Standley, 2013). Poorly aligned sequences were removed after manual inspection.

We clustered our 18S sequences at 97% similarity into Operational Taxonomic Units (OTUs) using an average-linkage hierarchical clustering method. This was done by first generating a distance matrix using the dist.seqs (cutoff=0.2) command in mothur and then clustering sequences using the cluster command. We used the get.oturep command (label=0.03, method=distance) in mothur to obtain as representative sequence of each OTU the sequence with the smallest distance to all other sequences in the cluster. We extracted the same sequences from the 28S sequence set as OTU representatives. From the total set of 1154 OTUs, we discarded all singletons to be conservative, and obtained 650 OTUs at the end of our pipeline.

### Taxonomic annotation

#### Phylogeny-aware annotation

We constructed an 18S tree, with both labelled references and our (yet unlabeled) queries, which was used as the basis for our taxonomic annotation pipeline. Known reference sequences (**RS**) were obtained from SILVA SSU Ref NR 99 release 132 (Quast et al., 2013). The reference sequence set comprised two subsets: (1) 504 RS representative of global eukaryotic diversity — these were derived from the 512 taxa dataset used in Mahe et al. 2017. (2) Two to five nearest neighbors of each query in the SILVA database. To obtain these, each query sequence was aligned (mafft --auto) with the top 50 BLAST hits respectively against high quality (pintail > 0) eukaryotic SILVA SSU sequences (50060 sequences) and pairwise ML distances were computed in RAxML (option -f x) under the GTR+GAMMA model of substitution (Stamatakis, 2014). Nearest neighbors were RS with the lowest pairwise maximum likelihood distances with the query. This resulted in 1157 RS after removing duplicates. These were then combined with the 504 sequences mentioned above for a total of 1661 RS. The final dataset thus comprised the 650 queries plus the 1661 RS (2311 sequences in total). The reference multiple sequence alignment (**MSA**) was built by aligning these sequences with mafft (--retree 2 --maxiterate 1000) and trimming with trimal (-gt 0.3 -st 0.001) resulting in 1589 alignment sites. The best unconstrained maximum likelihood (ML) tree was selected from 20 tree searches run using RAxML-NG (v. 0.6.0) (Kozlov, Darriba, Morel, & Stamatakis, 2018). We assumed that the SILVA taxonomy is correct and consistent with the exception of a few cases—preliminary tree searches detected several potentially mislabeled RS which were relabeled after careful inspection.

We derived a consensus taxonomy for each of our queries from two strategies (Fig 1b). Strategy 1: Using a custom program written with the Genesis library (https://github.com/lczech/genesis) to propagate the taxonomy of the closest related reference to each query. Specifically, the program propagates the taxonomic annotation up the tree (where one exists), solving conflicts at inner nodes by taking the intersection of the taxonomic annotation (i.e. lowest common ancestor). Once complete, it propagates that information down to the non-labeled taxa (queries). Strategy 2: Queries were first removed from the tree before being placed back one at a time using EPA-ng (v0.2.1-beta). The location and likelihood weights of the placements are then used to compute the taxonomic assignment and the confidence associated with each taxonomic rank as in SATIVA (Kozlov *et al*., 2016). This last step is implemented in the gappa tool “assign” (https://github.com/lczech/gappa). Finally, we used a custom perl script to calculate the union of taxonomic paths from strategy 1 and 2. (Supp. Table3; Text S1). Taxonomies assigned to each query were assigned to their 28S counterparts as they are physically linked on the same molecule.

#### Comparison with short reads

We evaluated the effect of query sequence length by running the taxonomic annotation pipeline with short Illumina reads that were generated *in silico*. We focused on the V4 region (~ 500 bp) of the SSU gene, which is commonly used in barcoding studies, most recently in Mahé *et al*., 2017. The new dataset (**MSA-V4**) was derived from the original MSA by using the V4 flanking primers, TAReuk454FWD1 and TAReukREV3 (Stoeck *et al*., 2010), to trim only the query sequences (median length ~ 340 nucleotides), leaving the rest of the MSA untouched. After running the taxonomic annotation pipeline, we performed the following analyses:

1. The accuracy of placement is crucial for correct taxonomic annotation and we compared that for the long and short queries using two metrics, as shown in Supplementary Figure 4. (i) LWR (likelihood weight ratio) of the most probable placement for each query — this is computed as the ratio of the likelihood of the tree with the query at branch x to the sum over the likelihoods of all other possible placements. While a low LWR generally indicates that the placement is uncertain, this is not necessarily always the case; it may also arise if there are several equally likely placements in the same neighborhood of the phylogeny, thus dividing the probability between them. For example, a tree with multiple similar reference sequences for the same taxon might induce such locally distributed, yet high overall and clade-specific, placement probabilities. (ii) EDPL (Expected Distance between Placement Locations) shows how far the placements are spread across the tree. It is computed as the sum of the distances between placements along the branches of the tree, weighted by their probability (LWR). A low EDPL score indicates local placement uncertainty while a large score indicates global uncertainty. In other words, the EDPL shows how concentrated the placements of a query sequence are on the tree, with low values indicating placements in several close-by neighboring branches, and high values indicating a spread of the placements across the tree.
2. Lastly, we conducted pairwise comparisons of the taxonomic assignments and the confidence for taxonomic ranks given to each query based on **MSA** and **MSA-V4**.

#### Comparison with sequence similarity based methods of taxonomic assignment

Traditionally, environmental reads and/or OTUs are assigned to taxonomic groups based on their sequence similarity with sequences in a reference database. To assess how our method compares with such sequence similarity based approaches, we initially constructed a reference database consisting of high quality (pintail > 0) eukaryotic sequences in SILVA SSU Ref NR99 release 132 trimmed with the 3NDF primer, and the 504 RS derived from (Mahé *et al*., 2017). OTUs were searched against this reference set using the global pairwise alignment strategy in vsearch v2.3.4 (Rognes et al., 2016) with

~~~
vsearch --usearch_global --dbmask none --qmask none --rowlen 0 --
notrunclabels --userfields query+id1+target --maxaccepts 0 --maxrejects 32 --
top_hits_only --output_no_hits --id 0.5 --iddef 1.
~~~

Sequences were taxonomically assigned based on the top hit, and in case of multiple top hits, the common ancestor of the hits was computed.

### Phylogenetic analyses

#### 18S+28S phylogeny

To phylogenetically resolve the biodiversity in our soil samples, we constructed a phylogeny using a concatenated 18S + 28S dataset. For each of the two genes, OTUs were aligned with their respective reference sequences using mafft v7.271 (--retree 2 --maxiterate 1000). Alignments were filtered with trimal (-gt 0.3 -st 0.001) and a perl script (https://github.com/iirisarri/phylogm/blob/master/concat_fasta.pl) was used to concatenate the SSU and LSU alignments. The phylogeny was inferred with RAxML v8.2.10 as offered on the Cipres web server (Miller, Pfeiffer, & Schwartz, 2010), with 20 tree searches under the GTR+GAMMA model of substitution and 300 non-parametric bootstrap replicates. The construction of the reference dataset is described below.

We derived our references from several public databases. We included sequences for which we could easily verify that the 18S and 28S genes originated from the same species or organism. Reference sequences were obtained through several means, described as follows: (i) Searched NCBI nt using the following search filters: ((ribosomal RNA) AND 4000:9000[Sequence Length]) AND Eukaryota[Organism]. (ii) BLASTed our whole reads (18S, ITS, 28S) against nt and retained sequences with a minimum HSP of 2500 bp and 80% similarity. (iii) Obtained all 18S and 28S sequences from SILVA release 132 possessing the same accession number in the SSU Ref NR 99 and LSU Ref databases. (iv) Included the 108 taxa dataset used in an article studying the eukaryote tree with 18S+28S genes (Moreira et al., 2007). And lastly (v) used barrnap to search all “protist” genomes available in Ensembl Release 92 (Zerbino et al., 2018). This resulted in 3479 taxa after removing duplicates, from which we manually selected sequences, in a best effort, to assemble the most representative dataset possible. Initial tree building attempts placed certain cercozoan and apicomplexan lineages aberrantly among Excavata and Amoebozoa due to long branch attraction. We therefore sorted taxa by branch length (as in Heiss *et al*., 2018) and removed the longest 118 (10.4 %) branches from subsequent analyses. The final dataset contained 589 OTUs and 430 reference sequences with 4305 alignment sites.

#### Apicomplexa phylogenies

For the Apicomplexa phylogenies, we downloaded 40 GenBank 18S accessions for which 28 accessions had 28S accessions also available. For the concatenated and full-length 18S genes trees (Supp. Fig 5-6), the references and queries were aligned for each gene separately using mafft v7.271 (--linsi) and alignments filtered with trimal (-gt 0.3 -st 0.001). The trees were inferred with RAxML, using the substitution model GTR+GAMMA from 20 searches and 100 bootstrap runs. For the 18S tree with V4 queries, we first shortened the queries by trimming the queries in the 18S alignment with universal eukaryotic primers TAReuk454FWD1 and TAReukREV3 (Stoeck et al., 2010). The 18S reference tree was inferred as described above and short V4 amplicons were placed with EPA-ng.

### Data availability

Raw PacBio Sequel reads have been submitted to the ENA database under accession number PRJEB25197.

## Acknowledgements

We thank Stefan Geisen, Gary Bending and Sally Hilton at University of Warwick for providing soil samples. This work was supported by a grant from Science for Life Laboratory available to FB, which covered salary of MJ, and experimental expenses. The work of DB and RF was supported by the Standard Research Grant (NE/H009426/1), UK Department of Environment, Food and Rural Affairs (Defra) under contract FC1214. The authors would like to acknowledge support of the Uppsala Genome Center for providing assistance in massive parallel sequencing. Work performed at Uppsala Genome Center has been funded by VR and Science for Life Laboratory, Sweden.

