## Supplemental Figures and Table for "Long metabarcoding of the eukaryotic rDNA operon to phylogenetically and taxonomically resolve environmental diversity"

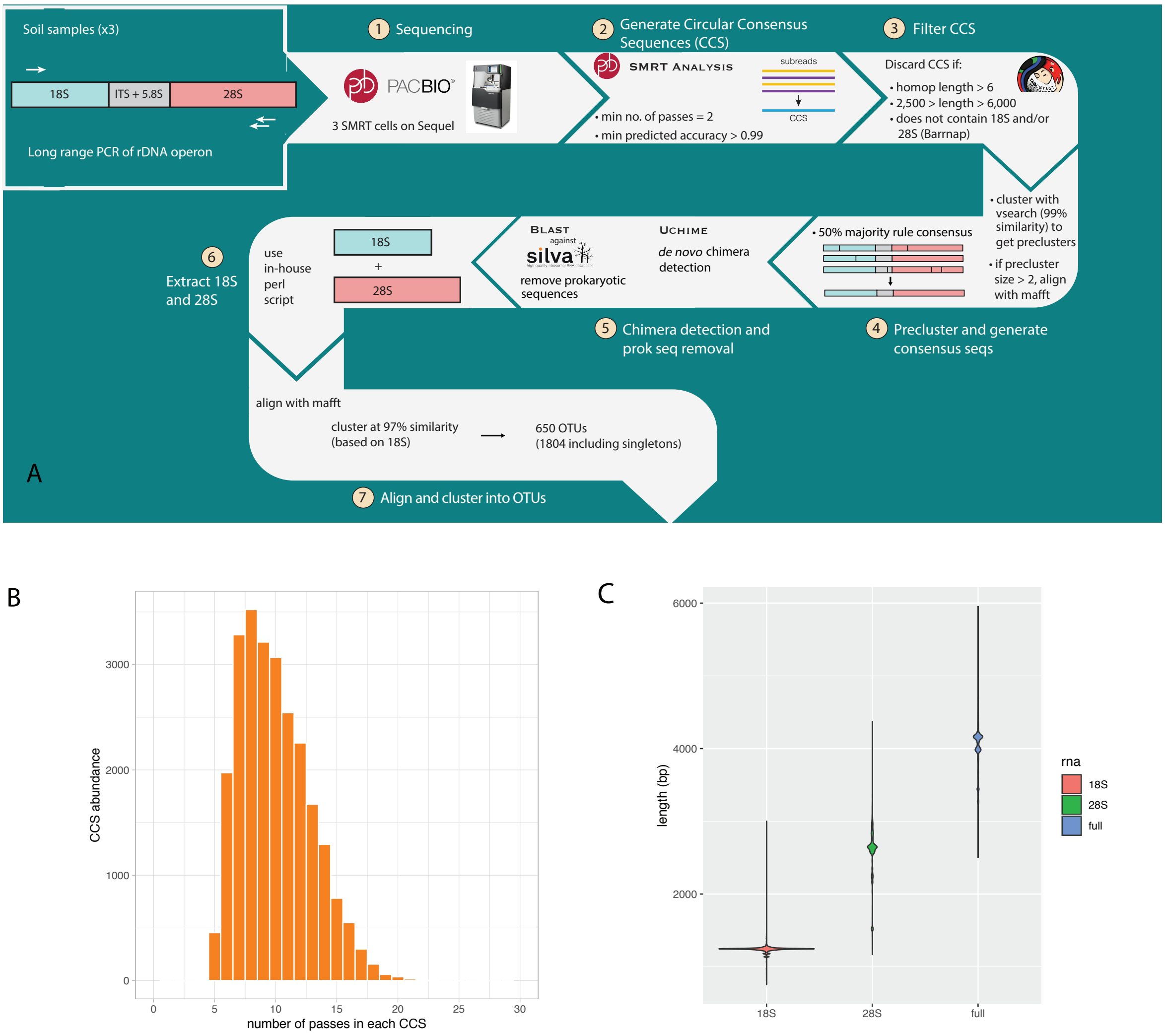


**Supplementary Fig 1.** (A) Overview of sequence curation pipeline for PacBio reads. (B) Distribution of number of passes in Circular Consensus Sequences (CCS) after curation of the sequences (i.e., after step 5 in the curation pipeline in panel A). As sequencing errors are randomly distributed in each pass, creating a consensus of each greatly reduces the error rate. A CCS generated from 10 passes has an accuracy rate of ~99.9%. (C) Violin plot showing the range of lengths (in bp) of the curated sequences for the 18S, 28S, and 18S-28S (including the ITS region).

**Supplementary Fig 2.** (on following page). ML tree generated for taxonomic annotation of queries. The tree was inferred from an alignment of the 18S gene (1589 bp) with 650 queries (branches in red) and 1661 reference sequences derived from the SILVA SSU database (branches in black; see Methods for details on taxon sampling). This best unconstrained tree was selected from 20 ML tree searches and bipartition support derived from 300 BS runs.

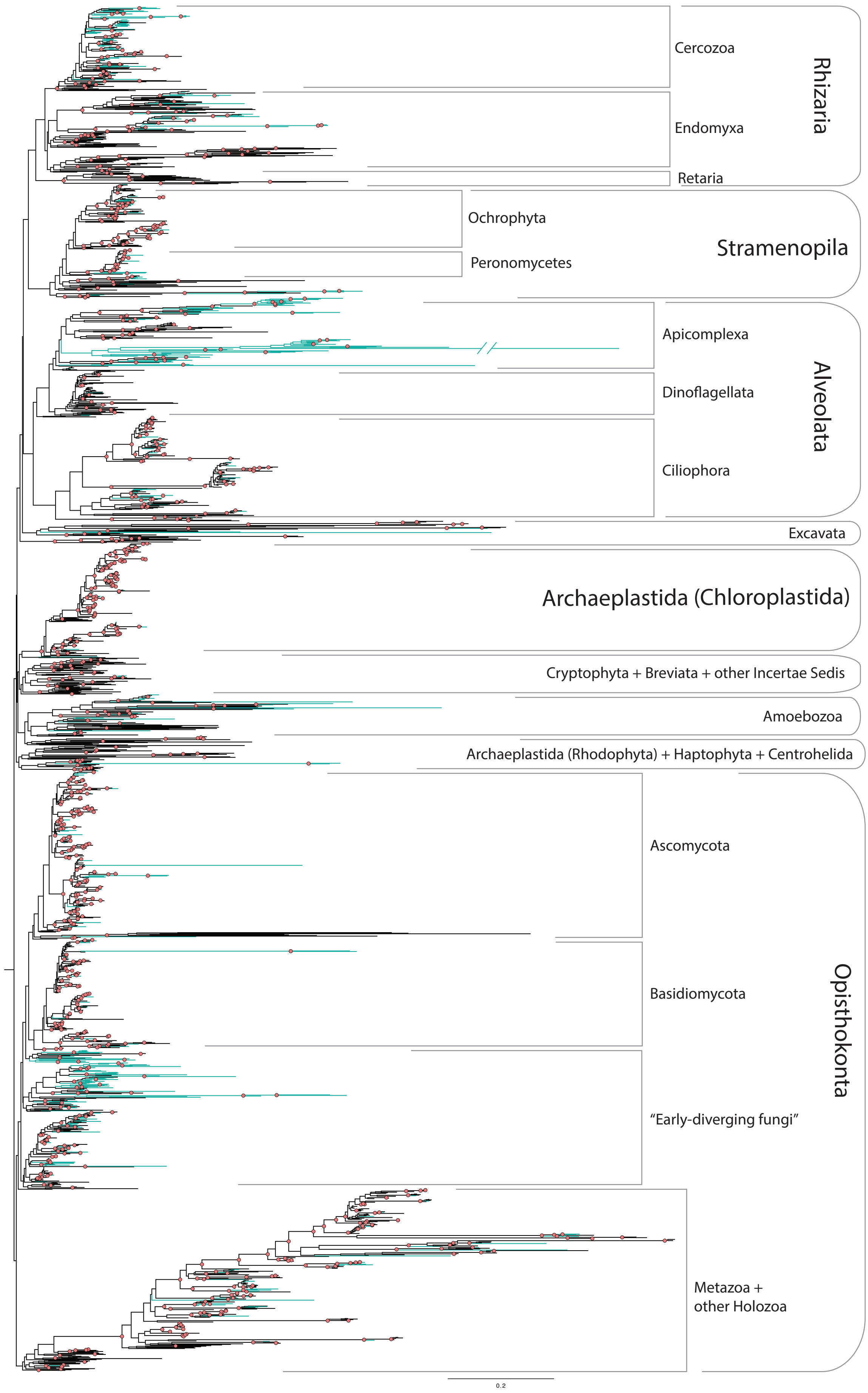
Cercozoa
Endomyxa
Retaria
Rhizaria
Ochrophyta
Peronomycetes
Stramenopila
Apicomplexa
Dinoflagellata
Ciliophora
Alveolata
Excavata
Archaeplastida (Chloroplastida)
Cryptophyta + Breviata + other Incertae Sedis
Amoebozoa
Archaeplastida (Rhodophyta) + Haptophyta + Centrohelida
Ascomycota
Basidiomycota
"Early-diverging fungi"
Metazoa + other Holozoa
Opisthokonta
0.2

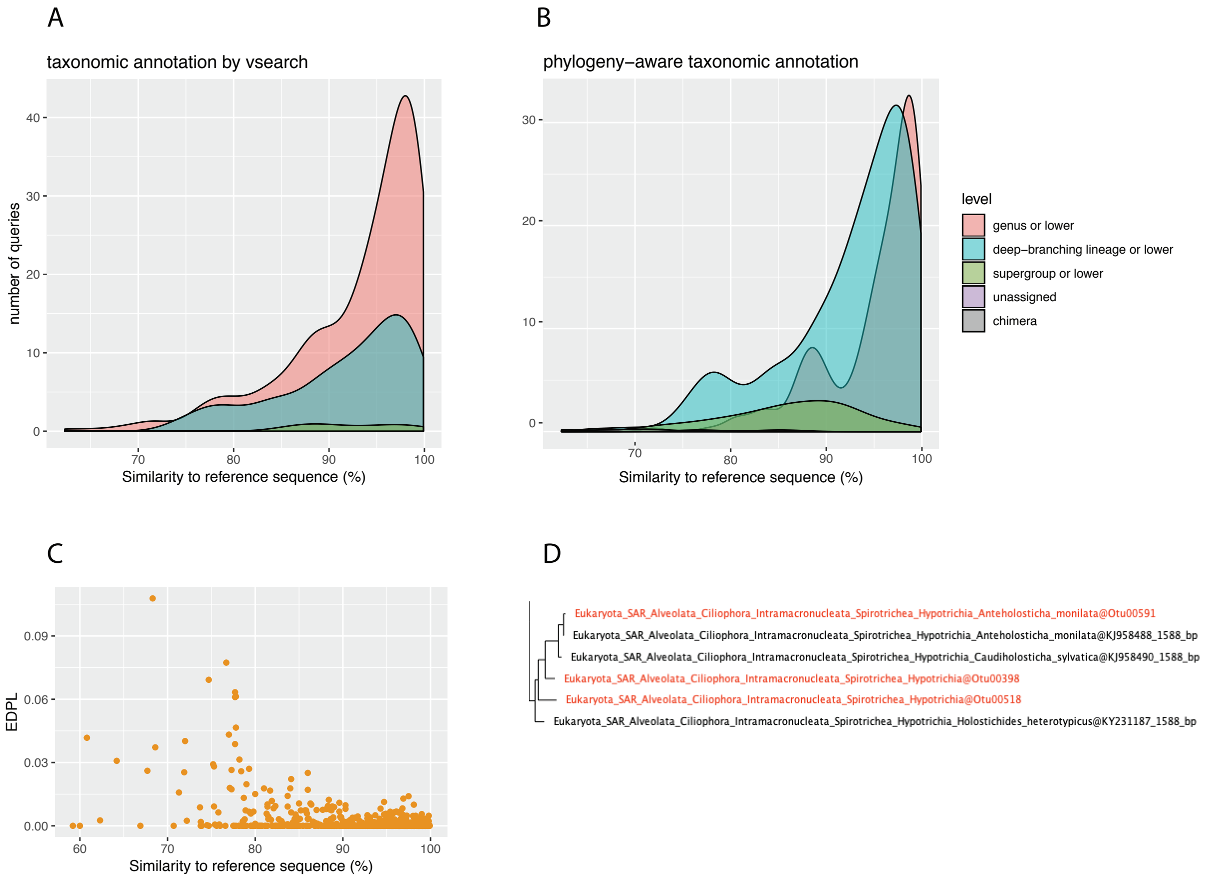


**Supplementary Fig 3.** Performance of our phylogeny-aware taxonomic annotation pipeline. (A) and (B) Density plots showing the lowest taxonomic rank assigned to each query based on their similarity to the closest reference. Pink corresponds to queries that were annotated down to either the genus or species level; cyan corresponds to queries that were annotated down to a major eukaryotic lineage (listed in Fig 2) or lower (family or sub-family); green corresponds to queries annotated to ranks above major eukaryotic lineages.
(A) Density plot when assigning taxonomy using a similarity-based annotation strategy (in this case, vsearch). There is no correlation between lowest taxonomic rank assigned to a query and its similarity to a reference, i.e. if the closest reference is labelled to species level, the query will also be classified down to species level regardless of whether the reference was identical or only 80% similar.
(B) In comparison, our phylogeny-aware annotation method results in a better correlation between taxonomic rank assigned and similarity to closest reference sequence.
(C) Scatterplot showing the EDPL (Expected Distance between Placement Locations) for queries placed on the pruned 18S tree with EPA (Evolutionary Placement Algorithm) in ‘Strategy 2’ of our taxonomic annotation pipeline. A high EDPL indicates that a query cannot be placed confidently in a location in the tree. The figure shows that the EPA assigns queries confidently to an evolutionary history even if there is no close reference sequence available. This can be seen from the low EDPL values even at low sequence identities.
(D) Zoomed in section of the 18S tree with reference tips labelled in black and query tips (in red) relabeled to their assigned taxonomy. The top-most query is correctly annotated down to the species level based on its position relative to its nearest neighbour. The other two are labelled only up to the family level because of their relatively deep-branching position in the tree.

A

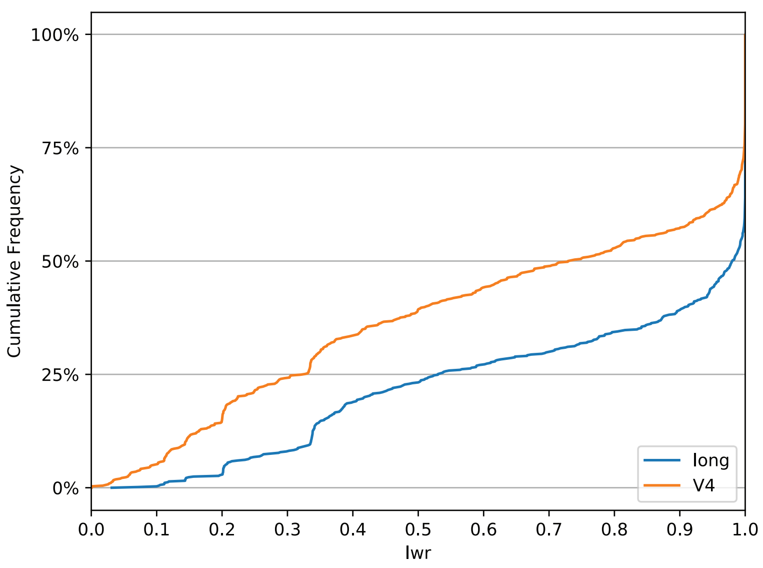


B

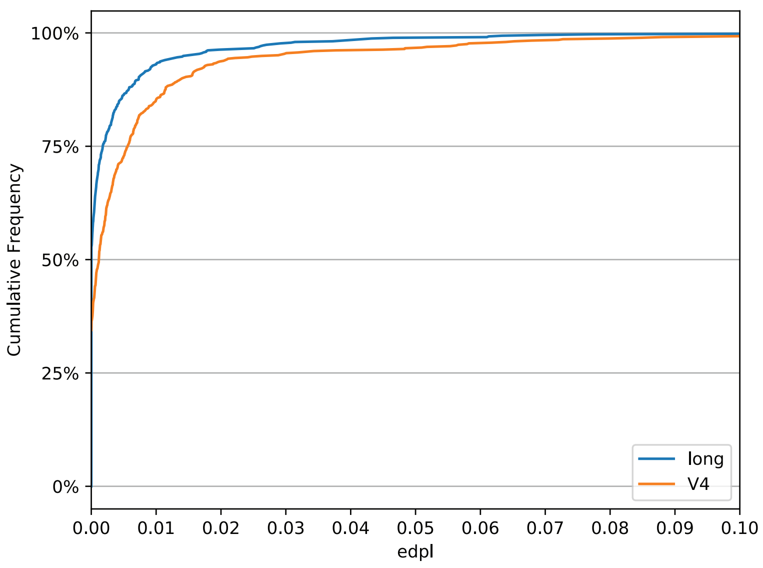


**Supplementary Fig 4.** Comparison of phylogenetic placement confidence for long (V4-end of 18S gene) vs. short queries (generated *in silico* by truncating to V4 region only). More long queries are placed with high confidence in the tree compared to V4 queries.
(A) The cumulative frequency plot shows the LWR value of the most likely placement of each query. At the far right, we see a steep rise to 100% caused by all the "remaining" queries, which have a LWR of exactly 1 (i.e perfectly confident placements). Here, the orange V4 line shows that it has fewer queries with the highest LWR possible.
(B) The EDPL cumulative frequency plot shows that the long reads start off at an EDPL value of 0 for more than 50% of the queries, while only ~35% of the V4 reads have an EDPL of 0. No queries (long or V4) have an EDPL higher than 0.1. Importantly, the long reads line is always ‘above’ the V4 line, meaning that the EDPL values are smaller for the long reads.

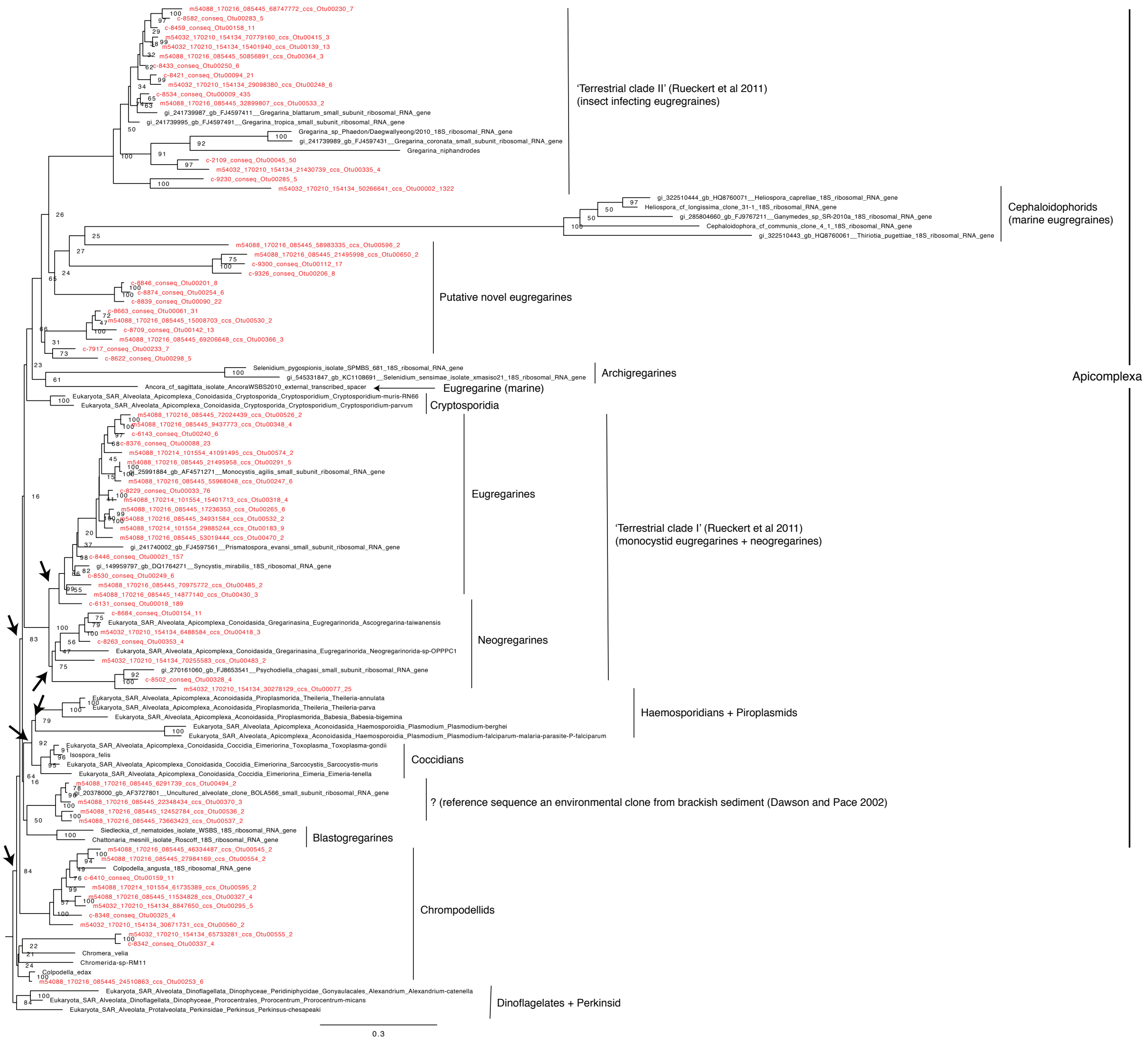


**Supplementary Fig 6-8**. Phylogenies of Apicomplexa and Chrompodellid queries (OTUs) found in soils, with particular focus on the gregarines, a paraphyletic group of obligate parasites in invertebrates. Gregarines were recently shown to be dominant in Neotropical soils (Mahé et al., 2017) and formed the second largest OTU from our soil samples (with 1322 sequences). All trees were inferred with the same taxon sampling. Reference sequence are black, query sequences are red.

**Supplementary Fig 6.** Phylogeny inferred from a concatenated alignment of 18S and 28S genes. Arrows point to strongly-moderately supported groups not statistically supported in the 18S tree (and in previously published 18S phylogenies; Rueckert & Horák, 2017; Rueckert, Simdyanov, Aleoshin, & Leander, 2011).

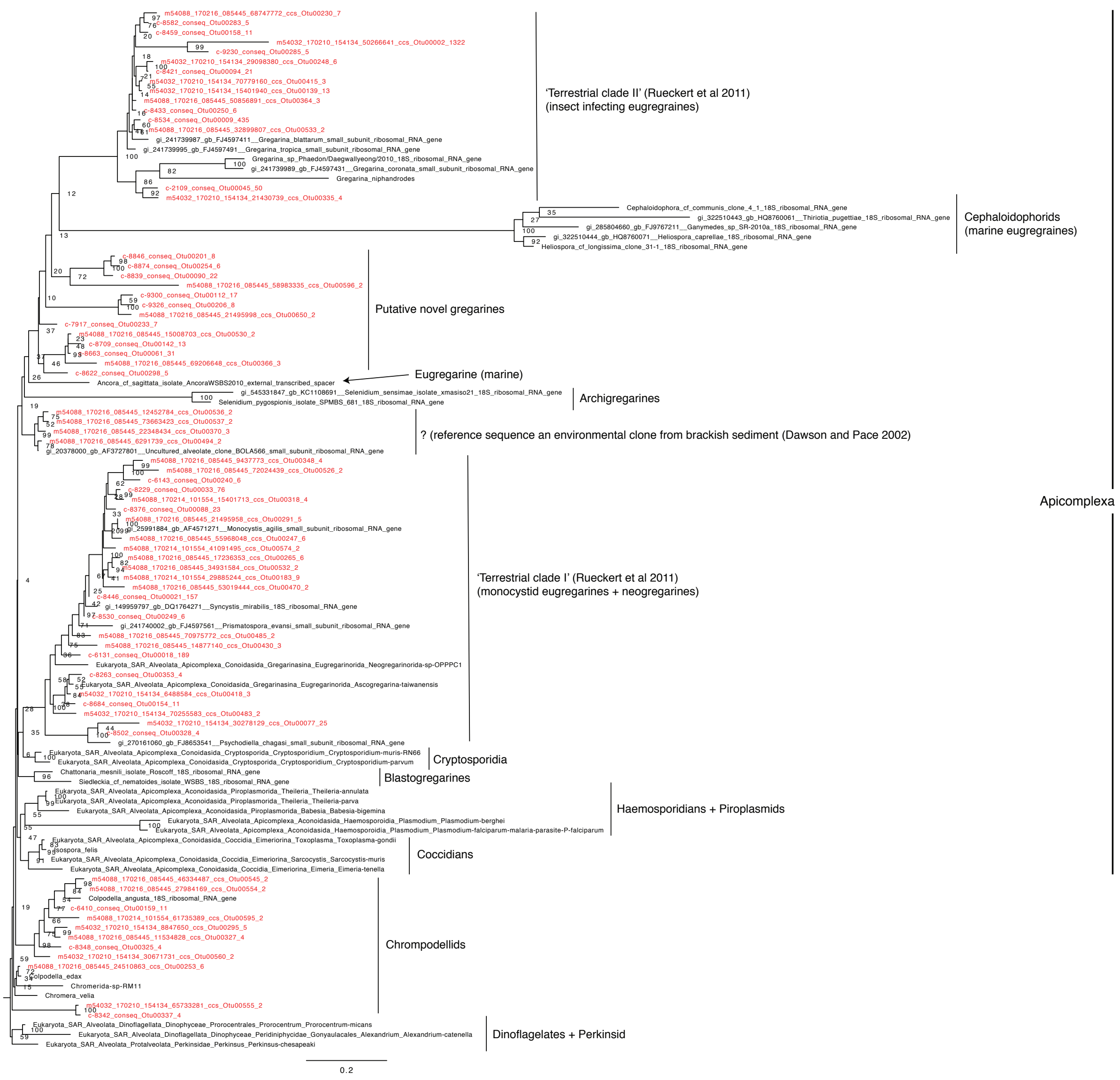


**Supplementary Fig 7.** Phylogeny inferred from an alignment of the 18S gene only. While the topology does not differ considerably from the concatenated tree, it has comparatively lower statistical support values.

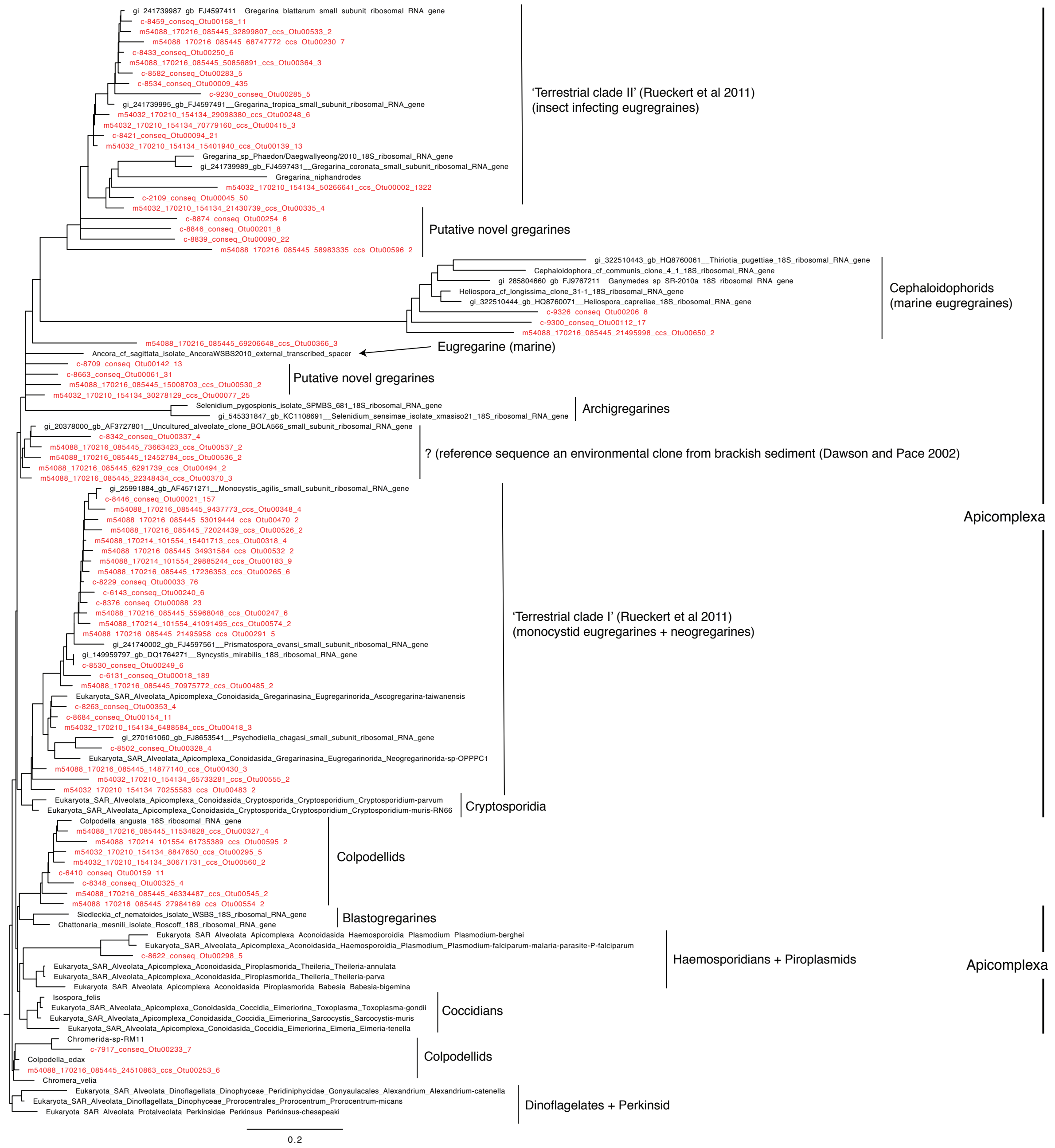


**Supplementary Fig 8.** Truncated V4 queries placed on the 18S tree inferred from references only. Here, the Apicomplexa is no longer monophyletic and several queries nest with a long branching group of marine eugregarines, a relationship that is not seen in the previous two phylogenies. This case is particularly interesting given that many OTUs from Neotropical soils (Mahé et al., 2017) were placed within similar marine lineages, and highlights the limitations of using short reads.

**Supplementary Table 1.** Table showing the annotation for each environmental query. "% similarity" refers to similarity to the closest reference in SILVA. Annotations in red indicate that the annotations were assigned by manual inspection. This occurred when a query was assigned with extremely low confidence, or if the annotations resulting from the two strategies (depicted in Fig1; main text) differed greatly.

| query | % similarity | annotation | confidence for each rank |
|---|---|---|---|
| c-1030_conseq_Otu00449_3_1588_bp | 93.1 | Eukaryota;Opisthokonta;Nucletmycea;Fungi;Chytridiomycota;Incertae_Sedis;Chytridiomycetes;Spizellomycetales;Olpidiaceae;Olpidium | 1;1;1;1;0.649691;0.649691;0.649691;0.649691;0.649691;0.649691 |
| c-1065_conseq_Otu00003_1281_1588_bp | 83 | Eukaryota;Opisthokonta;Nucletmycea;Fungi | 0.999712;0.999712;0.999712;0.999712 |
| c-1073_conseq_Otu00431_3_1588_bp | 98.8 | Eukaryota;SAR;Rhizaria;Cercozoa;Glissomonadida | 1;1;1;1;0.523693 |
| c-112_conseq_Otu00106_18_1588_bp | 92.8 | Eukaryota;Opisthokonta;Holozoa;Metazoa;Animalia;Eumetazoa;Bilateria;Tardigrada;Eutardigrada;Parachela | 0.998983;0.998983;0.998983;0.998983;0.998983;0.998983;0.998983;0.998533;0.998533;0.998533 |
| c-1164_conseq_Otu00032_77_1588_bp | 97.5 | Eukaryota;SAR;Rhizaria;Cercozoa;Thecofilosea;Cryomonadida;Rhizaspididae;Rhogostoma | 0.998411;0.998411;0.998411;0.998411;0.998411;0.998411;0.998411;0.998411 |
| c-1182_conseq_Otu00244_6_1588_bp | 98.8 | Eukaryota;Amoebozoa;Tubulinea;Arcellinida;Echinamoebida;Vermamoeba;vermiformis | 1;1;1;1;0.965796;0.965796;0.965796 |
| c-1236_conseq_Otu00093_21_1588_bp | 98 | Eukaryota;SAR;Rhizaria;Cercozoa;Cercomonadidae | 1;1;1;1;1 |
| c-1330_conseq_Otu00083_24_1588_bp | 97.6 | Eukaryota;SAR;Rhizaria;Cercozoa;Cercomonadidae;Paracercomonas | 1;1;1;1;1;0.999415 |
| c-1347_conseq_Otu00288_5_1588_bp | 83.2 | Eukaryota;Excavata;Discoba;Discicristata;Heterolobosea;Tetramitia;Tetramitus | 1;1;1;1;1;1;1 |
| c-1435_conseq_Otu00013_316_1588_bp | 99.2 | Eukaryota;Opisthokonta;Holozoa;Metazoa;Animalia;Eumetazoa;Bilateria;Nematoda;Chromadorea;Rhabditida | 0.999653;0.999653;0.999653;0.999653;0.999653;0.999653;0.999653;0.999653;0.999653;0.999653 |
| c-1449_conseq_Otu00044_50_1588_bp | 96.8 | Eukaryota;SAR;Rhizaria;Cercozoa;Vampyrellidae;Arachnula;impatiens | 1;1;1;1;1;0.903288;0.903288 |
| c-1565_conseq_Otu00190_9_1588_bp | 88.3 | Eukaryota;Opisthokonta;Nucletmycea;Fungi;Chytridiomycota;Incertae_Sedis;Chytridiomycetes;Incertae_Sedis | 0.999187;0.999187;0.999187;0.999187;0.999187;0.999187;0.999187; |
| c-1597_conseq_Otu00464_3_1588_bp | 86 | Eukaryota;Opisthokonta | |
| c-1639_conseq_Otu00133_13_1588_bp | 98.1 | Eukaryota;Opisthokonta;Nucletmycea;Fungi;Dikarya;Ascomycota;Pezizomycotina;Leotiomycetes;Helotiales | 0.929437;0.929437;0.929437;0.929437;0.929437;0.929437;0.929437;0.929437;0.870225 |
| c-1708_conseq_Otu00028_101_1588_bp | 99.6 | Eukaryota;Archaeplastida;Chloroplastida;Charophyta;Phragmoplastophyta;Streptophyta;Embryophyta;Bryophyta | 0.994739;0.994739;0.994739;0.994739;0.994739;0.994739;0.994739;0.993026 |
| c-1714_conseq_Otu00165_11_1588_bp | 83.9 | Eukaryota;Excavata;Discoba;Discicristata;Heterolobosea;Tetramitia;Vahlkampfia;inornata | 1;1;1;1;1;1;0.824585;0.824585 |
| c-1736_conseq_Otu00126_14_1588_bp | 93 | Eukaryota;Opisthokonta;Holozoa;Ichthyosporea;Ichthyophonae | 0.999809;0.999809;0.999809;0.999809;0.993632 |
| c-1764_conseq_Otu00216_8_1588_bp | 99.5 | Eukaryota;SAR;Stramenopiles;Ochrophyta;Eustigmatophyceae;Eustigmatales | 1;1;1;1;1;1 |
| c-1812_conseq_Otu00280_5_1588_bp | 96.1 | Eukaryota;SAR;Stramenopiles;Ochrophyta;Chrysophyceae | 0.999834;0.999834;0.999834;0.999834;0.999834 |
| c-1814_conseq_Otu00055_42_1588_bp | 99.7 | Eukaryota;Opisthokonta;Nucletmycea;Fungi;Dikarya;Basidiomycota;Agaricomycotina;Agaricomycetes;Agaricales;Tricholomataceae;Camarophyllopsis;schulzeri | 1;1;1;1;1;1;1;1;1;1;1;0.997343 |
| c-1845_conseq_Otu00306_4_1588_bp | 96.5 | Eukaryota;Opisthokonta;Nucletmycea;Fungi;Chytridiomycota;Incertae_Sedis;Chytridiomycetes | 0.99999;0.99999;0.99999;0.99999;0.99999;0.99999;0.99999 |
| c-1889_conseq_Otu00342_4_1588_bp | 98.2 | Eukaryota;Archaeplastida;Chloroplastida;Chlorophyta;Chlorophyceae;Sphaeropleales;Bracteacoccus | 0.98397;0.98397;0.98397;0.98397;0.98397;0.98397;0.98397 |
| c-1913_conseq_Otu00043_53_1588_bp | 99.8 | Eukaryota;Archaeplastida;Chloroplastida;Chlorophyta;Trebouxiophyceae | 1;1;1;1;1 |
| c-1973_conseq_Otu00047_46_1588_bp | 98.9 | Eukaryota;SAR;Rhizaria;Cercozoa;Phytomyxea;Spongospora;subterranea | 1;1;1;1;1;0.997535;0.997535 |
| c-1977_conseq_Otu00006_773_1588_bp | 99 | Eukaryota;Opisthokonta;Nucletmycea;Fungi;Mucoromycota;Mortierellomycotina;Incertae_Sedis;Mortierellales;Mortierellaceae;Mortierella | 0.999991;0.999991;0.999991;0.999991;0.999991;0.999991;0.999991;0.999991;0.999991;0.994564 |
| c-2001_conseq_Otu00239_7_1588_bp | 88.4 | Eukaryota;Opisthokonta;Nucletmycea;Fungi;Cryptomycota | 0.999998;0.999998;0.999998;0.999998;0.999998 |
| c-2016_conseq_Otu00099_20_1588_bp | 87.5 | Eukaryota;Opisthokonta;Nucletmycea;Fungi;Cryptomycota | 0.999994;0.999994;0.999994;0.999994;0.999994 |
| c-204_conseq_Otu00305_4_1588_bp | 91.9 | Eukaryota;Opisthokonta;Nucletmycea;Fungi;Chytridiomycota;Incertae_Sedis;Chytridiomycetes;Cladochytriales;Nowakowskiellaceae;Nowakowskiella | 1;1;1;1;1;1;1;1;1;1 |
| c-2056_conseq_Otu00208_8_1588_bp | 89 | Eukaryota;Opisthokonta;Nucletmycea;Fungi;Cryptomycota | 0.999764;0.999764;0.999764;0.999764;0.999764 |
| c-2077_conseq_Otu00343_4_1588_bp | 97.3 | Eukaryota;Opisthokonta;Nucletmycea;Fungi;Cryptomycota;LKM11 | 1;1;1;1;1;1 |
| c-2101_conseq_Otu00016_219_1588_bp | 98.6 | Eukaryota;Opisthokonta;Nucletmycea;Fungi;Dikarya;Ascomycota;Pezizomycotina;Sordariomycetes;Microascales;Microascaceae | 0.994589;0.994589;0.994589;0.994589;0.994589;0.994589;0.994589;0.994589;0.994589;0.994589 |
| c-2109_conseq_Otu00045_50_1588_bp | 76 | Eukaryota;SAR;Alveolata;Apicomplexa | 0.996007;0.996007;0.996007;0.996007 |
| c-2129_conseq_Otu00467_3_1588_bp | 95.7 | Eukaryota;Opisthokonta;Nucletmycea;Fungi;Chytridiomycota;Incertae_Sedis;Chytridiomycetes | 0.998003;0.998003;0.998003;0.998003;0.998003;0.998003;0.998003 |
| c-2182_conseq_Otu00027_105_1588_bp | 99.9 | Eukaryota;Opisthokonta;Nucletmycea;Fungi;Dikarya;Ascomycota;Pezizomycotina;Dothideomycetes;Capnodiales;Cladosporiaceae;Cladosporium | 0.999998;0.999998;0.999998;0.999998;0.999998;0.999998;0.999998;0.999998;0.999998;0.999998;0.998454 |
| c-2213_conseq_Otu00048_46_1588_bp | 96.8 | Eukaryota;SAR;Rhizaria;Cercozoa;Phytomyxea;Polymyxa;graminis | 1;1;1;1;1;0.936857;0.936857 |
| c-224_conseq_Otu00234_7_1588_bp | 95.5 | Eukaryota;SAR;Rhizaria;Cercozoa;Glissomonadida;Heteromita | 1;1;1;1;1;1 |
| c-2251_conseq_Otu00085_23_1588_bp | 97.4 | Eukaryota;SAR;Stramenopiles;Ochrophyta;Diatomea;Bacillariophytina;Bacillariophyceae;Hantzschia;amphioxys | 1;1;1;1;1;1;1;0.749439;0.749439 |
| c-2326_conseq_Otu00024_107_1588_bp | 99.7 | Eukaryota;SAR;Rhizaria;Cercozoa;Phytomyxea;Polymyxa;graminis | 1;1;1;1;1;1;1 |
| c-2351_conseq_Otu00102_19_1588_bp | 98.8 | Eukaryota;Opisthokonta;Nucletmycea;Fungi;Cryptomycota;LKM11 | 1;1;1;1;1;1 |
| c-2359_conseq_Otu00275_5_1588_bp | 93.2 | Eukaryota;Opisthokonta;Holozoa;Metazoa;Animalia;Eumetazoa;Bilateria;Arthropoda;Chelicerata;Arachnida;Acari | 1;1;1;1;1;1;1;0.990312;0.990312;0.990312;0.990312 |
| c-2360_conseq_Otu00089_22_1588_bp | 97 | Eukaryota;SAR;Stramenopiles;MAST_12;MAST_12C | 0.999984;0.999984;0.999984;0.999984;0.999984 |
| c-2372_conseq_Otu00344_4_1588_bp | 97.4 | Eukaryota;Opisthokonta;Nucletmycea;Fungi;Dikarya;Basidiomycota;Ustilaginomycotina;Exobasidiomycetes;Entylomatales;Incertae_Sedis;Tilletiopsis;Golubevia;pallescens | 1;1;1;1;1;1;1;1;0.566518;0.566518;0.566518;0.566518;0.566518 |
| c-2373_conseq_Otu00070_28_1588_bp | 93.6 | Eukaryota;Opisthokonta;Nucletmycea;Fungi;Cryptomycota;Incertae_Sedis;Incertae_Sedis;Incertae_Sedis;Incertae_Sedis;Rozella | 0.999636;0.999636;0.999636;0.946598;0.946598;0.946598;0.946598;0.946598;0.946598;0.946598 |
| c-2395_conseq_Otu00010_402_1588_bp | 99.7 | Eukaryota;SAR;Rhizaria;Cercozoa;Phytomyxea;Spongospora;subterranea | 1;1;1;1;1;0.932033;0.932033 |
| c-2415_conseq_Otu00284_5_1588_bp | 81.3 | Eukaryota;Amoebozoa;Tubulinea;Euamoebida;Saccamoeba;limax | 1;1;1;1;0.633482;0.633482 |
| c-2420_conseq_Otu00132_14_1588_bp | 96.8 | Eukaryota;Opisthokonta;Nucletmycea;Fungi;Mucoromycota;Mucoromycotina;Incertae_Sedis;Umbelopsidales;Umbelopsidaceae;Umbelopsis | 1;1;1;1;1;1;1;0.924081;0.924081;0.924081 |
| c-2505_conseq_Otu00046_47_1588_bp | 99.1 | Eukaryota;Opisthokonta;Nucletmycea;Fungi;Dikarya;Basidiomycota;Agaricomycotina;Agaricomycetes;Agaricales | 0.999995;0.999995;0.999995;0.999995;0.999995;0.999995;0.999995;0.999995;0.999995 |
| c-2520_conseq_Otu00001_6416_1588_bp | 99.5 | Eukaryota;Archaeplastida;Chloroplastida;Charophyta;Phragmoplastophyta;Streptophyta;Embryophyta;Tracheophyta;Spermatophyta;Magnoliophyta;Brassicales;Brassica | 0.999932;0.999932;0.999932;0.999932;0.999932;0.999932;0.999932;0.999932;0.999932;0.999932;0.999932;0.996383 |
| c-257_conseq_Otu00131_14_1588_bp | 98.3 | Eukaryota;Opisthokonta;Holozoa;Metazoa;Animalia;Eumetazoa;Bilateria;Annelida;Clitellata;Oligochaeta;Haplotaxida;Chamaedrilus;cognettii | 1;1;1;1;1;1;1;1;1;1;1;0.999801;0.927231 |
| c-2629_conseq_Otu00229_7_1588_bp | 98 | Eukaryota;Opisthokonta;Nucletmycea;Fungi;Dikarya;Basidiomycota;Agaricomycotina;Agaricomycetes;Agaricales | 0.999392;0.999392;0.999392;0.999392;0.999392;0.999392;0.999392;0.999392;0.999392 |
| c-2751_conseq_Otu00168_11_1588_bp | 92.9 | Eukaryota;SAR;Rhizaria;Cercozoa;Phytomyxea | 0.999994;0.999994;0.999994;0.999994;0.999994 |
| c-2785_conseq_Otu00114_16_1588_bp | 99.1 | Eukaryota;Archaeplastida;Chloroplastida;Chlorophyta;Chlorophyceae;Chlorosarcinales;Chlorosarcinopsis;delicata | 1;1;1;1;1;1;0.993497;0.993497 |
| c-2786_conseq_Otu00268_6_1588_bp | 95.4 | Eukaryota;Opisthokonta;Nucletmycea;Fungi;Chytridiomycota;Incertae_Sedis;Chytridiomycetes | 0.999221;0.999221;0.999221;0.999221;0.999221;0.999221;0.999221 |
| c-281_conseq_Otu00242_6_1588_bp | 96.1 | Eukaryota;SAR;Stramenopiles;Peronosporomycetes | 0.966192;0.966192;0.966192;0.966192 |
| c-2896_conseq_Otu00261_6_1588_bp | 92.4 | Eukaryota;SAR;Rhizaria;Cercozoa;Phytomyxea | 0.989856;0.989856;0.989856;0.989856;0.989856 |
| c-307_conseq_Otu00241_6_1588_bp | 84.5 | Eukaryota;Amoebozoa | 0.99985;0.99985 |
| c-3206_conseq_Otu00098_20_1588_bp | 99.4 | Eukaryota;Archaeplastida;Chloroplastida;Charophyta;Phragmoplastophyta;Streptophyta;Embryophyta;Tracheophyta;Spermatophyta;Magnoliophyta;Liliopsida;Poales | 1;1;1;1;1;1;1;1;1;1;1;0.9999 |
| c-3464_conseq_Otu00310_4_1588_bp | 95.6 | Eukaryota;Opisthokonta;Nucletmycea;Fungi;Dikarya;Basidiomycota;Agaricomycotina;Agaricomycetes;Corticiales;Corticiaceae | 0.999994;0.999994;0.999994;0.999994;0.999994;0.999994;0.999994;0.999994;0.999994;0.999994 |
| c-3514_conseq_Otu00345_4_1588_bp | 96.7 | Eukaryota;Archaeplastida;Chloroplastida;Chlorophyta;Chlorophyceae | 0.999993;0.999993;0.999993;0.999993;0.999993 |
| c-366_conseq_Otu00300_5_1588_bp | 99.3 | Eukaryota;SAR;Rhizaria;Cercozoa;Thecofilosea | 1;1;1;1;1 |
| c-3676_conseq_Otu00196_9_1588_bp | 95.6 | Eukaryota;SAR;Stramenopiles;MAST_12;MAST_12C | 1;1;1;1;0.709355 |
| c-372_conseq_Otu00117_16_1588_bp | 97.4 | Eukaryota;Opisthokonta;Holozoa;Metazoa;Animalia;Eumetazoa;Bilateria;Nematoda;Enoplea;Enoplia;Enoplida | 0.999743;0.999743;0.999743;0.999743;0.999743;0.999743;0.999743;0.999743;0.999743;0.999743;0.999743 |
| c-3731_conseq_Otu00301_5_1588_bp | 95.7 | Eukaryota;SAR;Rhizaria;Cercozoa;Phytomyxea;Polymyxa;graminis | 1;1;1;1;1;0.87656;0.87656 |
| c-3908_conseq_Otu00293_5_1588_bp | 99.7 | Eukaryota;Archaeplastida;Chloroplastida;Chlorophyta;Chlorophyceae;Actinochloris;BCP_LG3VF20 | 1;1;1;1;1;0.994305;0.994305 |
| c-3956_conseq_Otu00134_13_1588_bp | 99.8 | Eukaryota;Archaeplastida;Chloroplastida;Chlorophyta;Chlorophyceae;Chlamydomonadales | 0.999999;0.999999;0.999999;0.999999;0.999999;0.999999 |
| c-3965_conseq_Otu00331_4_1588_bp | 90.4 | Eukaryota;Opisthokonta;Nucletmycea;Fungi;LKM15 | 0.999756;0.999756;0.999756;0.999756;0.538436 |
| c-404_conseq_Otu00068_30_1588_bp | 98.7 | Eukaryota;SAR;Stramenopiles;Peronosporomycetes;Phytophthora | 1;1;1;1;1 |
| c-410_conseq_Otu00084_23_1588_bp | 99 | Eukaryota;Opisthokonta;Holozoa;Metazoa;Animalia;Eumetazoa;Bilateria;Annelida;Clitellata;Oligochaeta;Haplotaxida;Enchytraeus | 0.999616;0.999616;0.999616;0.999616;0.999616;0.999616;0.999616;0.999616;0.999616;0.999616;0.999616;0.999616 |
| c-4180_conseq_Otu00042_53_1588_bp | 94 | Eukaryota;SAR;Rhizaria;Cercozoa;Phytomyxea | 0.998957;0.998957;0.998957;0.998957;0.998957 |
| c-423_conseq_Otu00155_11_1588_bp | 98.5 | Eukaryota;SAR;Rhizaria;Cercozoa;Cercomonadidae;Cercomonas;edax | 1;1;1;1;1;0.998911;0.998911 |
| c-431_conseq_Otu00124_15_1588_bp | 90.3 | Eukaryota;Opisthokonta;Holozoa;Metazoa;Animalia;Eumetazoa;Bilateria;Nematoda;Chromadorea;Monhysterida | 0.999999;0.999999;0.999999;0.999999;0.999999;0.999999;0.999999;0.999999;0.999999;0.999999 |
| c-4331_conseq_Otu00056_35_1588_bp | 96.1 | Eukaryota;Opisthokonta;Nucletmycea;Fungi;Dikarya;Basidiomycota;Agaricomycotina;Agaricomycetes | 0.993537;0.993537;0.993537;0.993537;0.993537;0.993537;0.993537;0.993537 |
| c-4430_conseq_Otu00468_3_1588_bp | 99.1 | Eukaryota;Archaeplastida;Chloroplastida;Chlorophyta;Chlorophyceae | 0.999973;0.999973;0.999973;0.999973;0.999973 |
| c-4551_conseq_Otu00204_8_1588_bp | 97 | Eukaryota;Opisthokonta;Nucletmycea;Fungi;Chytridiomycota;Incertae_Sedis;Chytridiomycetes | 1;1;1;1;1;1;1 |
| c-4567_conseq_Otu00030_90_1588_bp | 98.9 | Eukaryota;SAR;Stramenopiles;Ochrophyta;Chrysophyceae;Chromulinales;Spumella | 0.99949;0.99949;0.99949;0.99949;0.99949;0.99949;0.99949 |
| c-463_conseq_Otu00236_7_1588_bp | 89.8 | Eukaryota;Amoebozoa;Tubulinea;Arcellinida;Echinamoebida | 0.989184;0.989184;0.989184;0.989184;0.912573 |
| c-4670_conseq_Otu00008_588_1588_bp | 99.8 | Eukaryota;Opisthokonta;Nucletmycea;Fungi;Dikarya;Basidiomycota;Agaricomycotina;Tremellomycetes;Filobasidiales;Piskurozymaceae;Solicoccozyma | 1;1;1;1;1;1;1;1;0.999157;0.999157;0.999157 |
| c-4706_conseq_Otu00213_8_1588_bp | 93.9 | Eukaryota;Opisthokonta;Nucletmycea;Fungi;Chytridiomycota;Incertae_Sedis;Chytridiomycetes | 1;1;1;1;1;1;1 |
| c-474_conseq_Otu00105_18_1588_bp | 99.5 | Eukaryota;SAR;Stramenopiles;Peronosporomycetes | 0.776288;0.776288;0.776288;0.776288 |
| c-4816_conseq_Otu00181_9_1588_bp | 92.8 | Eukaryota;Opisthokonta;Aphelidea | 0.992722;0.992722;0.992722 |
| c-4855_conseq_Otu00231_7_1588_bp | 97.8 | Eukaryota;Archaeplastida;Chloroplastida;Chlorophyta;Trebouxiophyceae;Incertae_Sedis;Parietochloris;alveolaris | 1;1;1;1;1;0.996248;0.996248;0.996248 |
| c-4893_conseq_Otu00115_16_1588_bp | 93.7 | Eukaryota;Opisthokonta;Nucletmycea;Fungi;Dikarya;Basidiomycota;Agaricomycotina;Agaricomycetes;Agaricales;Tricholomataceae;Camarophyllopsis | 0.995591;0.995591;0.995591;0.995591;0.995591;0.995591;0.995591;0.995591;0.995591;0.995591;0.572003 |
| c-4930_conseq_Otu00060_33_1588_bp | 99.8 | Eukaryota;Archaeplastida;Chloroplastida;Chlorophyta;Chlorophyceae;Chlamydomonadales;Chlorococcum | 1;1;1;1;1;0.998726;0.998726 |
| c-5093_conseq_Otu00038_64_1588_bp | 96.7 | Eukaryota;SAR;Alveolata;Ciliophora;Intramacronucleata;Conthreep;Colpodea;Colpodida | 0.988859;0.988859;0.988859;0.988859;0.988859;0.988859;0.988859;0.988082 |
| c-5136_conseq_Otu00022_129_1588_bp | 96.4 | Eukaryota;Opisthokonta;Nucletmycea;Fungi;Dikarya;Basidiomycota;Agaricomycotina;Agaricomycetes;Hymenochaetales;Basidioascus;CBS12316 | 1;1;1;1;1;1;1;0.995704;0.995704;0.995704;0.995704 |
| c-5148_conseq_Otu00082_24_1588_bp | 93 | Eukaryota;Opisthokonta;Nucletmycea;Fungi;Cryptomycota | 1;1;1;1;1 |
| c-5150_conseq_Otu00466_3_1588_bp | 99.2 | Eukaryota;Opisthokonta;Holozoa;Metazoa;Animalia;Eumetazoa;Bilateria;Arthropoda;Chelicerata;Arachnida;Acari | 1;1;1;1;1;1;1;1;1;1;1 |
| c-5167_conseq_Otu00123_15_1588_bp | 94.3 | Eukaryota;Opisthokonta;Nucletmycea;Fungi;Chytridiomycota;Incertae_Sedis;Chytridiomycetes | 1;1;1;1;1;1;1 |
| c-5184_conseq_Otu00111_17_1588_bp | 99.8 | Eukaryota;Opisthokonta;Nucletmycea;Fungi;Dikarya;Basidiomycota;Agaricomycotina;Tremellomycetes;Cystofilobasidiales;Mrakiaceae;Itersonilia | 1;1;1;1;1;1;1;1;1;1;0.996403 |
| c-520_conseq_Otu00461_3_1588_bp | 98.6 | Eukaryota;Amoebozoa;Schizoplasmodiida;Ceratiomyxella;tahitiensis | 1;1;0.873752;0.873752;0.873752 |
| c-5233_conseq_Otu00219_8_1588_bp | 99.9 | Eukaryota;Archaeplastida;Chloroplastida;Chlorophyta;Chlorophyceae;Chlamydomonadales;Tetracystis;sarcinalis | 1;1;1;1;1;1;0.988359;0.988359 |
| c-5240_conseq_Otu00164_11_1588_bp | 95.7 | Eukaryota;SAR;Alveolata;Ciliophora;Intramacronucleata;Conthreep;Colpodea;Platyophryida | 1;1;1;1;1;1;1;0.732511 |
| c-5292_conseq_Otu00119_16_1588_bp | 98.8 | Eukaryota;Opisthokonta;Nucletmycea;Fungi;Dikarya;Basidiomycota;Agaricomycotina;Agaricomycetes | 1;1;1;1;1;1;1;1 |

| query | % similarity | annotation | confidence for each rank |
|---|---|---|---|
| c-5298_conseq_Otu00145_12_1588_bp | 97.4 | Eukaryota;SAR;Stramenopiles;Ochrophyta;Chrysophyceae;Chromulinales | 1;1;1;1;1;1 |
| c-5323_conseq_Otu00091_22_1588_bp | 99.8 | Eukaryota;Opisthokonta;Nucletmycea;Fungi;Dikarya;Basidiomycota;Agaricomycotina;Tremellomycetes;Cystofilobasidiales;Cystofilobasidiaceae;Cystofilobasidium | 0.777046;0.777046;0.777046;0.777046;0.777046;0.777046;0.777046;0.777046;0.777046;0.776634;0.776634 |
| c-5380_conseq_Otu00066_31_1588_bp | 99.8 | Eukaryota;Archaeplastida;Chloroplastida;Chlorophyta;Chlorophyceae;Chlamydomonadales;Tetracystis | 1;1;1;1;0.999179;0.61441;0.61441 |
| c-5383_conseq_Otu00097_20_1588_bp | 98.6 | Eukaryota;Opisthokonta;Nucletmycea;Fungi;Dikarya;Basidiomycota;Pucciniomycotina;Microbotryomycetes | 1;1;1;1;1;1;1;0.9884 |
| c-5450_conseq_Otu00463_3_1588_bp | 82.6 | Eukaryota;SAR;Stramenopiles;Labyrinthulomycetes;Sorodiplophrys | 0.998702;0.998702;0.998702;0.998702;0.998702 |
| c-5485_conseq_Otu00128_14_1588_bp | 91.6 | Eukaryota;Opisthokonta;Nucletmycea;Fungi;Cryptomycota | 0.991707;0.991707;0.991707;0.991707;0.991707 |
| c-551_conseq_Otu00151_12_1588_bp | 84.5 | Eukaryota;Opisthokonta;Holozoa;Metazoa;Animalia;Eumetazoa;Bilateria;Nematoda;Enoplea;Dorylaimia | 1;1;1;1;1;1;1;1;1;1 |
| c-557_conseq_Otu00113_17_1588_bp | 94.9 | Eukaryota;Opisthokonta;Holozoa;Metazoa;Animalia;Eumetazoa;Bilateria;Arthropoda;Hexapoda;Ellipura;Collembola | 1;1;1;1;1;1;1;1;0.917082;0.917082;0.917082 |
| c-5661_conseq_Otu00465_3_1588_bp | 95.5 | Eukaryota;Opisthokonta;Nucletmycea;Fungi;Dikarya;Basidiomycota | 0.999998;0.999998;0.999998;0.999998;0.999998;0.999998 |
| c-5721_conseq_Otu00281_5_1588_bp | 98.1 | Eukaryota;Opisthokonta;Nucletmycea;Fungi;Chytridiomycota;Incertae_Sedis;Chytridiomycetes;Rhizophydiales | 0.999707;0.999707;0.999707;0.999707;0.999707;0.999707;0.999707;0.993763 |
| c-5731_conseq_Otu00235_7_1588_bp | 98.6 | Eukaryota;Opisthokonta;Nucletmycea;Fungi;Chytridiomycota;Incertae_Sedis;Chytridiomycetes;Rhizophydiales | 1;1;1;1;1;1;1;1 |
| c-5778_conseq_Otu00080_24_1588_bp | 96.1 | Eukaryota;Opisthokonta;Nucletmycea;Fungi;Dikarya;Ascomycota;Pezizomycotina;Eurotiomycetes;Onygenales;Gymnoascaceae;Leucothecium;emdenii | 1;1;1;1;1;1;1;1;1;0.99854;0.99854;0.99854 |
| c-5788_conseq_Otu00137_13_1588_bp | 88.6 | Eukaryota;Opisthokonta;Nucletmycea;Fungi;Zoopagomycota | 0.985211;0.670244;0.670244;0.670244;0.670244 |
| c-5829_conseq_Otu00224_7_1588_bp | 90.2 | Eukaryota;Opisthokonta;Nucletmycea;Fungi;Dikarya;Basidiomycota;Pucciniomycotina | 0.998929;0.998929;0.998929;0.998929;0.998929;0.998929;0.998929 |
| c-5833_conseq_Otu00040_54_1588_bp | 95.6 | Eukaryota;SAR;Alveolata;Ciliophora;Intramacronucleata;Conthreep;Colpodea;Colpodida | 0.995051;0.995051;0.995051;0.995051;0.995051;0.995051;0.995051;0.995051 |
| c-5853_conseq_Otu00272_6_1588_bp | 88.7 | Eukaryota;Opisthokonta;Nucletmycea;Fungi;Cryptomycota | 0.99126;0.99126;0.99126;0.99126;0.99126 |
| c-589_conseq_Otu00057_34_1588_bp | 97.8 | Eukaryota;Opisthokonta;Holozoa;Metazoa;Animalia;Eumetazoa;Bilateria;Nematoda;Chromadorea;Tylenchida | 0.99994;0.99994;0.99994;0.99994;0.99994;0.99994;0.99994;0.99994;0.99994;0.99994 |
| c-5912_conseq_Otu00176_10_1588_bp | 96.9 | Eukaryota;SAR;Alveolata;Ciliophora;Intramacronucleata;Conthreep;Colpodea | 1;1;1;1;1;1;1 |
| c-5922_conseq_Otu00173_10_1588_bp | 98.2 | Eukaryota;Opisthokonta;Nucletmycea;Fungi;Dikarya;Basidiomycota;Agaricomycotina;Agaricomycetes;Auriculariales | 0.999999;0.999999;0.999999;0.999999;0.999999;0.999999;0.999999;0.999999;0.999999 |
| c-5947_conseq_Otu00223_7_1588_bp | 99.6 | Eukaryota;Opisthokonta;Nucletmycea;Fungi;Dikarya;Basidiomycota;Pucciniomycotina;Cystobasidiomycetes;Cystobasidiales;Incertae_Sedis;Occultifur;externus | 0.999622;0.999622;0.999622;0.999622;0.999622;0.999622;0.999622;0.999622;0.999622;0.992919;0.992919;0.992919 |
| c-5948_conseq_Otu00218_8_1588_bp | 99.4 | Eukaryota;Opisthokonta;Nucletmycea;Fungi;Dikarya;Ascomycota;Pezizomycotina;Dothideomycetes;Pleosporales;Pleosporaceae;Pleospora;herbarum | 1;1;1;1;1;1;1;1;1;1;0.992206;0.992206 |
| c-5959_conseq_Otu00122_15_1588_bp | 97.9 | Eukaryota;Opisthokonta;Nucletmycea;Fungi;Dikarya;Ascomycota;Pezizomycotina;Laboulbeniomycetes | 1;1;1;1;1;1;1;0.804799 |
| c-5983_conseq_Otu00007_740_1588_bp | 98.9 | Eukaryota;Opisthokonta;Nucletmycea;Fungi;Dikarya;Ascomycota;Pezizomycotina;Sordariomycetes;Glomerellales;Plectosphaerellaceae;Plectosphaerella | 0.999999;0.999999;0.999999;0.999999;0.999999;0.999999;0.999999;0.999999;0.999999;0.999672;0.999672 |
| c-5997_conseq_Otu00062_31_1588_bp | 99 | Eukaryota;Opisthokonta;Nucletmycea;Fungi;Dikarya;Ascomycota;Pezizomycotina;Pezizomycetes;Pezizales;Rhizinaceae;Phymatotrichopsis | 0.992588;0.992588;0.992588;0.992588;0.992588;0.992588;0.992588;0.992588;0.992588;0.992551;0.992551 |
| c-6000_conseq_Otu00035_73_1588_bp | 98 | Eukaryota;Opisthokonta;Nucletmycea;Fungi;Dikarya;Ascomycota;Pezizomycotina;Sordariomycetes;Coniochaetales;Coniochaetaceae;Coniochaeta | 0.999999;0.999999;0.999999;0.999999;0.999999;0.999999;0.999999;0.999999;0.996781;0.996781;0.996781 |
| c-6005_conseq_Otu00020_161_1588_bp | 98.6 | Eukaryota;Opisthokonta;Nucletmycea;Fungi;Dikarya;Ascomycota;Pezizomycotina;Dothideomycetes;Pleosporales | 0.996779;0.996779;0.996779;0.996779;0.996779;0.996779;0.996779;0.996779;0.996779 |
| c-6009_conseq_Otu00460_3_1588_bp | 96.5 | Eukaryota;Opisthokonta;Nucletmycea;Fungi;Chytridiomycota;Incertae_Sedis;Chytridiomycetes;Rhizophydiales | 0.99922;0.99922;0.99922;0.99922;0.99922;0.99922;0.99922;0.99922 |
| c-6067_conseq_Otu00186_9_1588_bp | 93 | Eukaryota;SAR;Stramenopiles;Ochrophyta;Chrysophyceae | 0.510247;0.510247;0.510247;0.510247;0.510247 |
| c-6131_conseq_Otu00018_189_1588_bp | 87.3 | Eukaryota;SAR;Alveolata;Apicomplexa;Conoidasida;Gregarinasina | 0.999917;0.999917;0.999917;0.999917;0.999917;0.999917 |
| c-6137_conseq_Otu00067_31_1588_bp | 99.5 | Eukaryota;Opisthokonta;Nucletmycea;Fungi;Dikarya;Ascomycota;Pezizomycotina;Sordariomycetes;Xylariales;Apiosporaceae;Arthrinium | 1;1;1;1;1;1;1;1;1;0.999484;0.999484 |
| c-6143_conseq_Otu00240_6_1588_bp | 86.8 | Eukaryota;SAR;Alveolata;Apicomplexa;Conoidasida;Gregarinasina;Eugregarinorida;Syncystis;mirabilis | 1;1;1;1;1;1;0.703347;0.703347;0.703347 |
| c-6164_conseq_Otu00017_209_1588_bp | 98.2 | Eukaryota;Opisthokonta;Nucletmycea;Fungi;Dikarya;Ascomycota;Saccharomycotina;Saccharomycetes;Saccharomycetales;Incertae_Sedis;Nadsonia | 0.999942;0.999942;0.999942;0.999942;0.999942;0.999942;0.999942;0.999942;0.999942;0.999942;0.999942 |
| c-6170_conseq_Otu00108_18_1588_bp | 89.6 | Eukaryota;Opisthokonta;Nucletmycea;Fungi | 0.961847;0.961847;0.961847;0.961847 |
| c-6173_conseq_Otu00266_6_1588_bp | 97.2 | Eukaryota;SAR;Alveolata;Ciliophora;Intramacronucleata;Spirotrichea;Hypotrichia | 0.999818;0.999818;0.999818;0.999818;0.999818;0.999818;0.999818 |
| c-6224_conseq_Otu00029_92_1588_bp | 98.6 | Eukaryota;SAR;Alveolata;Ciliophora;Intramacronucleata;Spirotrichea;Hypotrichia | 0.999999;0.999999;0.999999;0.999999;0.999999;0.999999;0.999999 |
| c-6233_conseq_Otu00072_27_1588_bp | 98.5 | Eukaryota;Opisthokonta;Nucletmycea;Fungi;Dikarya;Ascomycota;Pezizomycotina;Pezizomycetes;Pezizales | 0.999804;0.999804;0.999804;0.999804;0.999804;0.999804;0.999804;0.999804;0.999804 |
| c-6244_conseq_Otu00271_6_1588_bp | 93.6 | Eukaryota;Opisthokonta;Nucletmycea;Fungi;Chytridiomycota;Incertae_Sedis;Chytridiomycetes;Rhizophydiales | 0.992453;0.992453;0.992453;0.992453;0.992453;0.992453;0.992453;0.992453 |
| c-6309_conseq_Otu00036_69_1588_bp | 99.1 | Eukaryota;SAR;Alveolata;Ciliophora;Intramacronucleata;Spirotrichea;Hypotrichia;uncultured;Oxytrich | 1;1;1;1;1;1;1;0.993314;0.993314 |
| c-6324_conseq_Otu00225_7_1588_bp | 98.9 | Eukaryota;SAR;Alveolata;Ciliophora;Intramacronucleata;Spirotrichea;Hypotrichia;Urostyla | 1;1;1;1;1;1;1;0.95621 |
| c-6332_conseq_Otu00025_107_1588_bp | 97.7 | Eukaryota;Opisthokonta;Nucletmycea;Fungi;Dikarya;Ascomycota;Pezizomycotina;Pezizomycetes;Pezizales;Pyronemataceae | 0.985336;0.985336;0.985336;0.985336;0.985336;0.985336;0.985336;0.985336;0.985336;0.985336 |
| c-6351_conseq_Otu00162_11_1588_bp | 99.5 | Eukaryota;SAR;Alveolata;Ciliophora;Intramacronucleata;Spirotrichea;Hypotrichia;Engelmanniella;mobilis | 0.990607;0.990607;0.990607;0.990607;0.990607;0.990607;0.990607;0.990607;0.990607 |
| c-6398_conseq_Otu00109_17_1588_bp | 98.1 | Eukaryota;Opisthokonta;Nucletmycea;Fungi;Dikarya;Ascomycota;Pezizomycotina;Dothideomycetes;Pleosporales | 0.920196;0.920196;0.920196;0.920196;0.920196;0.920196;0.920196;0.920196;0.920196 |
| c-6410_conseq_Otu00159_11_1588_bp | 91.9 | Eukaryota;SAR;Alveolata;Apicomplexa;Conoidasida;Cryptosporida;Cryptosporidium;Colpodella | 0.999907;0.999907;0.999907;0.999907;0.999907;0.999907;0.999907;0.999907 |
| c-6421_conseq_Otu00004_935_1588_bp | 99.8 | Eukaryota;Opisthokonta;Nucletmycea;Fungi;Dikarya;Ascomycota;Pezizomycotina;Leotiomycetes | 0.999999;0.999999;0.999999;0.999999;0.999999;0.999999;0.999999;0.999999 |
| c-6445_conseq_Otu00282_5_1588_bp | 96.7 | Eukaryota;SAR;Alveolata;Ciliophora;Intramacronucleata;Spirotrichea;Hypotrichia | 0.947552;0.947552;0.947552;0.947552;0.947552;0.947552;0.947552 |
| c-6454_conseq_Otu00031_81_1588_bp | 99.1 | Eukaryota;Opisthokonta;Nucletmycea;Fungi;Dikarya;Ascomycota;Pezizomycotina;Leotiomycetes;Helotiales;Incertae_Sedis;Tetracladium | 1;1;1;1;1;1;1;1;0.999377;0.999377;0.999377 |
| c-6456_conseq_Otu00059_33_1588_bp | 99.3 | Eukaryota;SAR;Alveolata;Ciliophora;Intramacronucleata;Spirotrichea;Hypotrichia | 0.980626;0.980626;0.980626;0.980626;0.980626;0.980626;0.980626 |
| c-6471_conseq_Otu00232_7_1588_bp | 95.8 | Eukaryota;Opisthokonta;Nucletmycea;Fungi;Dikarya;Ascomycota;Pezizomycotina;Dothideomycetes;Incertae_Sedis | 1;1;1;1;1;1;1;1;1 |
| c-648_conseq_Otu00346_4_1588_bp | 96.9 | Eukaryota;Archaeplastida;Chloroplastida;Charophyta;Phragmoplastophyta;Zygnematophyceae | 0.998173;0.998173;0.998173;0.998173;0.998173;0.998173 |
| c-6481_conseq_Otu00179_10_1588_bp | 97.3 | Eukaryota;Opisthokonta;Nucletmycea;Fungi;Dikarya;Ascomycota;Pezizomycotina;Dothideomycetes;Tubeufiales;Tubeufiaceae;Tubeufia;cerea | 0.999993;0.999993;0.999993;0.999993;0.999993;0.999993;0.999993;0.999993;0.999993;0.999993;0.63854;0.63854 |
| c-6538_conseq_Otu00051_44_1588_bp | 98.7 | Eukaryota;Opisthokonta;Nucletmycea;Fungi;Dikarya;Basidiomycota;Agaricomycotina;Tremellomycetes;Trichosporonales;Trichosporonaceae;Apiotrichum | 0.999999;0.999999;0.999999;0.999999;0.999999;0.999999;0.999999;0.999999;0.99935;0.99935;0.99935 |
| c-6656_conseq_Otu00015_231_1588_bp | 99.7 | Eukaryota;Opisthokonta;Nucletmycea;Fungi;Dikarya;Ascomycota;Pezizomycotina;Dothideomycetes;Pleosporales | 0.999576;0.999576;0.999576;0.999576;0.999576;0.999576;0.999576;0.999576;0.999576 |
| c-6725_conseq_Otu00152_12_1588_bp | 90.1 | Eukaryota;Opisthokonta;Nucletmycea;Fungi;Dikarya;Ascomycota;Pezizomycotina;Sordariomycetes;Hypocreales | 0.987395;0.987395;0.987395;0.987395;0.987395;0.987395;0.987395;0.987395;0.987395 |
| c-6814_conseq_Otu00005_923_1588_bp | 99.4 | Eukaryota;Opisthokonta;Nucletmycea;Fungi;Dikarya;Ascomycota;Pezizomycotina;Sordariomycetes | 0.992209;0.992209;0.992209;0.992209;0.992209;0.992209;0.992209;0.992209 |
| c-6882_conseq_Otu00157_11_1588_bp | 99.8 | Eukaryota;SAR;Alveolata;Ciliophora;Intramacronucleata;Spirotrichea;Hypotrichia;Hemiurosomoida;longa | 1;1;1;1;1;1;1;0.90841;0.90841 |
| c-6926_conseq_Otu00100_20_1588_bp | 99.2 | Eukaryota;Opisthokonta;Nucletmycea;Fungi;Dikarya;Basidiomycota;Agaricomycotina;Tremellomycetes;Tremellales;Bulleribasidiaceae;Vishniacozyma;victoriae | 1;1;1;1;1;1;1;1;1;1;1;0.988634 |
| c-694_conseq_Otu00197_8_1588_bp | 99.3 | Eukaryota;Opisthokonta;Holozoa;Metazoa;Animalia;Eumetazoa;Bilateria;Nematoda;Chromadorea;Rhabditida | 1;1;1;1;1;1;1;1;1;1 |
| c-6998_conseq_Otu00459_3_1588_bp | 98.5 | Eukaryota;SAR;Alveolata;Ciliophora;Intramacronucleata;Spirotrichea;Hypotrichia | 0.999992;0.999992;0.999992;0.999992;0.999992;0.999992;0.999992 |
| c-7059_conseq_Otu00252_6_1588_bp | 94.7 | Eukaryota;Opisthokonta;Nucletmycea;Fungi;Dikarya;Ascomycota;Pezizomycotina;Leotiomycetes | 0.973213;0.973213;0.973213;0.973213;0.973213;0.973213;0.973213;0.973213 |
| c-7298_conseq_Otu00462_3_1588_bp | 94.1 | Eukaryota;Opisthokonta;Nucletmycea;Fungi;Cryptomycota | 1;1;1;1;1 |
| c-7334_conseq_Otu00052_43_1588_bp | 99.9 | Eukaryota;Opisthokonta;Nucletmycea;Fungi;Dikarya;Ascomycota;Pezizomycotina;Dothideomycetes;Capnodiales;Extremaceae;Extremus;Capnodiales | 0.999999;0.999999;0.999999;0.999999;0.999999;0.999999;0.999999;0.999999;0.999999;0.999879;0.863598;0.863598 |
| c-7374_conseq_Otu00221_7_1588_bp | 99.4 | Eukaryota;Opisthokonta;Nucletmycea;Fungi;Dikarya;Ascomycota;Pezizomycotina;Sordariomycetes;Microascales | 0.999912;0.999912;0.999912;0.999912;0.999912;0.999912;0.999912;0.999912;0.999912 |
| c-743_conseq_Otu00058_33_1588_bp | 99.6 | Eukaryota;Opisthokonta;Holozoa;Metazoa;Animalia;Eumetazoa;Bilateria;Nematoda;Chromadorea;Rhabditida;Chiloplacus;propinquus | 0.999995;0.999995;0.999995;0.999995;0.999995;0.999995;0.999995;0.999995;0.999995;0.999995;0.626114;0.626114 |
| c-7528_conseq_Otu00329_4_1588_bp | 96.4 | Eukaryota;Opisthokonta;Nucletmycea;Fungi;Dikarya;Ascomycota;Pezizomycotina;Pezizomycetes;Pezizales;Ascobolaceae;Thecotheus;holmskjoldii | 1;1;1;1;1;1;1;1;1;0.779091;0.779091;0.779091 |
| c-7548_conseq_Otu00079_24_1588_bp | 98.8 | Eukaryota;Opisthokonta;Nucletmycea;Fungi;Dikarya;Ascomycota;Pezizomycotina;Sordariomycetes;Xylariales | 0.999986;0.999986;0.999986;0.999986;0.999986;0.999986;0.999986;0.999986;0.999986 |
| c-7549_conseq_Otu00148_12_1588_bp | 91.8 | Eukaryota;Opisthokonta;Nucletmycea;Fungi;Cryptomycota | 1;1;1;1;1 |
| c-7611_conseq_Otu00011_390_1588_bp | 99.4 | Eukaryota;Opisthokonta;Nucletmycea;Fungi;Dikarya;Ascomycota;Pezizomycotina;Sordariomycetes;Sordariales | 0.9997;0.9997;0.9997;0.9997;0.9997;0.9997;0.9997;0.9997;0.9997 |
| c-7670_conseq_Otu00012_375_1588_bp | 99.8 | Eukaryota;Opisthokonta;Nucletmycea;Fungi;Dikarya;Basidiomycota;Agaricomycotina;Tremellomycetes;Tremellales;Trimorphomycetaceae | 0.995835;0.995835;0.995835;0.995835;0.995835;0.995835;0.995835;0.995835;0.995835;0.989352 |
| c-7708_conseq_Otu00198_8_1588_bp | 95.6 | Eukaryota;SAR;Alveolata;Ciliophora;Intramacronucleata;Spirotrichea;Hypotrichia;Halteria | 0.969946;0.969946;0.969946;0.969946;0.969946;0.969946;0.969946;0.932725 |
| c-7844_conseq_Otu00350_4_1588_bp | 99.4 | Eukaryota;Opisthokonta;Holozoa;Metazoa;Animalia;Eumetazoa;Bilateria;Nematoda;Chromadorea;Rhabditida;Rhabditophanes;KR3021 | 1;1;1;1;1;1;1;1;1;1;0.99902;0.99902 |
| c-786_conseq_Otu00440_3_1588_bp | 88.9 | Eukaryota;SAR;Stramenopiles;Peronosporomycetes | 0.998775;0.998775;0.998775;0.998775 |
| c-7870_conseq_Otu00294_5_1588_bp | 94.2 | Eukaryota;Opisthokonta;Nucletmycea;Fungi;Cryptomycota;LKM11 | 0.99996;0.99996;0.99996;0.99996;0.99996;0.973215 |
| c-789_conseq_Otu00238_7_1588_bp | 97.2 | Eukaryota;SAR;Rhizaria;Cercozoa;Cercomonadidae;Cercomonas;ambigua | 1;1;1;1;1;0.650704;0.650704 |
| c-7917_conseq_Otu00233_7_1588_bp | 86.1 | Eukaryota;SAR;Alveolata | 0.976453;0.976453;0.976453 |
| c-7974_conseq_Otu00101_19_1588_bp | 99.8 | Eukaryota;Opisthokonta;Nucletmycea;Fungi;Dikarya;Basidiomycota;Agaricomycotina;Tremellomycetes;Tremellales;Bulleribasidiaceae;Dioszegia;crocea | 0.997813;0.997813;0.997813;0.997813;0.997813;0.997813;0.997813;0.997813;0.997813;0.997813;0.997813;0.949887 |
| c-806_conseq_Otu00116_16_1588_bp | 98.7 | Eukaryota;SAR;Rhizaria;Cercozoa;Cercomonadidae | 0.999999;0.999999;0.999999;0.999999;0.999999 |
| c-8229_conseq_Otu00033_76_1588_bp | 89.2 | Eukaryota;SAR;Alveolata;Apicomplexa;Conoidasida;Gregarinasina;Eugregarinorida;Syncystis;mirabilis | 1;1;1;1;1;1;0.673591;0.673591;0.673591 |
| c-8263_conseq_Otu00353_4_1588_bp | 92.3 | Eukaryota;SAR;Alveolata;Apicomplexa;Conoidasida;Gregarinasina | 0.993456;0.993456;0.993456;0.993456;0.993456;0.993456 |
| c-8326_conseq_Otu00210_8_1588_bp | 89.6 | Eukaryota;Opisthokonta;Nucletmycea;Fungi;Cryptomycota | 0.99853;0.99853;0.99853;0.99853;0.99853 |
| c-8342_conseq_Otu00337_4_1588_bp | 84.1 | Eukaryota;SAR;Alveolata | 0.779994;0.772631;0.772631 |
| c-8348_conseq_Otu00325_4_1588_bp | 89.4 | Eukaryota;SAR;Alveolata;Apicomplexa;Conoidasida;Cryptosporida;Cryptosporidium | 1;1;1;1;1;1;1 |
| c-8376_conseq_Otu00088_23_1588_bp | 88.6 | Eukaryota;SAR;Alveolata;Apicomplexa;Conoidasida;Gregarinasina;Eugregarinorida;Syncystis;mirabilis | 1;1;1;1;1;1;0.702318;0.702318;0.702318 |
| c-8387_conseq_Otu00135_13_1588_bp | 97.8 | Eukaryota;SAR;Alveolata;Ciliophora;Intramacronucleata;Conthreep;Oligohymenophorea;Peritrichia | 1;1;1;1;1;1;1;1 |
| c-8415_conseq_Otu00130_14_1588_bp | 96.6 | Eukaryota;SAR;Alveolata;Ciliophora;Intramacronucleata;Conthreep;Oligohymenophorea;Peritrichia | 1;1;1;1;1;1;1;1 |
| c-8421_conseq_Otu00094_21_1588_bp | 77.8 | Eukaryota;SAR;Alveolata;Apicomplexa | |
| c-8433_conseq_Otu00250_6_1588_bp | 77.8 | Eukaryota;SAR;Alveolata;Apicomplexa | |
| c-8446_conseq_Otu00021_157_1588_bp | 93.4 | Eukaryota;SAR;Alveolata;Apicomplexa;Conoidasida;Gregarinasina;Eugregarinorida;Syncystis;mirabilis | 1;1;1;1;1;1;0.621301;0.621301;0.621301 |
| c-8459_conseq_Otu00158_11_1588_bp | 77.7 | Eukaryota;SAR;Alveolata;Apicomplexa | |
| c-8502_conseq_Otu00328_4_1588_bp | 82.7 | Eukaryota;SAR;Alveolata;Apicomplexa;Conoidasida;Gregarinasina | 0.992146;0.992146;0.992146;0.992146;0.992146;0.992146 |
| c-8530_conseq_Otu00249_6_1588_bp | 94 | Eukaryota;SAR;Alveolata;Apicomplexa;Conoidasida;Gregarinasina;Eugregarinorida;Syncystis;mirabilis | 1;1;1;1;1;1;0.603452;0.603452;0.603452 |
| c-8534_conseq_Otu00009_435_1588_bp | 77.7 | Eukaryota;SAR;Alveolata;Apicomplexa | |

| query | % similarity | annotation | confidence for each rank |
|---|---|---|---|
| c-8541_conseq_Otu00127_14_1588_bp | 99.7 | Eukaryota;Opisthokonta;Nucletmycea;Fungi;Dikarya;Ascomycota;Saccharomycotina;Saccharomycetes;Saccharomycetales;Incertae_Sedis;Candida | 1;1;1;1;1;1;1;1;0.999028;0.999028 |
| c-8562_conseq_Otu00260_6_1588_bp | 91.7 | Eukaryota;Opisthokonta;Nucletmycea;Fungi;Cryptomycota | 1;1;1;1;1 |
| c-8582_conseq_Otu00283_5_1588_bp | 77.3 | Eukaryota;SAR;Alveolata;Apicomplexa | |
| c-8622_conseq_Otu00298_5_1588_bp | 85.7 | Eukaryota;SAR;Alveolata;Apicomplexa | 0.609329;0.609329;0.609329;0.547554 |
| c-8663_conseq_Otu00061_31_1588_bp | 85.1 | Eukaryota;SAR;Alveolata;Apicomplexa | 0.795619;0.795619;0.795619;0.54155 |
| c-868_conseq_Otu00034_75_1588_bp | 98.5 | Eukaryota;SAR;Rhizaria;Cercozoa;Glissomonadida | 0.989404;0.989404;0.989404;0.989404;0.989404 |
| c-8684_conseq_Otu00154_11_1588_bp | 93.5 | Eukaryota;SAR;Alveolata;Apicomplexa;Conoidasida;Gregarinasina | 1;1;1;1;1;1 |
| c-8709_conseq_Otu00142_13_1588_bp | 84.2 | Eukaryota;SAR;Alveolata | 0.987795;0.987795;0.987795 |
| c-8714_conseq_Otu00118_16_1588_bp | 88.5 | Eukaryota;Opisthokonta;Nucletmycea;Fungi;Cryptomycota | 0.993767;0.993767;0.993767;0.993767;0.993767 |
| c-8805_conseq_Otu00184_9_1588_bp | 77.3 | Eukaryota;SAR;Rhizaria;Cercozoa;Phytomyxea | 0.998044;0.998044;0.998044;0.998044;0.998044 |
| c-8830_conseq_Otu00303_5_1588_bp | 77.5 | Eukaryota;SAR;Rhizaria;Cercozoa;Phytomyxea | 0.997368;0.997368;0.997368;0.997368;0.997368 |
| c-8839_conseq_Otu00090_22_1588_bp | 82.1 | Eukaryota;SAR;Alveolata;Apicomplexa | |
| c-8846_conseq_Otu00201_8_1588_bp | 81.7 | Eukaryota;SAR;Alveolata;Apicomplexa | 0.575481;0.575481;0.575481;0.56411 |
| c-8874_conseq_Otu00254_6_1588_bp | 81.7 | Eukaryota;SAR;Alveolata;Apicomplexa | 0.96828;0.96828;0.96828;0.962167 |
| c-8891_conseq_Otu00023_117_1588_bp | 99 | Eukaryota;SAR;Alveolata;Ciliophora;Intramacronucleata;Litostomatea;Haptoria;Arcuospathidium | 0.998247;0.998247;0.998247;0.998247;0.998247;0.998247;0.998247;0.998247 |
| c-8898_conseq_Otu00087_23_1588_bp | 98.1 | Eukaryota;SAR;Alveolata;Ciliophora;Intramacronucleata;Litostomatea;Haptoria | 0.997935;0.997935;0.997935;0.997935;0.997935;0.997935;0.997935 |
| c-906_conseq_Otu00146_12_1588_bp | 98.3 | Eukaryota;Opisthokonta;Holozoa;Metazoa;Animalia;Eumetazoa;Bilateria;Nematoda;Chromadorea;Diplogasterida | 0.999904;0.999904;0.999904;0.999904;0.999904;0.999904;0.999904;0.999904;0.999904;0.999904 |
| c-9118_conseq_Otu00019_186_1588_bp | 92.2 | Eukaryota;SAR;Rhizaria;Cercozoa;Phytomyxea | 1;1;1;1;1 |
| c-9230_conseq_Otu00285_5_1588_bp | 75.1 | Eukaryota;SAR;Alveolata;Apicomplexa | |
| c-927_conseq_Otu00054_43_1588_bp | 97.4 | Eukaryota;Opisthokonta;Nucletmycea;Fungi;Dikarya;Basidiomycota;Agaricomycotina;Tremellomycetes;Holtermanniales;Incertae_Sedis;Holtermanniella | 0.998033;0.998033;0.998033;0.998033;0.998033;0.998033;0.998033;0.998033;0.998033;0.998033;0.998033 |
| c-9300_conseq_Otu00112_17_1588_bp | 79.5 | Eukaryota;SAR;Alveolata | 0.982333;0.982333;0.982333 |
| c-9326_conseq_Otu00206_8_1588_bp | 78.9 | Eukaryota;SAR;Alveolata | 0.934749;0.925077;0.925077 |
| c-935_conseq_Otu00069_29_1588_bp | 98.6 | Eukaryota;SAR;Rhizaria;Cercozoa;Glissomonadida;Orciraptor | 1;1;1;1;1;0.751392 |
| c-947_conseq_Otu00141_13_1588_bp | 98.9 | Eukaryota;Opisthokonta;Holozoa;Metazoa;Animalia;Eumetazoa;Bilateria;Nematoda;Chromadorea;Tylenchida;Pratylenchoides | 0.999948;0.999948;0.999948;0.999948;0.999948;0.999948;0.999948;0.999948;0.999948;0.999948;0.994859 |
| c-982_conseq_Otu00332_4_1588_bp | 92.8 | Eukaryota;Opisthokonta;Holozoa;Metazoa;Animalia;Eumetazoa;Bilateria;Arthropoda;Crustacea;Maxillopoda;Copepoda | 0.999341;0.999341;0.999341;0.999341;0.999341;0.999341;0.999341;0.999341;0.9292;0.9292;0.9292 |
| c-9835_conseq_Otu00170_10_1588_bp | 79.7 | Eukaryota;Opisthokonta | 0.989201;0.989201 |
| m54032_170210_154134_11797034_ccs_Otu00575_2_1588_bp | 85.2 | Eukaryota | 0.998158 |
| m54032_170210_154134_12583609_ccs_Otu00323_4_1588_bp | 93.1 | Eukaryota;SAR;Rhizaria;Cercozoa;Vampyrellidae | 0.998761;0.998761;0.998761;0.998761;0.998761 |
| m54032_170210_154134_13239088_ccs_Otu00193_9_1588_bp | 99 | Eukaryota;SAR;Rhizaria;Cercozoa | 0.624512;0.624512;0.624512;0.624512 |
| m54032_170210_154134_14418720_ccs_Otu00358_4_1588_bp | 86.2 | Eukaryota;SAR;Rhizaria;Cercozoa | 0.638659;0.638659;0.638659;0.638659 |
| m54032_170210_154134_14877555_ccs_Otu00563_2_1588_bp | 91.1 | Eukaryota;SAR;Stramenopiles;Bicosoecida | 0.999999;0.999999;0.999999;0.999999 |
| m54032_170210_154134_15401940_ccs_Otu00139_13_1588_bp | 78.4 | Eukaryota;SAR;Alveolata;Apicomplexa | |
| m54032_170210_154134_15795028_ccs_Otu00039_63_1588_bp | 87.9 | Eukaryota;SAR;Alveolata;Ciliophora;Intramacronucleata;Litostomatea;Haptoria | 0.995613;0.995613;0.995613;0.995613;0.995613;0.995613;0.995613 |
| m54032_170210_154134_15925581_ccs_Otu00192_9_1588_bp | 78.6 | Eukaryota;Opisthokonta;Nucletmycea;Fungi;Zoopagomycota | 0.988355;0.98831;0.98831;0.98831;0.98831 |
| m54032_170210_154134_16515714_ccs_Otu00255_6_1588_bp | 99.1 | Eukaryota;SAR;Stramenopiles;Ochrophyta;Chrysophyceae;Chromulinales;Pedospumella | 0.999181;0.999181;0.999181;0.999181;0.999181;0.998851;0.627196 |
| m54032_170210_154134_19137092_ccs_Otu00505_2_1588_bp | 95.6 | Eukaryota;SAR;Alveolata;Ciliophora;Intramacronucleata;Litostomatea;Haptoria | 0.994009;0.994009;0.994009;0.994009;0.994009;0.994009;0.994009 |
| m54032_170210_154134_19398856_ccs_Otu00185_9_1588_bp | 93.4 | Eukaryota;SAR;Rhizaria;Cercozoa | 0.999989;0.999989;0.999989;0.999989 |
| m54032_170210_154134_19595523_ccs_Otu00529_2_1588_bp | 96.3 | Eukaryota;SAR;Stramenopiles;Ochrophyta;Chrysophyceae | 0.999612;0.999612;0.999612;0.999612;0.999612 |
| m54032_170210_154134_21365411_ccs_Otu00592_2_1588_bp | 76.6 | Eukaryota | 0.999748 |
| m54032_170210_154134_21430739_ccs_Otu00335_4_1588_bp | 70.7 | Eukaryota;SAR;Alveolata;Apicomplexa | 0.98349;0.98349;0.98349;0.98349 |
| m54032_170210_154134_23790545_ccs_Otu00578_2_1588_bp | 85.8 | Eukaryota;SAR;Rhizaria;Cercozoa | 0.996087;0.996087;0.996087;0.996087 |
| m54032_170210_154134_24248919_ccs_Otu00565_2_1588_bp | 91.1 | Eukaryota;SAR;Rhizaria;Cercozoa;Imbricatea | 0.999683;0.999683;0.999683;0.999683;0.999683 |
| m54032_170210_154134_24511160_ccs_Otu00531_2_1588_bp | 89.7 | Eukaryota;SAR;Rhizaria;Cercozoa;Phytomyxea | 0.999999;0.999999;0.999999;0.999999;0.999999 |
| m54032_170210_154134_25166565_ccs_Otu00446_3_1588_bp | 94.2 | Eukaryota;SAR;Rhizaria;Cercozoa | 0.999966;0.999966;0.999966;0.999966 |
| m54032_170210_154134_26214518_ccs_Otu00063_31_1588_bp | 90.8 | Eukaryota;SAR;Rhizaria;Cercozoa;Glissomonadida;Heteromita | 0.999999;0.999999;0.999999;0.999999;0.999999;0.996752 |
| m54032_170210_154134_26477122_ccs_Otu00436_3_1588_bp | 98.9 | Eukaryota;Archaeplastida;Chloroplastida;Chlorophyta;Trebouxiophyceae | 1;1;1;1;1 |
| m54032_170210_154134_28443142_ccs_Otu00286_5_1588_bp | 86.8 | Eukaryota;Amoebozoa | 0.998347;0.998347 |
| m54032_170210_154134_28508344_ccs_Otu00631_2_1588_bp | 92.8 | Eukaryota;SAR;Alveolata;Ciliophora;Intramacronucleata;Conthreep;Colpodea | 1;1;1;1;1;1;1 |
| m54032_170210_154134_29098380_ccs_Otu00248_6_1588_bp | 77.3 | Eukaryota;SAR;Alveolata;Apicomplexa | |
| m54032_170210_154134_29360267_ccs_Otu00175_10_1588_bp | 97.4 | Eukaryota;Opisthokonta;Nucletmycea;Fungi;Dikarya;Basidiomycota;Agaricomycotina;Agaricomycetes;Agaricales;Cortinariaceae;Cortinarius | 1;1;1;1;1;1;1;1;1;1;1 |
| m54032_170210_154134_29622949_ccs_Otu00299_5_1588_bp | 93.6 | Eukaryota;Opisthokonta;Holozoa;Metazoa;Animalia;Eumetazoa;Bilateria;Rotifera;Bdelloidea | 1;1;1;1;1;1;1;1;0.881748 |
| m54032_170210_154134_30278129_ccs_Otu00077_25_1588_bp | 85 | Eukaryota;SAR;Alveolata;Apicomplexa;Conoidasida;Gregarinasina | 0.995963;0.995963;0.995963;0.995963;0.995963;0.995963 |
| m54032_170210_154134_30671731_ccs_Otu00560_2_1588_bp | 88 | Eukaryota;SAR;Alveolata;Apicomplexa;Conoidasida;Cryptosporida;Cryptosporidium | 0.981634;0.981634;0.981634;0.977376;0.977376;0.977376;0.977376 |
| m54032_170210_154134_31458233_ccs_Otu00169_11_1588_bp | 98.1 | Eukaryota;Opisthokonta;Nucletmycea;Fungi;Dikarya;Ascomycota;Taphrinomycotina;Archaeorhizomycetes;Archaeorhizomycetales;Archaeorhizomycetaceae;Archaeorhizomyces | 0.992608;0.932661;0.932661;0.932661;0.932661;0.932661;0.932661;0.932661;0.932661;0.932661;0.932661 |
| m54032_170210_154134_35259198_ccs_Otu00384_3_1588_bp | 97 | Eukaryota;SAR;Alveolata;Ciliophora;Intramacronucleata;Litostomatea;Haptoria | 1;1;1;1;1;1;1 |
| m54032_170210_154134_36569815_ccs_Otu00626_2_1588_bp | 88.1 | Eukaryota;SAR;Rhizaria;Cercozoa | 0.987945;0.987945;0.987945;0.987945 |
| m54032_170210_154134_38535521_ccs_Otu00371_3_1588_bp | 81.5 | Eukaryota;SAR;Rhizaria;Cercozoa | 0.997326;0.997326;0.997326;0.997326 |
| m54032_170210_154134_41550060_ccs_Otu00454_3_1588_bp | 94.6 | Eukaryota;SAR;Rhizaria;Cercozoa;Glissomonadida;Heteromita | 0.999999;0.999999;0.999999;0.999999;0.999999;0.999999 |
| m54032_170210_154134_42270821_ccs_Otu00426_3_1588_bp | 92.6 | Eukaryota;Opisthokonta;Nucletmycea;Fungi;Dikarya | 1;1;1;1;1 |
| m54032_170210_154134_42729615_ccs_Otu00395_3_1588_bp | 98.1 | Eukaryota;Opisthokonta;Nucletmycea;Fungi;Dikarya;Ascomycota;Pezizomycotina;Leotiomycetes;Thelebolales;Thelebolaceae;Thelebolus | 0.998431;0.998431;0.998431;0.998431;0.998431;0.998431;0.998431;0.998431;0.939804;0.939804;0.939804 |
| m54032_170210_154134_45679577_ccs_Otu00429_3_1588_bp | 98.6 | Eukaryota;Opisthokonta;Nucletmycea;Fungi;Dikarya;Basidiomycota;Agaricomycotina;Agaricomycetes;Agaricales | 1;1;1;1;1;1;1;1;1 |
| m54032_170210_154134_47579417_ccs_Otu00476_2_1588_bp | 96.3 | Eukaryota;SAR;Rhizaria;Cercozoa;Cercomonadidae | 0.994604;0.994604;0.994604;0.994604;0.994604 |
| m54032_170210_154134_48496837_ccs_Otu00375_3_1588_bp | 88.8 | Eukaryota;SAR;Rhizaria;Cercozoa | 0.994215;0.994215;0.994215;0.994215 |
| m54032_170210_154134_49545999_ccs_Otu00410_3_1588_bp | 97.1 | Eukaryota;SAR;Rhizaria;Cercozoa | 1;1;1;1 |
| m54032_170210_154134_50201219_ccs_Otu00495_2_1588_bp | 97.4 | Eukaryota;Opisthokonta;Nucletmycea;Fungi;Dikarya;Ascomycota;Pezizomycotina;Leotiomycetes;Incertae_Sedis;Myxotrichaceae | 0.963871;0.963871;0.963871;0.963871;0.963871;0.963871;0.963871;0.963871;0.963871;0.963871 |
| m54032_170210_154134_50266641_ccs_Otu00002_1322_1588_bp | 75.4 | Eukaryota;SAR;Alveolata;Apicomplexa | |
| m54032_170210_154134_50725473_ccs_Otu00442_3_1588_bp | 99 | Eukaryota;Opisthokonta;Nucletmycea;Fungi;Dikarya;Ascomycota;Pezizomycotina;Sordariomycetes;Xylariales | 0.99971;0.99971;0.99971;0.99971;0.99971;0.99971;0.99971;0.99971;0.99971 |
| m54032_170210_154134_50921770_ccs_Otu00649_2_1588_bp | 95.4 | Eukaryota;SAR;Rhizaria;Cercozoa;Glissomonadida;Heteromita | 1;1;1;1;1;1 |
| m54032_170210_154134_51118331_ccs_Otu00611_2_1588_bp | 91.5 | Eukaryota;SAR;Rhizaria;Cercozoa | 0.98343;0.98343;0.98343;0.98343 |
| m54032_170210_154134_51184124_ccs_Otu00507_2_1588_bp | 98 | Eukaryota;Opisthokonta;Nucletmycea;Fungi;Dikarya;Ascomycota;Pezizomycotina;Leotiomycetes;Thelebolales;Thelebolaceae;Thelebolus | 0.998431;0.998431;0.998431;0.998431;0.998431;0.998431;0.998431;0.998431;0.939795;0.939795;0.939795 |

| query | % similarity | annotation | confidence for each rank |
|---|---|---|---|
| m54032_170210_154134_52757196_ccs_Otu00506_2_1588_bp | 98.2 | Eukaryota;Opisthokonta;Nucletmycea;Fungi;Dikarya;Ascomycota;Pezizomycotina;Leotiomycetes;Thelebolales;Thelebolaceae;Thelebolus | 1;1;1;1;1;1;1;0.997051;0.997051;0.997051 |
| m54032_170210_154134_52887793_ccs_Otu00502_2_1588_bp | 97.1 | Eukaryota;Opisthokonta;Nucletmycea;Fungi;Mucoromycota;Mortierellomycotina;Incertae_Sedis;Mortierellales;Mortierellaceae;Mortierella;verticillata | 0.996202;0.996202;0.996202;0.996202;0.996202;0.996202;0.996202;0.996202;0.996202;0.996202;0.996202 |
| m54032_170210_154134_52888536_ccs_Otu00324_4_1588_bp | 97.1 | Eukaryota;Incertae_Sedis;Breviatea;Breviata;anathema | 1;1;1;0.946544;0.946544 |
| m54032_170210_154134_53019089_ccs_Otu00551_2_1588_bp | 92.6 | Eukaryota;Opisthokonta;Holozoa;Metazoa;Animalia;Eumetazoa;Bilateria;Tardigrada;Eutardigrada;Parachela | 0.99951;0.99951;0.99951;0.99951;0.99951;0.99951;0.99951;0.99906;0.99906;0.99906 |
| m54032_170210_154134_54001751_ccs_Otu00515_2_1588_bp | 89.3 | Eukaryota;SAR;Rhizaria;Cercozoa;Imbricatea;Nudifila | 1;1;1;1;1;0.985618 |
| m54032_170210_154134_55050984_ccs_Otu00404_3_1588_bp | 93.5 | Eukaryota;SAR;Alveolata;Ciliophora;Intramacronucleata;Conthreep;Colpodea | 1;1;1;1;1;1;1 |
| m54032_170210_154134_55641023_ccs_Otu00049_45_1588_bp | 82.5 | Eukaryota;SAR;Rhizaria;Cercozoa;Cercomonadidae | 1;1;1;1;1 |
| m54032_170210_154134_55968023_ccs_Otu00573_2_1588_bp | 97 | Eukaryota;SAR;Stramenopiles;Ochrophyta;Chrysophyceae;Chromulinales;Uroglena | 0.999932;0.999932;0.999932;0.999932;0.999932;0.999932;0.999932 |
| m54032_170210_154134_56099783_ccs_Otu00352_4_1588_bp | 94.3 | Eukaryota;SAR;Stramenopiles;Ochrophyta;Chrysophyceae | 0.992323;0.992323;0.992323;0.992323;0.992323 |
| m54032_170210_154134_58327959_ccs_Otu00512_2_1588_bp | 96.6 | Eukaryota;SAR;Rhizaria;Cercozoa | 0.750674;0.750674;0.750674;0.750674 |
| m54032_170210_154134_58524441_ccs_Otu00538_2_1588_bp | 90 | Eukaryota;Opisthokonta;Nucletmycea;Fungi | 0.805284;0.805284;0.805284;0.805284 |
| m54032_170210_154134_61734990_ccs_Otu00188_9_1588_bp | 84.7 | Eukaryota;Opisthokonta;Nucletmycea;Fungi;Zoopagomycota | 0.991292;0.708901;0.708901;0.708901;0.708901 |
| m54032_170210_154134_61800677_ccs_Otu00424_3_1588_bp | 93.8 | Eukaryota;Opisthokonta;Nucletmycea;Fungi;Mucoromycota;Mucoromycotina;Incertae_Sedis;Endogonales;Endogonaceae | 1;1;1;1;1;1;1;1;1 |
| m54032_170210_154134_62128945_ccs_Otu00412_3_1588_bp | 97.8 | Eukaryota;Opisthokonta;Nucletmycea;Fungi;Dikarya;Ascomycota;Pezizomycotina;Dothideomycetes;Hysteriales;Hysteriaceae;Farlowiella;carmichaeliana | 1;1;1;1;1;1;1;0.9942;0.9942;0.9942;0.9942;0.9942 |
| m54032_170210_154134_63046257_ccs_Otu00149_12_1588_bp | 97.8 | Eukaryota;Opisthokonta;Nucletmycea;Fungi;Dikarya;Basidiomycota;Agaricomycotina;Agaricomycetes;Sebacinales;Sebacinaceae | 0.992627;0.992627;0.992627;0.992627;0.992627;0.992627;0.992627;0.992627;0.992627;0.992627 |
| m54032_170210_154134_64422483_ccs_Otu00405_3_1588_bp | 98.7 | Eukaryota;SAR;Stramenopiles;Ochrophyta;Chrysophyceae | 0.99406;0.99406;0.99406;0.99406;0.99406 |
| m54032_170210_154134_6488584_ccs_Otu00418_3_1588_bp | 94.7 | Eukaryota;SAR;Alveolata;Apicomplexa;Conoidasida;Gregarinasina | 1;1;1;1;1;1 |
| m54032_170210_154134_65601609_ccs_Otu00389_3_1588_bp | 98.7 | Eukaryota;SAR;Alveolata;Ciliophora;Intramacronucleata;Litostomatea;Haptoria;Arcuospathidium | 0.999828;0.999828;0.999828;0.999828;0.999828;0.999828;0.999828;0.999828 |
| m54032_170210_154134_65733281_ccs_Otu00555_2_1588_bp | 84 | Eukaryota;SAR;Alveolata | 0.965488;0.965362;0.965362 |
| m54032_170210_154134_65995478_ccs_Otu00607_2_1588_bp | 92.8 | Eukaryota;SAR;Alveolata;Ciliophora;Intramacronucleata;Conthreep;Colpodea | 1;1;1;1;1;1;1 |
| m54032_170210_154134_6684917_ccs_Otu00177_10_1588_bp | 96 | Eukaryota;SAR;Rhizaria;Cercozoa;Glissomonadida;Heteromita | 1;1;1;1;1;1 |
| m54032_170210_154134_67371271_ccs_Otu00564_2_1588_bp | 96.9 | Eukaryota;Opisthokonta;Nucletmycea;Fungi;Chytridiomycota;Incertae_Sedis;Chytridiomycetes | 0.992966;0.992966;0.992966;0.992966;0.992966;0.992966;0.992966 |
| m54032_170210_154134_67437517_ccs_Otu00251_6_1588_bp | 93.6 | Eukaryota;SAR;Rhizaria;Cercozoa | 0.998252;0.998252;0.998252;0.998252 |
| m54032_170210_154134_69206382_ccs_Otu00489_2_1588_bp | 94.5 | Eukaryota;Incertae_Sedis;Breviatea;Breviata;anathema | 1;1;1;0.779644;0.779644 |
| m54032_170210_154134_69272056_ccs_Otu00589_2_1588_bp | 99.6 | Eukaryota;Opisthokonta;Nucletmycea;Fungi;Dikarya;Ascomycota;Pezizomycotina;Dothideomycetes;Incertae_Sedis;Gloniaceae;Cenococcum;geophilum | 1;1;1;1;1;1;1;1;0.997155;0.997155;0.997155;0.997155 |
| m54032_170210_154134_70255583_ccs_Otu00483_2_1588_bp | 93.6 | Eukaryota;SAR;Alveolata;Apicomplexa;Conoidasida;Gregarinasina;Neogregarinorida;Mattesia;Mattesia;SV2003 | 1;1;1;1;1;1;0.752423;0.752423;0.752423;0.752423 |
| m54032_170210_154134_70779160_ccs_Otu00415_3_1588_bp | 78.2 | Eukaryota;SAR;Alveolata;Apicomplexa | |
| m54032_170210_154134_71107544_ccs_Otu00648_2_1588_bp | 95.8 | Eukaryota;SAR;Rhizaria;Cercozoa;Cercomonadidae | 0.999933;0.999933;0.999933;0.999933;0.999933 |
| m54032_170210_154134_72352254_ccs_Otu00381_3_1588_bp | 96.4 | Eukaryota;SAR;Rhizaria;Cercozoa;Thecofilosea;Cryomonadida;Rhizaspididae;Rhogostoma | 1;1;1;1;1;1;1;1 |
| m54032_170210_154134_7340600_ccs_Otu00317_4_1588_bp | 68.7 | chimera | |
| m54032_170210_154134_74187149_ccs_Otu00513_2_1588_bp | 95.6 | Eukaryota;SAR;Stramenopiles;MAST_12;MAST_12C | 1;1;1;1;0.864306 |
| m54032_170210_154134_74449096_ccs_Otu00618_2_1588_bp | 96.8 | Eukaryota;Opisthokonta;Nucletmycea;Fungi;Dikarya;Basidiomycota;Agaricomycotina;Tremellomycetes;Tremellales | 1;1;1;1;1;1;1;1;1 |
| m54032_170210_154134_8782393_ccs_Otu00200_8_1588_bp | 96.5 | Eukaryota;SAR;Rhizaria;Cercozoa;Cercomonadidae | 0.999387;0.999387;0.999387;0.999387;0.999387 |
| m54032_170210_154134_8847650_ccs_Otu00295_5_1588_bp | 87.7 | Eukaryota;SAR;Alveolata;Apicomplexa;Conoidasida;Cryptosporida;Cryptosporidium;Colpodella | 0.999985;0.999985;0.999985;0.999985;0.999985;0.999985;0.999985;0.893537 |
| m54032_170210_154134_9241405_ccs_Otu00480_2_1588_bp | 86 | Eukaryota;SAR;Rhizaria;Cercozoa | 0.99273;0.99273;0.99273;0.99273 |
| m54088_170214_101554_10027668_ccs_Otu00605_2_1588_bp | 88.4 | Eukaryota;Opisthokonta;Holozoa;Metazoa;Animalia;Eumetazoa;Bilateria;Nematoda;Enoplea;Enoplia;Triplonchida | 1;1;1;1;1;1;1;1;1;1;0.999903 |
| m54088_170214_101554_10945034_ccs_Otu00638_2_1588_bp | 62.3 | chimera | |
| m54088_170214_101554_15401713_ccs_Otu00318_4_1588_bp | 87.7 | Eukaryota;SAR;Alveolata;Apicomplexa;Conoidasida;Gregarinasina;Eugregarinorida;Syncystis;mirabilis | 1;1;1;1;1;1;0.674781;0.674781;0.674781 |
| m54088_170214_101554_15926018_ccs_Otu00634_2_1588_bp | 95.5 | Eukaryota;Opisthokonta;Nucletmycea;Fungi;Chytridiomycota;Incertae_Sedis;Chytridiomycetes;Chytridiales;Chytridiaceae | 1;1;1;1;1;1;1;0.99921;0.99921 |
| m54088_170214_101554_16843333_ccs_Otu00453_3_1588_bp | 97.9 | Eukaryota;Opisthokonta;Holozoa;Metazoa;Animalia;Eumetazoa;Bilateria;Tardigrada;Eutardigrada;Parachela | 1;1;1;1;1;1;1;1;1;1 |
| m54088_170214_101554_18547470_ccs_Otu00333_4_1588_bp | 87.2 | Eukaryota;SAR;Rhizaria;Cercozoa;Vampyrellidae | 1;1;1;1;1 |
| m54088_170214_101554_19923796_ccs_Otu00539_2_1588_bp | 99.4 | Eukaryota;SAR;Rhizaria;Cercozoa;Imbricatea;Nudifila | 0.997494;0.997494;0.997494;0.997494;0.997494;0.997494 |
| m54088_170214_101554_20709485_ccs_Otu00580_2_1588_bp | 98.4 | Eukaryota;SAR;Stramenopiles;Ochrophyta;Diatomea;Bacillariophytina;Bacillariophyceae;Fistulifera;Navicula;JB12 | 0.999969;0.999969;0.999969;0.999969;0.999969;0.999969;0.999969;0.999969;0.989244;0.989244 |
| m54088_170214_101554_21365230_ccs_Otu00390_3_1588_bp | 93.3 | Eukaryota;Opisthokonta;Nucletmycea;Fungi;Cryptomycota | 1;1;1;1;1 |
| m54088_170214_101554_21889432_ccs_Otu00304_5_1588_bp | 95.4 | Eukaryota;SAR;Rhizaria;Cercozoa;Glissomonadida;Heteromita | 0.999998;0.999998;0.999998;0.999998;0.999998;0.999998 |
| m54088_170214_101554_21955249_ccs_Otu00199_8_1588_bp | 92.3 | Eukaryota;Opisthokonta;Nucletmycea;Fungi;Dikarya;Ascomycota;Saccharomycotina;Saccharomycetes;Saccharomycetales;Dipodascaceae;Geotrichum | 1;1;1;1;1;1;1;1;1;0.531374;0.531374 |
| m54088_170214_101554_22479816_ccs_Otu00520_2_1588_bp | 94.3 | Eukaryota;Opisthokonta;Nucletmycea;Fungi;Dikarya;Basidiomycota;Pucciniomycotina;Pucciniomycetes;Platygloeales | 1;1;1;1;1;1;1;1;0.999304 |
| m54088_170214_101554_22806837_ccs_Otu00647_2_1588_bp | 90.4 | Eukaryota;SAR;Rhizaria;Cercozoa;Vampyrellidae | 1;1;1;1;1 |
| m54088_170214_101554_23396962_ccs_Otu00612_2_1588_bp | 96.5 | Eukaryota;SAR;Rhizaria;Cercozoa;Glissomonadida;Bodomorpha | 0.999996;0.999996;0.999996;0.999996;0.999996;0.999996 |
| m54088_170214_101554_24510766_ccs_Otu00373_3_1588_bp | 98.4 | Eukaryota;Opisthokonta;Nucletmycea;Fungi;Chytridiomycota;Incertae_Sedis;Chytridiomycetes | 0.988345;0.988345;0.988345;0.988345;0.988345;0.988345;0.988345 |
| m54088_170214_101554_24838546_ccs_Otu00508_2_1588_bp | 98 | Eukaryota;SAR;Stramenopiles;Peronosporomycetes;Pythium;rostratifingens | 0.999054;0.999054;0.999054;0.999054;0.999054;0.957565 |
| m54088_170214_101554_25297533_ccs_Otu00150_12_1588_bp | 86.2 | Eukaryota;Opisthokonta;Holozoa;Metazoa;Animalia;Eumetazoa;Bilateria;Nematoda;Enoplea;Dorylaimia | 1;1;1;1;1;1;1;1;1;1 |
| m54088_170214_101554_26215023_ccs_Otu00525_2_1588_bp | 97.9 | Eukaryota;SAR;Alveolata;Ciliophora;Intramacronucleata;Litostomatea;Haptoria | 0.98671;0.98671;0.98671;0.98671;0.98671;0.98671;0.98671 |
| m54088_170214_101554_26935601_ccs_Otu00125_14_1588_bp | 99 | Eukaryota;Opisthokonta;Holozoa;Metazoa;Animalia;Eumetazoa;Bilateria;Rotifera;Bdelloidea | 1;1;1;1;1;1;1;1;1 |
| m54088_170214_101554_27001758_ccs_Otu00519_2_1588_bp | 91.9 | Eukaryota;SAR;Alveolata;Protalveolata;Perkinsidae | 1;1;1;1;1 |
| m54088_170214_101554_29098954_ccs_Otu00487_2_1588_bp | 93.3 | Eukaryota;SAR;Alveolata;Ciliophora;Intramacronucleata;Conthreep;Colpodea;Colpodida | 0.992228;0.992228;0.992228;0.992228;0.992228;0.992228;0.992228;0.992228 |
| m54088_170214_101554_29885244_ccs_Otu00183_9_1588_bp | 89 | Eukaryota;SAR;Alveolata;Apicomplexa;Conoidasida;Gregarinasina;Eugregarinorida;Syncystis;mirabilis | 1;1;1;1;1;1;0.746032;0.746032;0.746032 |
| m54088_170214_101554_30343641_ccs_Otu00594_2_1588_bp | 97 | Eukaryota;Opisthokonta;Nucletmycea;Fungi;Dikarya;Basidiomycota;Agaricomycotina;Tremellomycetes;Tremellales;Bulleribasidiaceae;Vishniacozyma | 0.951602;0.951602;0.951602;0.951602;0.951602;0.951602;0.951602;0.951602;0.951602;0.943805;0.943805 |
| m54088_170214_101554_32637044_ccs_Otu00497_2_1588_bp | 79.6 | Eukaryota;SAR;Rhizaria;Cercozoa;Vampyrellidae | 1;1;1;1;1 |

| query | % similarity | annotation | confidence for each rank |
|---|---|---|---|
| m54088_170214_101554_32768766_ccs_Otu00585_2_1588_bp | 99.1 | Eukaryota;Opisthokonta;Nucletmycea;Fungi;Dikarya;Ascomycota;Saccharomycotina;Saccharomycetes;Saccharomycetales;Debaryomycetaceae;Debaryomyces | 0.999949;0.999949;0.999949;0.999949;0.999949;0.999949;0.999949;0.999949;0.999949;0.995975;0.995975 |
| m54088_170214_101554_35193430_ccs_Otu00441_3_1588_bp | 97.1 | Eukaryota;SAR;Rhizaria;Cercozoa;Glissomonadida | 0.989071;0.989071;0.989071;0.989071;0.978433 |
| m54088_170214_101554_35848711_ccs_Otu00214_8_1588_bp | 78.9 | Eukaryota;SAR;Rhizaria;Cercozoa;Phytomyxea | 0.99929;0.99929;0.99929;0.99929;0.99929 |
| m54088_170214_101554_36438655_ccs_Otu00425_3_1588_bp | 92.2 | Eukaryota;SAR;Alveolata;Ciliophora;Intramacronucleata;Spirotrichea;Euplotia;Aspidisca | 1;1;1;1;1;1;1;0.593202 |
| m54088_170214_101554_36897706_ccs_Otu00614_2_1588_bp | 71.9 | Eukaryota;Haptophyta | 0.987475;0.831718 |
| m54088_170214_101554_38535673_ccs_Otu00071_28_1588_bp | 98.7 | Eukaryota;SAR;Alveolata;Ciliophora;Intramacronucleata;Litostomatea;Haptoria | 0.998432;0.998432;0.998432;0.998432;0.998432;0.998432;0.998432 |
| m54088_170214_101554_41091495_ccs_Otu00574_2_1588_bp | 88.3 | Eukaryota;SAR;Alveolata;Apicomplexa;Conoidasida;Gregarinasina;Eugregarinorida;Syncystis;mirabilis | 1;1;1;1;1;1;0.704892;0.704892;0.704892 |
| m54088_170214_101554_42140459_ccs_Otu00380_3_1588_bp | 75.2 | Eukaryota;Opisthokonta;Holozoa;Metazoa;Animalia;Eumetazoa;Bilateria | 0.969844;0.969844;0.969844;0.969844;0.969844;0.969844;0.969844 |
| m54088_170214_101554_43778960_ccs_Otu00316_4_1588_bp | 98.8 | Eukaryota;Opisthokonta;Holozoa;Metazoa;Animalia;Eumetazoa;Bilateria;Mollusca;Gastropoda;Heterobranchia | 0.996791;0.996791;0.996791;0.996791;0.996791;0.996791;0.996791;0.996791;0.996791;0.996791 |
| m54088_170214_101554_44761499_ccs_Otu00365_3_1588_bp | 77.7 | Eukaryota;SAR;Rhizaria;Cercozoa;Phytomyxea | 0.99788;0.99788;0.99788;0.99788;0.99788 |
| m54088_170214_101554_45351375_ccs_Otu00456_3_1588_bp | 90.1 | Eukaryota;Opisthokonta;Nucletmycea;Fungi;Cryptomycota;Incertae_Sedis;Incertae_Sedis;Incertae_Sedis;Incertae_Sedis;Rozella | 0.993278;0.993278;0.993278;0.794448;0.794448;0.794448;0.794448;0.794448;0.794448;0.794448 |
| m54088_170214_101554_46858937_ccs_Otu00579_2_1588_bp | 94.4 | Eukaryota;Opisthokonta;Holozoa;Metazoa;Animalia;Eumetazoa;Bilateria;Nematoda;Chromadorea;Rhabditida | 1;1;1;1;1;1;1;1;1;1 |
| m54088_170214_101554_47120562_ccs_Otu00492_2_1588_bp | 92.4 | Eukaryota;Opisthokonta;Nucletmycea;Fungi;LKM15 | 1;1;1;1;1 |
| m54088_170214_101554_48300108_ccs_Otu00504_2_1588_bp | 95.8 | Eukaryota;Opisthokonta;Holozoa;Metazoa;Animalia;Eumetazoa;Bilateria;Nematoda;Enoplea;Dorylaimia;Dorylaimida | 1;1;1;1;1;1;1;1;1;1;1 |
| m54088_170214_101554_49676803_ccs_Otu00635_2_1588_bp | 95.3 | Eukaryota;Opisthokonta;Holozoa;Metazoa;Animalia;Eumetazoa;Bilateria;Nematoda;Chromadorea;Tylenchida | 1;1;1;1;1;1;1;1;1;1 |
| m54088_170214_101554_51250069_ccs_Otu00547_2_1588_bp | 80.7 | Eukaryota;Opisthokonta;Nucletmycea;Fungi;Zoopagomycota;Entomophthoromycotina;Entomophthoromycetes;Entomophthorales;Entomophthoraceae;Entomophthora;muscae | 1;1;1;1;0.536155;0.536155;0.536155;0.536155;0.536155;0.536155;0.536155 |
| m54088_170214_101554_51839075_ccs_Otu00445_3_1588_bp | 84.9 | Eukaryota;Opisthokonta;Holozoa;Metazoa;Animalia;Eumetazoa;Bilateria;Nematoda;Chromadorea;Tylenchida | 1;1;1;1;1;1;1;1;1;0.768214 |
| m54088_170214_101554_52822462_ccs_Otu00408_3_1588_bp | 93.7 | Eukaryota;SAR;Stramenopiles;Ochrophyta;Xanthophyceae;Tribonematales | 0.994452;0.994452;0.994452;0.994452;0.994452;0.994452 |
| m54088_170214_101554_54263951_ccs_Otu00455_3_1588_bp | 87.7 | Eukaryota;SAR;Rhizaria;Cercozoa;Vampyrellidae | 0.999962;0.999962;0.999962;0.999962;0.999962 |
| m54088_170214_101554_54461103_ccs_Otu00443_3_1588_bp | 99.2 | Eukaryota;SAR;Rhizaria;Cercozoa;Cercomonadidae | 1;1;1;1;1 |
| m54088_170214_101554_55181446_ccs_Otu00475_2_1588_bp | 98.3 | Eukaryota;Archaeplastida;Chloroplastida;Chlorophyta;Trebouxiophyceae;Prasiolales;Stichococcus | 0.999622;0.999622;0.999622;0.999622;0.999622;0.999622;0.999622 |
| m54088_170214_101554_55378723_ccs_Otu00439_3_1588_bp | 97.3 | Eukaryota;Opisthokonta;Nucletmycea;Fungi;Chytridiomycota;Incertae_Sedis;Chytridiomycetes;Spizellomycetales;Spizellomycetaceae | 0.981415;0.981415;0.981415;0.981415;0.981415;0.981415;0.981415;0.964116;0.964116 |
| m54088_170214_101554_57541528_ccs_Otu00292_5_1588_bp | 98.9 | Eukaryota;SAR;Alveolata;Ciliophora;Intramacronucleata;Litostomatea;Haptoria;Apobryophyllum | 1;1;1;1;1;1;1;1 |
| m54088_170214_101554_5898454_ccs_Otu00136_13_1588_bp | 98.2 | Eukaryota;Opisthokonta;Holozoa;Metazoa;Animalia;Eumetazoa;Bilateria;Nematoda;Chromadorea;Tylenchida;Filenchus | 0.991352;0.991352;0.991352;0.991352;0.991352;0.991352;0.991352;0.991352;0.991352;0.991352;0.991352 |
| m54088_170214_101554_59572300_ccs_Otu00588_2_1588_bp | 84.3 | Eukaryota;Opisthokonta;Nucletmycea;Fungi;Cryptomycota | 0.993794;0.993794;0.993794;0.993794;0.993794 |
| m54088_170214_101554_59769536_ccs_Otu00209_8_1588_bp | 94.8 | Eukaryota;SAR;Rhizaria;Cercozoa;Glissomonadida;Heteromita | 0.99943;0.99943;0.99943;0.99943;0.99943;0.99943 |
| m54088_170214_101554_61473411_ccs_Otu00296_5_1588_bp | 99.1 | Eukaryota;Opisthokonta;Holozoa;Metazoa;Animalia;Eumetazoa;Bilateria;Nematoda;Chromadorea;Rhabditida;Eucephalobus;striatus | 0.99999;0.99999;0.99999;0.99999;0.99999;0.99999;0.99999;0.99999;0.99999;0.99999;0.660132;0.660132 |
| m54088_170214_101554_61735389_ccs_Otu00595_2_1588_bp | 88.7 | Eukaryota;SAR;Alveolata;Apicomplexa;Conoidasida;Cryptosporida;Cryptosporidium;Colpodella | 1;1;1;1;1;1;1;1 |
| m54088_170214_101554_62718575_ccs_Otu00171_10_1588_bp | 98.4 | Eukaryota;Archaeplastida;Chloroplastida;Charophyta;Phragmoplastophyta;Streptophyta;Embryophyta;Tracheophyta;Spermatophyta;Magnoliophyta;Brassicales;Brassica | 1;1;1;1;1;1;1;1;1;1;1;1 |
| m54088_170214_101554_62915227_ccs_Otu00320_4_1588_bp | 96.9 | Eukaryota;Opisthokonta;Holozoa;Metazoa;Animalia;Eumetazoa;Bilateria;Nematoda;Chromadorea;Monhysterida | 1;1;1;1;1;1;1;1;1;1 |
| m54088_170214_101554_63636059_ccs_Otu00041_53_1588_bp | 98.2 | Eukaryota;SAR;Alveolata;Ciliophora;Intramacronucleata;Conthreep;Colpodea;Colpodida | 0.997867;0.997867;0.997867;0.997867;0.997867;0.997867;0.997867;0.997815 |
| m54088_170214_101554_63898102_ccs_Otu00629_2_1588_bp | 96.7 | Eukaryota;SAR;Stramenopiles;Peronosporomycetes;Pythium | 0.997683;0.997683;0.997683;0.997683;0.997683 |
| m54088_170214_101554_64553700_ccs_Otu00534_2_1588_bp | 87 | Eukaryota;SAR;Rhizaria;Cercozoa | 0.999843;0.999843;0.999843;0.999843 |
| m54088_170214_101554_67044099_ccs_Otu00608_2_1588_bp | 96.1 | Eukaryota;Opisthokonta;Nucletmycea;Fungi;Dikarya;Ascomycota;Pezizomycotina;Sordariomycetes;Hypocreales | 0.965684;0.965684;0.965684;0.965684;0.965684;0.965684;0.965684;0.965684;0.936152 |
| m54088_170214_101554_67698771_ccs_Otu00140_13_1588_bp | 89 | Eukaryota;Opisthokonta;Holozoa;Metazoa;Animalia;Eumetazoa;Bilateria;Nematoda;Chromadorea;Monhysterida;Eumonhystera | 1;1;1;1;1;1;1;1;1;1;0.565688 |
| m54088_170214_101554_69533769_ccs_Otu00403_3_1588_bp | 95.6 | Eukaryota;Opisthokonta;Nucletmycea;Fungi;Chytridiomycota;Incertae_Sedis;Chytridiomycetes;Rhizophydiales;Rhizophydiaceae;Rhizophydium | 1;1;1;1;1;1;1;1;1;1 |
| m54088_170214_101554_7012738_ccs_Otu00270_6_1588_bp | 98.6 | Eukaryota;Archaeplastida;Chloroplastida;Chlorophyta;Ulvophyceae;Planophila;SAG_32_98 | 1;1;1;1;1;0.934232;0.934232 |
| m54088_170214_101554_73925144_ccs_Otu00121_15_1588_bp | 71.3 | Eukaryota;Haptophyta | 0.963267;0.956535 |
| m54088_170214_101554_8978932_ccs_Otu00632_2_1588_bp | 91.6 | Eukaryota;SAR;Rhizaria;Cercozoa;Cercomonadidae | 0.990908;0.990908;0.990908;0.990908;0.990908 |
| m54088_170214_101554_9044787_ccs_Otu00322_4_1588_bp | 96.3 | Eukaryota;Opisthokonta;Nucletmycea;Fungi;Dikarya;Ascomycota;Pezizomycotina;Sordariomycetes;Sordariales;Chaetomiaceae;Chaetomium | 1;1;1;1;1;1;1;1;1;1;0.997601 |
| m54088_170214_101554_9110177_ccs_Otu00387_3_1588_bp | 95.7 | Eukaryota;SAR;Rhizaria;Cercozoa;Phytomyxea;Polymyxa;graminis | 1;1;1;1;1;0.875795;0.875795 |
| m54088_170214_101554_9306600_ccs_Otu00477_2_1588_bp | 85.1 | Eukaryota;Opisthokonta;Holozoa;Metazoa;Animalia;Eumetazoa;Bilateria;Nematoda;Enoplea;Dorylaimia | 1;1;1;1;1;1;1;1;1;1 |
| m54088_170214_101554_9634642_ccs_Otu00617_2_1588_bp | 94.3 | Eukaryota;Opisthokonta;Nucletmycea;Fungi;Chytridiomycota;Incertae_Sedis;Chytridiomycetes | 1;1;1;1;1;1;1 |
| m54088_170214_101554_9765633_ccs_Otu00451_3_1588_bp | 90.1 | Eukaryota;Opisthokonta;Holozoa;Metazoa;Animalia;Eumetazoa;Bilateria;Nematoda;Chromadorea;Rhabditida | 0.992351;0.992351;0.992351;0.992351;0.992351;0.992351;0.992351;0.992351;0.992351;0.992351 |
| m54088_170214_101554_9961801_ccs_Otu00267_6_1588_bp | 79.7 | Eukaryota;Opisthokonta;Holozoa;Metazoa;Animalia;Eumetazoa;Bilateria;Nematoda;Chromadorea;Rhabditida | 1;1;1;1;1;1;1;1;1;1 |
| m54088_170216_085445_10486515_ccs_Otu00541_2_1588_bp | 84.1 | Eukaryota;SAR;Rhizaria;Cercozoa;Vampyrellidae | 0.999987;0.999987;0.999987;0.999987;0.999987 |
| m54088_170216_085445_11010804_ccs_Otu00482_2_1588_bp | 92.4 | Eukaryota;SAR;Rhizaria;Cercozoa | 0.99886;0.99886;0.99886;0.99886 |
| m54088_170216_085445_11075909_ccs_Otu00262_6_1588_bp | 99 | Eukaryota;Opisthokonta;Holozoa;Metazoa;Animalia;Eumetazoa;Bilateria;Nematoda;Chromadorea;Tylenchida | 0.99957;0.99957;0.99957;0.99957;0.99957;0.99957;0.99957;0.99957;0.99957;0.99957 |
| m54088_170216_085445_11534828_ccs_Otu00327_4_1588_bp | 87.8 | Eukaryota;SAR;Alveolata;Apicomplexa;Conoidasida;Cryptosporida;Cryptosporidium;Colpodella | 1;1;1;1;1;1;1;0.979499 |
| m54088_170216_085445_11600425_ccs_Otu00278_5_1588_bp | 91.9 | Eukaryota;SAR;Rhizaria;Cercozoa;Cercomonadidae | 1;1;1;1;1 |
| m54088_170216_085445_12058985_ccs_Otu00562_2_1588_bp | 93.8 | Eukaryota;Opisthokonta;Nucletmycea;Fungi;Cryptomycota | 1;1;1;1;1 |
| m54088_170216_085445_12124471_ccs_Otu00521_2_1588_bp | 97.8 | Eukaryota;SAR;Alveolata;Ciliophora;Intramacronucleata;Litostomatea;Haptoria | 1;1;1;1;1;1;1 |
| m54088_170216_085445_12452784_ccs_Otu00536_2_1588_bp | 87.2 | Eukaryota;SAR;Alveolata;Apicomplexa;Conoidasida;Gregarinasina;Archigregarinorida;Selenidium | 1;1;1;1;1;1;1;1 |
| m54088_170216_085445_12517891_ccs_Otu00543_2_1588_bp | 92.8 | Eukaryota;Opisthokonta;Nucletmycea;Fungi;Dikarya;Basidiomycota;Pucciniomycotina | 0.999768;0.999768;0.999768;0.999768;0.999768;0.999768;0.999768 |
| m54088_170216_085445_12714314_ccs_Otu00302_5_1588_bp | 99.3 | Eukaryota;Archaeplastida;Chloroplastida;Chlorophyta;Trebouxiophyceae;Incertae_Sedis;Neocystis | 0.998885;0.998885;0.998885;0.998885;0.98369;0.98369;0.98369 |
| m54088_170216_085445_13173047_ccs_Otu00474_2_1588_bp | 96.1 | Eukaryota;SAR;Alveolata;Ciliophora;Intramacronucleata;Litostomatea;Haptoria | 0.720751;0.720751;0.720751;0.720751;0.720751;0.720751;0.720751 |
| m54088_170216_085445_13501296_ccs_Otu00572_2_1588_bp | 94.2 | Eukaryota;Opisthokonta;Nucletmycea;Fungi;Chytridiomycota;Incertae_Sedis;Chytridiomycetes | 1;1;1;1;1;1;1 |
| m54088_170216_085445_13566238_ccs_Otu00639_2_1588_bp | 96.4 | Eukaryota;SAR;Rhizaria;Cercozoa;Imbricatea;Nudifila | 1;1;1;1;1;0.869803 |
| m54088_170216_085445_13631611_ccs_Otu00452_3_1588_bp | 99.4 | Eukaryota;Opisthokonta;Nucletmycea;Fungi;Zoopagomycota;Entomophthoromycotina;Basidiobolomycetes;Basidiobolales;Basidiobolaceae;Basidiobolus | 0.999203;0.999203;0.999203;0.999203;0.999203;0.999203;0.999203;0.999203;0.999203;0.999203 |

| query | % similarity | annotation | confidence for each rank |
|---|---|---|---|
| m54088_170216_085445_14025583_ccs_Otu00584_2_1588_bp | 94.9 | Eukaryota;Opisthokonta;Nucletmycea;Fungi;Dikarya;Ascomycota;Pezizomycotina;Pezizomycetes;Pezizales;Ascobolaceae;Thecotheus;holmskjoldii | 0.999993;0.999993;0.999993;0.999993;0.999993;0.999993;0.999993;0.999993;0.999993;0.533398;0.533398;0.533398 |
| m54088_170216_085445_14156144_ccs_Otu00205_8_1588_bp | 90.4 | Eukaryota;Amoebozoa | 0.997746;0.997746 |
| m54088_170216_085445_14222207_ccs_Otu00479_2_1588_bp | 91.7 | Eukaryota;Opisthokonta;Nucletmycea;Fungi;Cryptomycota | 1;1;1;1;1 |
| m54088_170216_085445_14418261_ccs_Otu00416_3_1588_bp | 90.3 | Eukaryota;SAR;Rhizaria;Cercozoa;Imbricatea;Silicofilosea | 1;1;1;1;1;0.84135 |
| m54088_170216_085445_14418389_ccs_Otu00553_2_1588_bp | 98.2 | Eukaryota;Archaeplastida;Chloroplastida;Chlorophyta;Chlorophyceae | 0.999945;0.999945;0.999945;0.999945;0.998125 |
| m54088_170216_085445_14615057_ccs_Otu00593_2_1588_bp | 73.7 | Eukaryota;Amoebozoa;Tubulinea;Euamoebida | 0.99909;0.99909;0.99909;0.819115 |
| m54088_170216_085445_14746128_ccs_Otu00388_3_1588_bp | 88.1 | Eukaryota;SAR;Rhizaria;Cercozoa;Vampyrellidae | 0.999997;0.999997;0.999997;0.999997;0.999997 |
| m54088_170216_085445_14746490_ccs_Otu00535_2_1588_bp | 97.7 | Eukaryota;SAR;Stramenopiles;Ochrophyta;Diatomea;Bacillariophytina;Bacillariophyceae;Pinnularia | 1;1;1;1;1;1;1;1 |
| m54088_170216_085445_14811749_ccs_Otu00401_3_1588_bp | 92.1 | Eukaryota;SAR;Rhizaria;Cercozoa | 0.997266;0.997266;0.997266;0.997266 |
| m54088_170216_085445_14877140_ccs_Otu00430_3_1588_bp | 86.4 | Eukaryota;SAR;Alveolata;Apicomplexa;Conoidasida;Gregarinasina | 0.999995;0.999995;0.999995;0.999995;0.999995;0.999995 |
| m54088_170216_085445_15008021_ccs_Otu00583_2_1588_bp | 83.7 | Eukaryota;Opisthokonta;Nucletmycea;Fungi;Zoopagomycota;Zoopagomycotina;Incertae_Sedis;Zoopagales | 0.999998;0.999916;0.999916;0.999916;0.999916;0.743543;0.743543;0.743543 |
| m54088_170216_085445_15008703_ccs_Otu00530_2_1588_bp | 85.4 | Eukaryota;SAR;Alveolata;Apicomplexa | |
| m54088_170216_085445_15532752_ccs_Otu00037_68_1588_bp | 95 | Eukaryota;SAR;Alveolata;Ciliophora;Intramacronucleata;Litostomatea;Haptoria | 0.828324;0.828324;0.828324;0.828324;0.828324;0.804621;0.804621 |
| m54088_170216_085445_15598163_ccs_Otu00570_2_1588_bp | 65.5 | Eukaryota;Amoebozoa;Tubulinea;Euamoebida | 0.997094;0.997094;0.997094;0.997094 |
| m54088_170216_085445_15925521_ccs_Otu00542_2_1588_bp | 89.1 | Eukaryota;Opisthokonta;Nucletmycea;Fungi;Dikarya;Ascomycota;Pezizomycotina;Sordariomycetes;Hypocreales | 0.997965;0.997965;0.997965;0.997965;0.997965;0.997965;0.997965;0.997965;0.997965 |
| m54088_170216_085445_15991085_ccs_Otu00110_17_1588_bp | 98.6 | Eukaryota;SAR;Rhizaria;Cercozoa;Imbricatea;Spongomonadida;Spongomonas | 1;1;1;1;1;0.916731;0.916731 |
| m54088_170216_085445_16515600_ccs_Otu00276_5_1588_bp | 99 | Eukaryota;SAR;Stramenopiles;Ochrophyta;Chrysophyceae | 1;1;1;1;1 |
| m54088_170216_085445_17236277_ccs_Otu00613_2_1588_bp | 87.2 | Eukaryota;Opisthokonta;Nucletmycea;Fungi;Cryptomycota | 0.999989;0.999989;0.999989;0.999989;0.999989 |
| m54088_170216_085445_17236353_ccs_Otu00265_6_1588_bp | 88.7 | Eukaryota;SAR;Alveolata;Apicomplexa;Conoidasida;Gregarinasina;Eugregarinorida;Syncystis;mirabilis | 1;1;1;1;1;1;0.673801;0.673801;0.673801 |
| m54088_170216_085445_17302405_ccs_Otu00065_31_1588_bp | 96.7 | Eukaryota;SAR;Alveolata;Ciliophora;Intramacronucleata;Conthreep;Colpodea;Colpodida | 0.99995;0.99995;0.99995;0.99995;0.99995;0.99995;0.99995;0.998118 |
| m54088_170216_085445_17367886_ccs_Otu00524_2_1588_bp | 89.9 | Eukaryota;Opisthokonta;Nucletmycea;Fungi;Chytridiomycota;Incertae_Sedis;Chytridiomycetes;Gromochytriales;Gromochytriaceae | 0.999955;0.999955;0.999955;0.999955;0.999955;0.999955;0.999955;0.592204;0.592204 |
| m54088_170216_085445_17563730_ccs_Otu00194_9_1588_bp | 93.2 | Eukaryota;SAR;Alveolata;Ciliophora;Postciliodesmatophora;Heterotrichea | 0.999958;0.999958;0.999958;0.999958;0.999958;0.999958 |
| m54088_170216_085445_17957571_ccs_Otu00369_3_1588_bp | 84.6 | Eukaryota;SAR;Stramenopiles;Labyrinthulomycetes;Sorodiplophrys | 0.99929;0.99929;0.99929;0.99929;0.99929 |
| m54088_170216_085445_18350587_ccs_Otu00511_2_1588_bp | 97.2 | Eukaryota;Opisthokonta;Holozoa;Metazoa;Animalia;Eumetazoa;Bilateria;Nematoda;Enoplea;Dorylaimia;Dorylaimida | 1;1;1;1;1;1;1;1;1;1;0.917646 |
| m54088_170216_085445_18612763_ccs_Otu00096_21_1588_bp | 99.5 | Eukaryota;Opisthokonta;Holozoa;Metazoa;Animalia;Eumetazoa;Bilateria;Nematoda;Chromadorea;Araeolaimida | 0.999742;0.999742;0.999742;0.999742;0.999742;0.999742;0.999742;0.999742;0.999742;0.999742 |
| m54088_170216_085445_18678296_ccs_Otu00620_2_1588_bp | 95 | Eukaryota;SAR;Rhizaria;Cercozoa;Glissomonadida;Heteromita | 0.996915;0.996915;0.996915;0.996915;0.996915;0.996915 |
| m54088_170216_085445_18743923_ccs_Otu00566_2_1588_bp | 97.3 | Eukaryota;Opisthokonta;Nucletmycea;Fungi;Dikarya;Ascomycota;Pezizomycotina | 0.994954;0.994954;0.994954;0.994954;0.994954;0.994954;0.994954 |
| m54088_170216_085445_18809289_ccs_Otu00409_3_1588_bp | 88.5 | Eukaryota;Opisthokonta;Nucletmycea;Fungi;Cryptomycota | 0.984046;0.984046;0.984046;0.984046;0.984046 |
| m54088_170216_085445_18809347_ccs_Otu00220_8_1588_bp | 99 | Eukaryota;SAR;Rhizaria;Cercozoa;Cercomonadidae | 1;1;1;1;1 |
| m54088_170216_085445_18874910_ccs_Otu00182_9_1588_bp | 87.6 | Eukaryota;Amoebozoa;Tubulinea;Euamoebida | 1;1;1;1 |
| m54088_170216_085445_19137150_ccs_Otu00444_3_1588_bp | 97.6 | Eukaryota;Opisthokonta;Nucletmycea;Fungi;Chytridiomycota;Incertae_Sedis;Chytridiomycetes;Fimicolochytrium;alabamae | 0.999994;0.999994;0.999994;0.999994;0.999994;0.999994;0.999994;0.989593;0.989593 |
| m54088_170216_085445_19202225_ccs_Otu00076_25_1588_bp | 99.8 | Eukaryota;Archaeplastida;Chloroplastida;Chlorophyta;Chlorophyceae;Chlamydomonadales | 1;1;1;1;1;0.999347 |
| m54088_170216_085445_19268401_ccs_Otu00396_3_1588_bp | 96.7 | Eukaryota;SAR;Alveolata;Ciliophora;Intramacronucleata;Spirotrichea;Hypotrichia | 1;1;1;1;1;1;1 |
| m54088_170216_085445_19333452_ccs_Otu00577_2_1588_bp | 82.8 | Eukaryota;SAR;Rhizaria;Cercozoa;Vampyrellidae | 0.989945;0.989945;0.989945;0.989945;0.989945 |
| m54088_170216_085445_19465103_ccs_Otu00367_3_1588_bp | 95 | Eukaryota;Opisthokonta;Holozoa;Metazoa;Animalia;Eumetazoa;Bilateria;Nematoda;Chromadorea;Tylenchida;Malenchus;andrassyi | 1;1;1;1;1;1;1;1;1;1;0.940698;0.940698 |
| m54088_170216_085445_19660986_ccs_Otu00438_3_1588_bp | 88.4 | Eukaryota;Amoebozoa;Discosea;Longamoebia;Centramoebida;Acanthamoeba | 0.999465;0.999465;0.999465;0.999465;0.999465;0.999465 |
| m54088_170216_085445_19726707_ccs_Otu00624_2_1588_bp | 81.1 | Eukaryota;Amoebozoa | 0.999998;0.999998 |
| m54088_170216_085445_20382483_ccs_Otu00226_7_1588_bp | 95.5 | Eukaryota;SAR;Rhizaria;Cercozoa;Imbricatea;Nudifila | 0.999387;0.999387;0.999387;0.999387;0.999387;0.999387 |
| m54088_170216_085445_20775842_ccs_Otu00422_3_1588_bp | 94.5 | Eukaryota;SAR;Stramenopiles;Ochrophyta;Diatomea;Bacillariophytina;Bacillariophyceae;Pinnularia | 0.983727;0.983727;0.983727;0.983727;0.983727;0.983727;0.983727;0.983727 |
| m54088_170216_085445_20972431_ccs_Otu00143_12_1588_bp | 99.1 | Eukaryota;Opisthokonta;Nucletmycea;Fungi;Dikarya;Ascomycota;Pezizomycotina;Sordariomycetes;Hypocreales;Incertae_Sedis;Sarocladium | 1;1;1;1;1;1;1;1;1;0.999032;0.999032 |
| m54088_170216_085445_21495958_ccs_Otu00291_5_1588_bp | 88.7 | Eukaryota;SAR;Alveolata;Apicomplexa;Conoidasida;Gregarinasina;Eugregarinorida;Syncystis;mirabilis | 1;1;1;1;1;1;0.657293;0.657293;0.657293 |
| m54088_170216_085445_21495998_ccs_Otu00650_2_1588_bp | 80 | Eukaryota | 0.713872 |
| m54088_170216_085445_21496217_ccs_Otu00382_3_1588_bp | 97.5 | Eukaryota;SAR;Rhizaria;Cercozoa | 0.97095;0.97095;0.97095;0.97095 |
| m54088_170216_085445_21823776_ccs_Otu00256_6_1588_bp | 96.3 | Eukaryota;SAR;Rhizaria;Cercozoa;Cercomonadidae | 1;1;1;1;1 |
| m54088_170216_085445_22216834_ccs_Otu00437_3_1588_bp | 99.5 | Eukaryota;Archaeplastida;Chloroplastida;Chlorophyta;Chlorophyceae;Chlamydomonadales;Tetracystis | 0.992498;0.992498;0.992498;0.992498;0.992498;0.992498;0.978689 |
| m54088_170216_085445_22282805_ccs_Otu00178_10_1588_bp | 88.3 | Eukaryota;SAR;Rhizaria;Cercozoa;Imbricatea | 0.999128;0.999128;0.999128;0.999128;0.999128 |
| m54088_170216_085445_22348434_ccs_Otu00370_3_1588_bp | 87.9 | Eukaryota;SAR;Alveolata;Apicomplexa;Conoidasida;Gregarinasina;Archigregarinorida;Selenidium | 1;1;1;1;1;1;1;1 |
| m54088_170216_085445_22741555_ccs_Otu00569_2_1588_bp | 94.7 | Eukaryota;Opisthokonta;Holozoa;Metazoa;Animalia;Eumetazoa;Bilateria;Nematoda;Chromadorea;Tylenchida;Ditylenchus | 0.998724;0.998724;0.998724;0.998724;0.998724;0.998724;0.998724;0.998724;0.998724;0.998724;0.998724 |
| m54088_170216_085445_23069176_ccs_Otu00628_2_1588_bp | 94.8 | Eukaryota;Amoebozoa;Tubulinea;Euamoebida;BOLA868 | 0.999966;0.999966;0.999966;0.999966;0.999451 |
| m54088_170216_085445_23069283_ccs_Otu00523_2_1588_bp | 84.1 | Eukaryota;Opisthokonta;Nucletmycea;Fungi;Zoopagomycota | 0.999947;0.895072;0.895072;0.895072;0.895072 |
| m54088_170216_085445_23593712_ccs_Otu00556_2_1588_bp | 75.9 | Eukaryota;Excavata;Discoba | 0.988818;0.987918;0.987918 |
| m54088_170216_085445_23659089_ccs_Otu00315_4_1588_bp | 98.7 | Eukaryota;Opisthokonta;Holozoa;Metazoa;Animalia;Eumetazoa;Bilateria;Annelida;Clitellata;Oligochaeta;Haplotaxida;Rhyacodrilus;falciformis | 0.996096;0.996096;0.996096;0.996096;0.996096;0.996096;0.996096;0.996096;0.996096;0.996096;0.996096;0.955768;0.955768 |
| m54088_170216_085445_23986877_ccs_Otu00616_2_1588_bp | 77 | Eukaryota;Opisthokonta;Nucletmycea | 0.979574;0.979574;0.979574 |
| m54088_170216_085445_24380309_ccs_Otu00393_3_1588_bp | 94.6 | Eukaryota;Opisthokonta;Nucletmycea;Fungi;Chytridiomycota;Incertae_Sedis;Chytridiomycetes;Rhizophydiales;Rhizophydiaceae | 1;1;1;1;1;1;1;1;1 |
| m54088_170216_085445_24510743_ccs_Otu00312_4_1588_bp | 99.5 | Eukaryota;SAR;Stramenopiles;Peronosporomycetes;Pythium | 0.990385;0.990385;0.990385;0.990385;0.990385 |
| m54088_170216_085445_24510863_ccs_Otu00253_6_1588_bp | 98.5 | Eukaryota;SAR;Alveolata;Protalveolata;Colpodellida;Colpodella | 0.999688;0.999688;0.999688;0.999609;0.999609;0.999609 |
| m54088_170216_085445_24707166_ccs_Otu00334_4_1588_bp | 97.8 | Eukaryota;Archaeplastida;Chloroplastida;Chlorophyta;Chlorophyceae;Chlamydomonadales;Tetracystis;diplobionticoidea | 1;1;1;1;1;1;0.623889;0.623889 |
| m54088_170216_085445_25363099_ccs_Otu00144_12_1588_bp | 93.9 | Eukaryota;Opisthokonta;Nucletmycea;Fungi;LKM15 | 1;1;1;1;1 |
| m54088_170216_085445_25428597_ccs_Otu00147_12_1588_bp | 95.8 | Eukaryota;SAR;Rhizaria;Cercozoa;Glissomonadida;Heteromita | 0.995874;0.995874;0.995874;0.995874;0.995874;0.995874 |

| query | % similarity | annotation | confidence for each rank |
|---|---|---|---|
| m54088_170216_085445_25756540_ccs_Otu00527_2_1588_bp | 93.1 | Eukaryota;Opisthokonta;Holozoa;Metazoa;Animalia;Eumetazoa;Bilateria;Arthropoda;Hexapoda;Ellipura;Collembola | 0.999903;0.999903;0.999903;0.999903;0.999903;0.999903;0.999903;0.999903;0.99713;0.99713;0.99713 |
| m54088_170216_085445_26149163_ccs_Otu00458_3_1588_bp | 90.4 | Eukaryota;Opisthokonta;Nucletmycea;Fungi;Cryptomycota | 1;1;1;1;1 |
| m54088_170216_085445_26215025_ccs_Otu00202_8_1588_bp | 99.3 | Eukaryota;Opisthokonta;Nucletmycea;Fungi;Dikarya;Ascomycota;Pezizomycotina;Leotiomycetes;Helotiales;Helotiaceae;Hymenoscyphus;repandus | 1;1;1;1;1;1;1;1;1;0.99039;0.99039;0.99039 |
| m54088_170216_085445_26411408_ccs_Otu00498_2_1588_bp | 84.5 | Eukaryota;SAR;Rhizaria;Cercozoa;Phytomyxea | 1;1;1;1;1 |
| m54088_170216_085445_26411972_ccs_Otu00640_2_1588_bp | 79.9 | Eukaryota;Amoebozoa;Tubulinea;Euamoebida | 1;1;1;1 |
| m54088_170216_085445_26608194_ccs_Otu00228_7_1588_bp | 95.5 | Eukaryota;Amoebozoa;Discosea;Flabellinia;Vannellida;Platyamoeba;Platyamoeba;placida | 0.999756;0.999756;0.999756;0.999756;0.999756;0.854706;0.854706;0.854706 |
| m54088_170216_085445_26739551_ccs_Otu00227_7_1588_bp | 94.2 | Eukaryota;SAR;Stramenopiles;Ochrophyta;Chrysophyceae;Incertae_Sedis;Chlamydomyxa | 0.99993;0.99993;0.99993;0.99993;0.99993;0.99993;0.99993 |
| m54088_170216_085445_27001616_ccs_Otu00355_4_1588_bp | 96.2 | Eukaryota;SAR;Rhizaria;Cercozoa;Cercomonadidae;Metabolomonas | 1;1;1;1;1;0.832502 |
| m54088_170216_085445_27001832_ccs_Otu00499_2_1588_bp | 90.6 | Eukaryota;Opisthokonta;Nucletmycea;Fungi | 0.999959;0.999959;0.999959;0.999959 |
| m54088_170216_085445_27329184_ccs_Otu00308_4_1588_bp | 84.5 | Eukaryota;Opisthokonta;Nucletmycea;Fungi;LKM15 | 0.999956;0.999956;0.999956;0.999956;0.999956 |
| m54088_170216_085445_27787507_ccs_Otu00339_4_1588_bp | 95.3 | Eukaryota;SAR;Stramenopiles;Ochrophyta;Diatomea;Bacillariophytina;Bacillariophyceae;Surirella;angusta | 0.996968;0.996968;0.996968;0.996968;0.996968;0.996968;0.996968;0.996968;0.954253 |
| m54088_170216_085445_27853223_ccs_Otu00372_3_1588_bp | 94.7 | Eukaryota;Opisthokonta;Holozoa;Metazoa;Animalia;Eumetazoa;Bilateria;Nematoda;Chromadorea;Tylenchida | 1;1;1;1;1;1;1;1;1;1 |
| m54088_170216_085445_27984169_ccs_Otu00554_2_1588_bp | 90.9 | Eukaryota;SAR;Alveolata;Apicomplexa;Conoidasida;Cryptosporida;Cryptosporidium;Colpodella | 1;1;1;1;1;1;1;1 |
| m54088_170216_085445_28050397_ccs_Otu00297_5_1588_bp | 93.7 | Eukaryota;SAR;Rhizaria;Cercozoa;Incertae_Sedis;Gymnophrys | 0.999998;0.999998;0.999998;0.999998;0.999998;0.999998 |
| m54088_170216_085445_28115316_ccs_Otu00360_3_1588_bp | 92.5 | Eukaryota;Opisthokonta;Nucletmycea;Fungi;Cryptomycota;LKM11 | 0.990818;0.990818;0.990818;0.990818;0.990818;0.570115 |
| m54088_170216_085445_28705118_ccs_Otu00104_18_1588_bp | 98.1 | Eukaryota;SAR;Alveolata;Ciliophora;Intramacronucleata;Conthreep;Colpodea;Platyophryida | 1;1;1;1;1;1;1;0.991014 |
| m54088_170216_085445_28770867_ccs_Otu00361_3_1588_bp | 94 | Eukaryota;Opisthokonta;Nucletmycea;Fungi;Mucoromycota;Glomeromycotina;Glomeromycetes;Glomeromycotina | 1;1;1;1;1;1;1;0.670105 |
| m54088_170216_085445_28901655_ccs_Otu00195_9_1588_bp | 89.2 | Eukaryota;SAR;Rhizaria;Cercozoa;Phytomyxea | 1;1;1;1;1 |
| m54088_170216_085445_29098128_ccs_Otu00469_2_1588_bp | 94.3 | Eukaryota;SAR;Stramenopiles;Labyrinthulomycetes | 0.995078;0.995078;0.995078;0.995078 |
| m54088_170216_085445_29098653_ccs_Otu00311_4_1588_bp | 98.2 | Eukaryota;Archaeplastida;Chloroplastida;Chlorophyta;Chlorophyceae | 0.999999;0.999999;0.999999;0.999999;0.999999 |
| m54088_170216_085445_29164066_ccs_Otu00622_2_1588_bp | 98.4 | Eukaryota;SAR;Alveolata;Ciliophora;Intramacronucleata;Conthreep;Oligohymenophorea;Scuticociliatia;Homalogastra | 1;1;1;1;1;1;1;1;0.524127 |
| m54088_170216_085445_29295581_ccs_Otu00586_2_1588_bp | 98.8 | Eukaryota;Archaeplastida;Chloroplastida;Chlorophyta;Chlorophyceae;Chlamydomonadales;Spongiochloris;spongiosa | 1;1;1;1;1;0.524476;0.524476;0.524476 |
| m54088_170216_085445_29688018_ccs_Otu00428_3_1588_bp | 98.2 | Eukaryota;SAR;Stramenopiles;Ochrophyta;Chrysophyceae;Chromulinales;JBNA46 | 0.993763;0.993763;0.993763;0.993763;0.993763;0.993763;0.871779 |
| m54088_170216_085445_29950661_ccs_Otu00402_3_1588_bp | 99.2 | Eukaryota;Opisthokonta;Nucletmycea;Fungi;Dikarya;Ascomycota;Pezizomycotina;Orbiliomycetes;Orbiliales;Orbiliaceae | 0.997577;0.997577;0.997577;0.997577;0.997577;0.997577;0.997577;0.996294;0.996294;0.996294 |
| m54088_170216_085445_30146696_ccs_Otu00092_21_1588_bp | 94.9 | Eukaryota;SAR;Stramenopiles;MAST_12;MAST_12C | 1;1;1;1;0.978928 |
| m54088_170216_085445_30146982_ccs_Otu00514_2_1588_bp | 96.4 | Eukaryota;Opisthokonta;Holozoa;Metazoa;Animalia;Eumetazoa;Bilateria;Platyhelminthes;Rhabditophora;Rhabdocoela;Neodalyellida | 0.999979;0.999979;0.999979;0.999979;0.999979;0.999979;0.999979;0.999979;0.988536;0.988536;0.988536 |
| m54088_170216_085445_30147354_ccs_Otu00385_3_1588_bp | 82.3 | Eukaryota;Opisthokonta;Nucletmycea;Fungi;Zoopagomycota | 0.997267;0.995993;0.995993;0.995993;0.995993 |
| m54088_170216_085445_30278563_ccs_Otu00290_5_1588_bp | 93.2 | Eukaryota;Opisthokonta;Holozoa;Metazoa;Animalia;Eumetazoa;Bilateria;Arthropoda;Hexapoda;Ellipura;Collembola | 0.999999;0.999999;0.999999;0.999999;0.999999;0.999999;0.999999;0.999999;0.861841;0.861841;0.861841 |
| m54088_170216_085445_30343509_ccs_Otu00488_2_1588_bp | 84.2 | Eukaryota;Opisthokonta;Holozoa;Metazoa;Animalia;Eumetazoa;Bilateria;Nematoda;Chromadorea;Tylenchida | 1;1;1;1;1;1;1;1;1;0.774694 |
| m54088_170216_085445_30343794_ccs_Otu00243_6_1588_bp | 80.3 | Eukaryota;Excavata;Discoba;Discicristata;Euglenozoa;Euglenida;Heteronematina;Petalomonas | 0.999998;0.999998;0.999998;0.999998;0.999998;0.887446;0.887446;0.887446 |
| m54088_170216_085445_30344074_ccs_Otu00336_4_1588_bp | 84.7 | Eukaryota;Opisthokonta;Holozoa;Metazoa;Animalia;Eumetazoa;Bilateria;Nematoda;Chromadorea;Tylenchida | 1;1;1;1;1;1;1;1;1;0.810225 |
| m54088_170216_085445_30539924_ccs_Otu00074_27_1588_bp | 94 | Eukaryota;SAR;Rhizaria;Cercozoa;Cercomonadidae | 0.99321;0.99321;0.99321;0.99321;0.99321 |
| m54088_170216_085445_30802413_ccs_Otu00129_14_1588_bp | 99.8 | Eukaryota;Archaeplastida;Chloroplastida;Chlorophyta;Trebouxiophyceae;Chloroidium | 1;1;1;1;1;0.999731 |
| m54088_170216_085445_31064386_ccs_Otu00432_3_1588_bp | 86.8 | Eukaryota;SAR;Rhizaria;Cercozoa;Vampyrellidae | 1;1;1;1;1 |
| m54088_170216_085445_31195377_ccs_Otu00307_4_1588_bp | 83.2 | Eukaryota;SAR;Stramenopiles;Labyrinthulomycetes;Sorodiplophrys | 0.99929;0.99929;0.99929;0.99929;0.99929 |
| m54088_170216_085445_31392396_ccs_Otu00120_16_1588_bp | 95.4 | Eukaryota;Amoebozoa;Tubulinea;Euamoebida;BOLA868 | 1;1;1;1;0.999858 |
| m54088_170216_085445_31457582_ccs_Otu00548_2_1588_bp | 95.9 | Eukaryota;Opisthokonta;Nucletmycea;Fungi;Dikarya;Ascomycota;Pezizomycotina;Pezizomycetes;Pezizales;Pezizaceae | 0.998378;0.998378;0.998378;0.998378;0.998378;0.998378;0.998378;0.998378;0.998378;0.998378 |
| m54088_170216_085445_32244194_ccs_Otu00646_2_1588_bp | 79 | Eukaryota;Opisthokonta;Nucletmycea;Fungi;Zoopagomycota;Incertae_Sedis | 0.990557;0.990557;0.990557;0.990557;0.990557;0.582835 |
| m54088_170216_085445_32309981_ccs_Otu00450_3_1588_bp | 99 | Eukaryota;Opisthokonta;Nucletmycea;Fungi;Chytridiomycota;Incertae_Sedis;Chytridiomycetes;Incertae_Sedis;Incertae_Sedis | 0.999999;0.999999;0.999999;0.999999;0.999999;0.999999;0.999999;0.967526;0.967526 |
| m54088_170216_085445_32374926_ccs_Otu00637_2_1588_bp | 77.8 | Eukaryota;Opisthokonta;Nucletmycea;Fungi;Dikarya;Ascomycota;Pezizomycotina;Sordariomycetes;Incertae_Sedis;Incertae_Sedis;Atractospora;decumbens | 1;1;1;1;1;1;1;1;1;1;0.812459;0.812459 |
| m54088_170216_085445_32571849_ccs_Otu00259_6_1588_bp | 98.6 | Eukaryota;Opisthokonta;Holozoa;Metazoa;Animalia;Eumetazoa;Bilateria;Nematoda;Chromadorea;Araeolaimida;Anaplectus;porosus | 0.999916;0.999916;0.999916;0.999916;0.999916;0.999916;0.999916;0.999916;0.999916;0.999916;0.999916;0.995863 |
| m54088_170216_085445_32899807_ccs_Otu00533_2_1588_bp | 75.3 | Eukaryota;SAR;Alveolata;Apicomplexa | |
| m54088_170216_085445_33358557_ccs_Otu00645_2_1588_bp | 84.1 | Eukaryota;Opisthokonta;Holozoa;Metazoa;Animalia;Eumetazoa;Bilateria;Nematoda;Chromadorea;Tylenchida | 1;1;1;1;1;1;1;1;1;0.827461 |
| m54088_170216_085445_33358660_ccs_Otu00357_4_1588_bp | 97.9 | Eukaryota;Opisthokonta;Nucletmycea;Fungi;Dikarya;Ascomycota;Pezizomycotina;Sordariomycetes;Hypocreales | |
| m54088_170216_085445_33751449_ccs_Otu00571_2_1588_bp | 91.1 | Eukaryota;SAR;Rhizaria;Cercozoa | 1;1;1;1 |
| m54088_170216_085445_33751827_ccs_Otu00421_3_1588_bp | 98.5 | Eukaryota;Archaeplastida;Chloroplastida;Chlorophyta;Chlorophyceae;Chlamydomonadales;Chlorococcum | 0.999999;0.999999;0.999999;0.999999;0.999999;0.999999;0.999576 |
| m54088_170216_085445_33752006_ccs_Otu00641_2_1588_bp | 97.1 | Eukaryota;Opisthokonta;Holozoa;Metazoa;Animalia;Eumetazoa;Bilateria;Nematoda;Chromadorea;Tylenchida;Paratylenchus | 1;1;1;1;1;1;1;1;1;1;0.999527 |
| m54088_170216_085445_33882525_ccs_Otu00500_2_1588_bp | 99 | Eukaryota;Archaeplastida;Chloroplastida;Charophyta;Phragmoplastophyta;Streptophyta;Embryophyta;Tracheophyta;Spermatophyta;Magnoliophyta;Asterales;Diplostephium | 0.793436;0.793436;0.793436;0.793436;0.793436;0.793436;0.793436;0.793436;0.793436;0.793436;0.77061;0.689461 |
| m54088_170216_085445_34210541_ccs_Otu00095_21_1588_bp | 95.7 | Eukaryota;Opisthokonta;Nucletmycea;Fungi;Dikarya;Ascomycota;Pezizomycotina;Sordariomycetes;Hypocreales | 0.967856;0.967856;0.967856;0.967856;0.967856;0.967856;0.967856;0.967856;0.967856 |
| m54088_170216_085445_34538072_ccs_Otu00493_2_1588_bp | 79.5 | Eukaryota;Opisthokonta;Nucletmycea;Fungi;Zoopagomycota | 0.999962;0.995533;0.995533;0.995533;0.995533 |
| m54088_170216_085445_34799849_ccs_Otu00581_2_1588_bp | 73.8 | Eukaryota;Amoebozoa | 0.998504;0.998504 |
| m54088_170216_085445_34931584_ccs_Otu00532_2_1588_bp | 89.3 | Eukaryota;SAR;Alveolata;Apicomplexa;Conoidasida;Gregarinasina;Eugregarinorida;Syncystis;mirabilis | 1;1;1;1;1;1;0.703311;0.703311;0.703311 |
| m54088_170216_085445_35586178_ccs_Otu00644_2_1588_bp | 95.7 | Eukaryota;SAR;Rhizaria;Cercozoa;Glissomonadida;Heteromita | 1;1;1;1;1;1 |
| m54088_170216_085445_35586753_ccs_Otu00086_23_1588_bp | 94.7 | Eukaryota;SAR;Rhizaria;Cercozoa;Vampyrellidae;Arachnula;impatiens | 0.999374;0.999374;0.999374;0.999374;0.999374;0.516543;0.516543 |
| m54088_170216_085445_35651729_ccs_Otu00550_2_1588_bp | 88.9 | Eukaryota;SAR;Rhizaria;Cercozoa | 0.992751;0.992751;0.992751;0.992751 |
| m54088_170216_085445_35652237_ccs_Otu00212_8_1588_bp | 78.2 | Eukaryota;Opisthokonta;Nucletmycea;Fungi;Zoopagomycota | 0.984643;0.984643;0.984643;0.984643;0.984643 |
| m54088_170216_085445_36438951_ccs_Otu00351_4_1588_bp | 98.9 | Eukaryota;Opisthokonta;Nucletmycea;Fungi;Dikarya;Ascomycota;Pezizomycotina;Eurotiomycetes;Onygenales;Onygenaceae;Auxarthron | 0.996126;0.996126;0.996126;0.996126;0.996126;0.996126;0.996126;0.996126;0.996126;0.996126;0.996126 |
| m54088_170216_085445_36963277_ccs_Otu00379_3_1588_bp | 78.9 | Eukaryota;Opisthokonta;Nucletmycea | 0.995297;0.995297;0.995297 |
| m54088_170216_085445_37552392_ccs_Otu00590_2_1588_bp | 98.5 | Eukaryota;SAR;Alveolata;Ciliophora;Intramacronucleata;Spirotrichea;Hypotrichia;Holosticha | 1;1;1;1;1;1;1;0.732948 |

| query | % similarity | annotation | confidence for each rank |
|---|---|---|---|
| m54088_170216_085445_37618562_ccs_Otu00406_3_1588_bp | 88.2 | Eukaryota;Opisthokonta;Nucletmycea;Fungi;Cryptomycota | 0.999925;0.999925;0.999925;0.999925;0.999925 |
| m54088_170216_085445_37748960_ccs_Otu00287_5_1588_bp | 99.4 | Eukaryota;Opisthokonta;Aphelidea | 1;1;1 |
| m54088_170216_085445_37814414_ccs_Otu00517_2_1588_bp | 97.3 | Eukaryota;Opisthokonta;Nucletmycea;Fungi;Dikarya;Ascomycota;Pezizomycotina;Dothideomycetes;Pleosporales | 1;1;1;1;1;1;1;1;1 |
| m54088_170216_085445_37946095_ccs_Otu00606_2_1588_bp | 90 | Eukaryota;Opisthokonta;Aphelidea;Paraphelidium | 1;1;1;0.850218 |
| m54088_170216_085445_38207693_ccs_Otu00217_8_1588_bp | 96.5 | Eukaryota;Opisthokonta;Holozoa;Metazoa;Animalia;Eumetazoa;Bilateria;Nematoda;Chromadorea;Monhysterida;Eumonhystera | 1;1;1;1;1;1;1;1;1;1;1 |
| m54088_170216_085445_38273143_ccs_Otu00633_2_1588_bp | 99.3 | Eukaryota;Opisthokonta;Nucletmycea;Fungi;Dikarya;Ascomycota;Pezizomycotina;Pezizomycetes;Pezizales;Pyronemataceae;Pyronema;domesticum | 0.998995;0.998995;0.998995;0.998995;0.998995;0.998995;0.998995;0.998995;0.998995;0.998995;0.998995;0.977758 |
| m54088_170216_085445_38339149_ccs_Otu00078_24_1588_bp | 87.4 | Eukaryota;SAR;Rhizaria;Cercozoa | 0.989892;0.989892;0.989892;0.989892 |
| m54088_170216_085445_38536093_ccs_Otu00174_10_1588_bp | 95.3 | Eukaryota;SAR;Stramenopiles;Labyrinthulomycetes | 0.998649;0.998649;0.998649;0.998649 |
| m54088_170216_085445_38666371_ccs_Otu00309_4_1588_bp | 87.3 | Eukaryota;Opisthokonta;Nucletmycea;Fungi;Cryptomycota;Incertae_Sedis;Incertae_Sedis;Incertae_Sedis;Incertae_Sedis;Paramicrosporidium | 1;1;1;1;1;0.574425;0.574425;0.574425;0.574425;0.574425 |
| m54088_170216_085445_38798233_ccs_Otu00600_2_1588_bp | 98.4 | Eukaryota;Archaeplastida;Chloroplastida;Chlorophyta;Trebouxiophyceae;Prasiolales | 0.982815;0.982815;0.982815;0.982815;0.982815;0.982815 |
| m54088_170216_085445_38863687_ccs_Otu00222_7_1588_bp | 96.9 | Eukaryota;Amoebozoa;Tubulinea;Euamoebida;BOLA868 | 0.992254;0.992254;0.992254;0.992254;0.992254 |
| m54088_170216_085445_39059933_ccs_Otu00559_2_1588_bp | 89 | Eukaryota;SAR;Alveolata;Ciliophora;Intramacronucleata;Conthreep;Colpodea | 0.987444;0.987444;0.987444;0.987444;0.987444;0.987444;0.773764 |
| m54088_170216_085445_39584092_ccs_Otu00447_3_1588_bp | 92.4 | Eukaryota;Amoebozoa;Discosea;Flabellinia;Vannellida | 0.983133;0.983133;0.983133;0.983133;0.983133 |
| m54088_170216_085445_39781056_ccs_Otu00321_4_1588_bp | 94.3 | Eukaryota;Opisthokonta;Nucletmycea;Fungi;Chytridiomycota;Incertae_Sedis;Chytridiomycetes;Spizellomycetales;Powellomycetaceae | 0.983056;0.983056;0.983056;0.983056;0.983056;0.983056;0.983056;0.983056;0.983056 |
| m54088_170216_085445_39977600_ccs_Otu00516_2_1588_bp | 93.4 | Eukaryota;SAR;Rhizaria;Cercozoa;Phytomyxea | 1;1;1;1;1 |
| m54088_170216_085445_40043093_ccs_Otu00627_2_1588_bp | 88.1 | Eukaryota;SAR;Stramenopiles;Peronosporomycetes | 0.999999;0.999999;0.999999;0.99995 |
| m54088_170216_085445_40305065_ccs_Otu00383_3_1588_bp | 97.7 | Eukaryota;Opisthokonta;Holozoa;Metazoa;Animalia;Eumetazoa;Bilateria;Nematoda;Enoplea;Enoplia;Triplonchida | 0.999999;0.999999;0.999999;0.999999;0.999999;0.999999;0.999999;0.999999;0.999999;0.999999;0.999999 |
| m54088_170216_085445_40567609_ccs_Otu00528_2_1588_bp | 99.4 | Eukaryota;Archaeplastida;Chloroplastida;Chlorophyta;Chlorophyceae;Chlorosarcinales;Neochlorosarcina;negevensis | 1;1;1;1;1;0.998288;0.998288;0.998288 |
| m54088_170216_085445_40960080_ccs_Otu00417_3_1588_bp | 99.8 | Eukaryota;Archaeplastida;Chloroplastida;Chlorophyta;Trebouxiophyceae;Ctenocladales;Leptosira | 1;1;1;1;0.962903;0.962903;0.962903 |
| m54088_170216_085445_41026276_ccs_Otu00378_3_1588_bp | 75.4 | Eukaryota;Opisthokonta;Nucletmycea | 0.988582;0.988582;0.988582 |
| m54088_170216_085445_41222300_ccs_Otu00246_6_1588_bp | 91.4 | Eukaryota;Opisthokonta;Nucletmycea;Fungi;Cryptomycota;LKM11 | 0.996098;0.996098;0.996098;0.996098;0.996098;0.996098 |
| m54088_170216_085445_41484748_ccs_Otu00582_2_1588_bp | 95.6 | Eukaryota;SAR;Alveolata;Ciliophora;Intramacronucleata;Conthreep;Colpodea;Colpodida | 1;1;1;1;1;1;1;0.997604 |
| m54088_170216_085445_41550544_ccs_Otu00163_11_1588_bp | 95.8 | Eukaryota;Opisthokonta;Holozoa;Metazoa;Animalia;Eumetazoa;Bilateria;Nematoda;Chromadorea;Rhabditida;Mesorhabditis | 1;1;1;1;1;1;1;1;1;1;0.982772 |
| m54088_170216_085445_41878422_ccs_Otu00414_3_1588_bp | 80.3 | Eukaryota;SAR;Stramenopiles;Labyrinthulomycetes;Sorodiplophrys | 1;1;1;1;0.627976 |
| m54088_170216_085445_42074584_ccs_Otu00435_3_1588_bp | 78.9 | Eukaryota;Opisthokonta;Nucletmycea;Fungi;Zoopagomycota | 0.992212;0.989833;0.989833;0.989833;0.989833 |
| m54088_170216_085445_42140250_ccs_Otu00338_4_1588_bp | 95.7 | Eukaryota;Amoebozoa;Tubulinea;Arcellinida;Echinamoebida | 1;1;1;1;1 |
| m54088_170216_085445_42336553_ccs_Otu00448_3_1588_bp | 97.7 | Eukaryota;SAR;Rhizaria;Cercozoa;Cercomonadidae | 1;1;1;1;1 |
| m54088_170216_085445_43254029_ccs_Otu00314_4_1588_bp | 97 | Eukaryota;SAR;Stramenopiles;Ochrophyta;Diatomea;Bacillariophytina;Bacillariophyceae;Pinnularia;nodosa | 0.999998;0.999998;0.999998;0.999998;0.999998;0.999998;0.999998;0.999998;0.991872 |
| m54088_170216_085445_43254285_ccs_Otu00359_3_1588_bp | 84.3 | Eukaryota;Opisthokonta;Holozoa;Metazoa;Animalia;Eumetazoa;Bilateria;Nematoda;Enoplea;Dorylaimia;Mermithida;Romanomermis;culicivorax | 1;1;1;1;1;1;1;1;1;1;0.558905;0.558905;0.558905 |
| m54088_170216_085445_43778558_ccs_Otu00341_4_1588_bp | 90.9 | Eukaryota;SAR;Rhizaria;Cercozoa;Cercomonadidae | 0.999965;0.999965;0.999965;0.999965;0.999965 |
| m54088_170216_085445_43778918_ccs_Otu00561_2_1588_bp | 95.5 | Eukaryota;Opisthokonta;Nucletmycea;Fungi;Cryptomycota;LKM11 | 1;1;1;1;1;0.545821 |
| m54088_170216_085445_43779019_ccs_Otu00481_2_1588_bp | 99.4 | Eukaryota;Opisthokonta;Nucletmycea;Fungi;Dikarya;Ascomycota;Pezizomycotina;Eurotiomycetes;Eurotiales;Trichocomaceae;Talaromyces | 0.999967;0.999967;0.999967;0.999967;0.999967;0.999967;0.999967;0.999967;0.999967;0.998744;0.998744 |
| m54088_170216_085445_43844301_ccs_Otu00420_3_1588_bp | 91.4 | Eukaryota;Opisthokonta;Nucletmycea;Fungi;Cryptomycota | 1;1;1;1;1 |
| m54088_170216_085445_43909533_ccs_Otu00472_2_1588_bp | 94.7 | Eukaryota;Opisthokonta;Nucletmycea;Fungi;Dikarya;Ascomycota;Pezizomycotina;Pezizomycetes;Pezizales;Ascobolaceae;Saccobolus;dilutellus | 1;1;1;1;1;1;1;1;1;0.738242;0.738242;0.738242 |
| m54088_170216_085445_44302653_ccs_Otu00597_2_1588_bp | 94.2 | Eukaryota;Opisthokonta;Nucletmycea;Fungi;Cryptomycota | 1;1;1;1;1 |
| m54088_170216_085445_44368744_ccs_Otu00496_2_1588_bp | 84.7 | Eukaryota;Opisthokonta;Nucletmycea;Fungi;Cryptomycota | 0.985757;0.985757;0.985757;0.985757;0.985757 |
| m54088_170216_085445_44761206_ccs_Otu00107_18_1588_bp | 94 | Eukaryota;Amoebozoa;Tubulinea;Euamoebida;BOLA868 | 1;1;1;1;0.999053 |
| m54088_170216_085445_45023512_ccs_Otu00598_2_1588_bp | 96 | Eukaryota;SAR;Alveolata;Ciliophora;Intramacronucleata;Spirotrichea;Hypotrichia | 1;1;1;1;1;1;1 |
| m54088_170216_085445_45089714_ccs_Otu00391_3_1588_bp | 96.6 | Eukaryota;Opisthokonta;Nucletmycea;Fungi;Dikarya;Ascomycota;Pezizomycotina;Leotiomycetes | |
| m54088_170216_085445_45220318_ccs_Otu00643_2_1588_bp | 72.2 | chimera | |
| m54088_170216_085445_45614013_ccs_Otu00478_2_1588_bp | 98.6 | Eukaryota;Opisthokonta;Nucletmycea;Fungi;Mucoromycota;Mortierellomycotina;Incertae_Sedis;Mortierellales;Mortierellaceae;Mortierella | 0.776843;0.776843;0.776843;0.776843;0.776843;0.776843;0.776843;0.776843;0.776843;0.558708 |
| m54088_170216_085445_45744837_ccs_Otu00491_2_1588_bp | 87.7 | Eukaryota;SAR;Rhizaria;Cercozoa | 0.723122;0.723122;0.723122;0.723122 |
| m54088_170216_085445_45940853_ccs_Otu00330_4_1588_bp | 90.4 | Eukaryota;SAR;Rhizaria;Cercozoa | 0.99997;0.99997;0.99997;0.99997 |
| m54088_170216_085445_46334487_ccs_Otu00545_2_1588_bp | 89.7 | Eukaryota;SAR;Alveolata;Apicomplexa;Conoidasida;Cryptosporida;Cryptosporidium;Colpodella;tetrahymenae | 1;1;1;1;1;1;1;1;0.512451 |
| m54088_170216_085445_46465519_ccs_Otu00189_9_1588_bp | 86.4 | Eukaryota;Opisthokonta;Nucletmycea;Fungi;Zoopagomycota | 0.999963;0.999963;0.999963;0.999963;0.999963 |
| m54088_170216_085445_46727794_ccs_Otu00026_107_1588_bp | 99.3 | Eukaryota;Opisthokonta;Nucletmycea;Fungi;Dikarya;Ascomycota;Pezizomycotina;Eurotiomycetes;Eurotiales;Aspergillaceae;Penicillium;capsulatum | 0.99999;0.99999;0.99999;0.99999;0.99999;0.99999;0.99999;0.99999;0.99999;0.99999;0.99737;0.99737 |
| m54088_170216_085445_46924579_ccs_Otu00601_2_1588_bp | 95.8 | Eukaryota;Amoebozoa;Schizoplasmodiida;Phalansterium | 1;1;1;1 |
| m54088_170216_085445_48169818_ccs_Otu00619_2_1588_bp | 95.7 | Eukaryota;Opisthokonta;Holozoa;Metazoa;Animalia;Eumetazoa;Bilateria;Nematoda;Chromadorea;Tylenchida;Nematoda | 0.999634;0.999634;0.999634;0.999634;0.999634;0.999634;0.999634;0.999634;0.999634;0.999634; |
| m54088_170216_085445_48300753_ccs_Otu00377_3_1588_bp | 96.6 | Eukaryota;Opisthokonta;Holozoa;Metazoa;Animalia;Eumetazoa;Bilateria;Arthropoda;Hexapoda;Ellipura;Collembola | 1;1;1;1;1;1;1;1;0.959903;0.959903;0.959903 |
| m54088_170216_085445_48562330_ccs_Otu00540_2_1588_bp | 68.6 | Eukaryota;Opisthokonta;Nucletmycea;Fungi;Dikarya;Basidiomycota;Agaricomycotina | 0.702486;0.702486;0.702486;0.702486;0.702486;0.702486;0.694199 |
| m54088_170216_085445_48890164_ccs_Otu00510_2_1588_bp | 93.8 | Eukaryota;Incertae_Sedis;Breviatea;Breviata;anathema | 1;1;1;0.842379;0.842379 |
| m54088_170216_085445_49742462_ccs_Otu00362_3_1588_bp | 95.3 | Eukaryota;Opisthokonta;Nucletmycea;Fungi;LKM15 | 1;1;1;1;1 |
| m54088_170216_085445_49742821_ccs_Otu00237_7_1588_bp | 99.4 | Eukaryota;Archaeplastida;Chloroplastida;Chlorophyta;Trebouxiophyceae;Spongiococcum;tetrasporum | 0.999999;0.999999;0.999999;0.999999;0.777661;0.777661;0.777661 |
| m54088_170216_085445_50070093_ccs_Otu00473_2_1588_bp | 99.6 | Eukaryota;Archaeplastida;Chloroplastida;Chlorophyta;Trebouxiophyceae;Incertae_Sedis;Coccomyxa;simplex | 1;1;1;1;0.999314;0.999314;0.999314;0.999314 |
| m54088_170216_085445_50266620_ccs_Otu00167_11_1588_bp | 99.1 | Eukaryota;Opisthokonta;Holozoa;Metazoa;Animalia;Eumetazoa;Bilateria;Nematoda;Enoplea;Dorylaimia;Dorylaimida | 0.979932;0.979932;0.979932;0.979932;0.979932;0.979932;0.979932;0.979932;0.979932;0.979932;0.979932 |
| m54088_170216_085445_50594320_ccs_Otu00269_6_1588_bp | 99.5 | Eukaryota;Opisthokonta;Nucletmycea;Fungi;Dikarya;Ascomycota;Pezizomycotina;Eurotiomycetes;Chaetothyriales;Herpotrichiellaceae | 1;1;1;1;1;1;1;1;0.999469;0.999469 |
| m54088_170216_085445_50725413_ccs_Otu00407_3_1588_bp | 92.1 | Eukaryota;SAR;Rhizaria;Cercozoa | 0.979862;0.979862;0.979862;0.979862 |
| m54088_170216_085445_50856891_ccs_Otu00364_3_1588_bp | 77.7 | Eukaryota;SAR;Alveolata;Apicomplexa | |
| m54088_170216_085445_50921618_ccs_Otu00374_3_1588_bp | 95.2 | Eukaryota;Amoebozoa;Tubulinea;Euamoebida;BOLA868 | 1;1;1;1;0.999711 |

| query | % similarity | annotation | confidence for each rank |
|---|---|---|---|
| m54088_170216_085445_50922389_ccs_Otu00203_8_1588_bp | 91.5 | Eukaryota;Opisthokonta;Nucletmycea;Fungi;Cryptomycota | 1;1;1;1;1 |
| m54088_170216_085445_51053085_ccs_Otu00376_3_1588_bp | 95.1 | Eukaryota;Amoebozoa;Tubulinea;Euamoebida;BOLA868 | 0.998831;0.998831;0.998831;0.998831;0.998335 |
| m54088_170216_085445_51250051_ccs_Otu00153_11_1588_bp | 97.4 | Eukaryota;SAR;Stramenopiles;Ochrophyta;Chrysophyceae;Ochromonadales;Ochromonas | 1;1;1;1;1;0.998596;0.998596 |
| m54088_170216_085445_52691947_ccs_Otu00207_8_1588_bp | 92.8 | Eukaryota;SAR;Rhizaria;Cercozoa;Imbricatea | 1;1;1;1;1 |
| m54088_170216_085445_52887729_ccs_Otu00558_2_1588_bp | 91.5 | Eukaryota;Opisthokonta;Holozoa;Metazoa;Animalia;Eumetazoa;Bilateria;Tardigrada;Eutardigrada;Parachela | 1;1;1;1;1;1;1;1;1;1 |
| m54088_170216_085445_53019444_ccs_Otu00470_2_1588_bp | 87.6 | Eukaryota;SAR;Alveolata;Apicomplexa;Conoidasida;Gregarinasina;Eugregarinorida;Syncystis;mirabilis | 1;1;1;1;1;1;0.649337;0.649337;0.649337 |
| m54088_170216_085445_53346699_ccs_Otu00484_2_1588_bp | 95.7 | Eukaryota;Opisthokonta;Nucletmycea;Fungi;Chytridiomycota;Incertae_Sedis;Chytridiomycetes | 1;1;1;1;1;1;1 |
| m54088_170216_085445_53608653_ccs_Otu00434_3_1588_bp | 90.3 | Eukaryota;Opisthokonta | 0.985635;0.985635 |
| m54088_170216_085445_53871035_ccs_Otu00471_2_1588_bp | 93.5 | Eukaryota;Opisthokonta;Nucletmycea;Fungi;Cryptomycota | 1;1;1;1;1 |
| m54088_170216_085445_53936589_ccs_Otu00340_4_1588_bp | 91.3 | Eukaryota;Opisthokonta;Nucletmycea;Fungi;Cryptomycota | 1;1;1;1;1 |
| m54088_170216_085445_53936950_ccs_Otu00103_18_1588_bp | 95.9 | Eukaryota;Amoebozoa;Tubulinea;Euamoebida;BOLA868 | 0.999968;0.999968;0.999968;0.999968;0.999346 |
| m54088_170216_085445_54526020_ccs_Otu00397_3_1588_bp | 86.8 | Eukaryota;Excavata;Discoba;Discicristata;Heterolobosea;Tetramitia;Vahlkampfia;inornata | 1;1;1;1;1;1;0.92022;0.92022 |
| m54088_170216_085445_54592263_ccs_Otu00609_2_1588_bp | 78.7 | Eukaryota;Opisthokonta;Nucletmycea;Fungi;Zoopagomycota | 0.984268;0.984268;0.984268;0.984268;0.984268 |
| m54088_170216_085445_54723061_ccs_Otu00257_6_1588_bp | 98.3 | Eukaryota;SAR;Rhizaria;Cercozoa;Cercomonadidae | 0.995696;0.995696;0.995696;0.995696;0.995696 |
| m54088_170216_085445_54853828_ccs_Otu00081_24_1588_bp | 98.2 | Eukaryota;SAR;Alveolata;Ciliophora;Intramacronucleata;Litostomatea;Haptoria;Pseudoholophrya;terricola | 1;1;1;1;1;1;1;0.699986;0.699986 |
| m54088_170216_085445_54854424_ccs_Otu00615_2_1588_bp | 98.7 | Eukaryota;SAR;Alveolata;Ciliophora;Intramacronucleata;Spirotrichea;Hypotrichia | 0.975526;0.975526;0.975526;0.975526;0.975526;0.975526;0.975526 |
| m54088_170216_085445_55181982_ccs_Otu00423_3_1588_bp | 98.1 | Eukaryota;SAR;Alveolata;Ciliophora;Intramacronucleata;Conthreep;Oligohymenophorea;Peritrichia | 1;1;1;1;1;1;1;1 |
| m54088_170216_085445_55247558_ccs_Otu00368_3_1588_bp | 96.3 | Eukaryota;SAR;Rhizaria;Cercozoa;Glissomonadida | 1;1;1;1;0.709328 |
| m54088_170216_085445_55312730_ccs_Otu00549_2_1588_bp | 93.5 | Eukaryota;Opisthokonta;Nucletmycea;Fungi;Chytridiomycota;Incertae_Sedis;Chytridiomycetes;Rhizophydiales;Rhizophydiaceae;Rhizophydium | 1;1;1;1;1;1;1;1;1;1 |
| m54088_170216_085445_55313015_ccs_Otu00604_2_1588_bp | 86.6 | Eukaryota;Opisthokonta;Nucletmycea;Fungi;Cryptomycota | 0.999441;0.999441;0.999441;0.999441;0.999441 |
| m54088_170216_085445_55640456_ccs_Otu00313_4_1588_bp | 90.6 | Eukaryota;SAR;Alveolata;Ciliophora;Intramacronucleata;Conthreep;Oligohymenophorea | 0.987773;0.987773;0.987773;0.987773;0.987773;0.987773;0.987773 |
| m54088_170216_085445_55968048_ccs_Otu00247_6_1588_bp | 88.6 | Eukaryota;SAR;Alveolata;Apicomplexa;Conoidasida;Gregarinasina;Eugregarinorida;Syncystis;mirabilis | 1;1;1;1;1;1;0.646998;0.646998;0.646998 |
| m54088_170216_085445_56033960_ccs_Otu00319_4_1588_bp | 98.8 | Eukaryota;SAR;Stramenopiles;Peronosporomycetes | 0.989247;0.989247;0.989247;0.989247 |
| m54088_170216_085445_56099267_ccs_Otu00161_11_1588_bp | 97.8 | Eukaryota;Opisthokonta;Nucletmycea;Fungi;Dikarya;Basidiomycota;Agaricomycotina;Agaricomycetes;Trechisporales;Hydnodontaceae;Trechispora | 1;1;1;1;1;1;1;1;0.999793;0.999793;0.999793 |
| m54088_170216_085445_56426936_ccs_Otu00073_27_1588_bp | 98.5 | Eukaryota;Opisthokonta;Nucletmycea;Fungi;Cryptomycota;LKM11 | 1;1;1;1;1;1 |
| m54088_170216_085445_56754609_ccs_Otu00398_3_1588_bp | 97.4 | Eukaryota;SAR;Alveolata;Ciliophora;Intramacronucleata;Spirotrichea;Hypotrichia | 1;1;1;1;1;1;1 |
| m54088_170216_085445_56754766_ccs_Otu00433_3_1588_bp | 96.8 | Eukaryota;Opisthokonta;Nucletmycea;Fungi;Zoopagomycota;Zoopagomycotina;Incertae_Sedis;Zoopagales;Piptocephalidaceae;Piptocephalis;xenophila | 1;1;1;1;1;1;1;1;0.672769;0.672769;0.672769 |
| m54088_170216_085445_56886113_ccs_Otu00273_6_1588_bp | 96 | Eukaryota;Opisthokonta;Holozoa;Metazoa;Animalia;Eumetazoa;Bilateria;Nematoda;Chromadorea;Tylenchida;Pratylenchus;fallax | 0.999999;0.999999;0.999999;0.999999;0.999999;0.999999;0.999999;0.999999;0.999999;0.999999;0.999999;0.827172 |
| m54088_170216_085445_57213530_ccs_Otu00191_9_1588_bp | 98.9 | Eukaryota;Opisthokonta;Nucletmycea;Fungi;Dikarya;Basidiomycota;Agaricomycotina;Agaricomycetes | 0.996518;0.996518;0.996518;0.996518;0.996518;0.996518;0.996518;0.996518 |
| m54088_170216_085445_57540917_ccs_Otu00518_2_1588_bp | 96.8 | Eukaryota;SAR;Alveolata;Ciliophora;Intramacronucleata;Spirotrichea;Hypotrichia | 0.998647;0.998647;0.998647;0.998647;0.998647;0.998647;0.998647 |
| m54088_170216_085445_57606393_ccs_Otu00642_2_1588_bp | 75.6 | Eukaryota;Opisthokonta;Nucletmycea;Fungi;Dikarya;Ascomycota;Pezizomycotina;Sordariomycetes | 0.99602;0.99602;0.99602;0.99602;0.99602;0.99602;0.99602;0.99602 |
| m54088_170216_085445_57803402_ccs_Otu00014_265_1588_bp | 99.4 | Eukaryota;SAR;Alveolata;Ciliophora;Intramacronucleata;Spirotrichea;Hypotrichia | 0.968775;0.968775;0.968775;0.968775;0.968775;0.968775;0.968775 |
| m54088_170216_085445_57868929_ccs_Otu00602_2_1588_bp | 96.5 | Eukaryota;Opisthokonta;Holozoa;Metazoa;Animalia;Eumetazoa;Bilateria;Tardigrada;Eutardigrada;Parachela | 1;1;1;1;1;1;1;1;1;1 |
| m54088_170216_085445_57999443_ccs_Otu00544_2_1588_bp | 90.2 | Eukaryota;Opisthokonta;Aphelidea;Paraphelidium | 1;1;1;0.884395 |
| m54088_170216_085445_58327852_ccs_Otu00187_9_1588_bp | 98.2 | Eukaryota;Opisthokonta;Nucletmycea;Fungi;Dikarya;Ascomycota;Pezizomycotina;Sordariomycetes;Hypocreales | 1;1;1;1;1;1;1;1;1 |
| m54088_170216_085445_58852084_ccs_Otu00486_2_1588_bp | 91.3 | Eukaryota;Opisthokonta;Holozoa;Metazoa;Animalia;Eumetazoa;Bilateria;Arthropoda;Crustacea;Maxillopoda;Copepoda | 0.999224;0.999224;0.999224;0.999224;0.999224;0.999224;0.999224;0.999224;0.998842;0.998842;0.998842 |
| m54088_170216_085445_58983335_ccs_Otu00596_2_1588_bp | 75.8 | Eukaryota;SAR;Alveolata;Apicomplexa | |
| m54088_170216_085445_59048878_ccs_Otu00386_3_1588_bp | 98.7 | Eukaryota;Opisthokonta;Nucletmycea;Fungi;Zoopagomycota;Kickxellomycotina;Incertae_Sedis;Kickxellales;Kickxellaceae;Coemansia | 1;1;1;1;1;0.95191;0.95191;0.95191;0.95191;0.95191 |
| m54088_170216_085445_59179415_ccs_Otu00354_4_1588_bp | 98.3 | Eukaryota;Opisthokonta;Holozoa;Metazoa;Animalia;Eumetazoa;Bilateria;Nematoda;Chromadorea;Tylenchida;Nematoda | 0.987059;0.987059;0.987059;0.987059;0.987059;0.987059;0.987059;0.987059;0.987059;0.987059; |
| m54088_170216_085445_59507136_ccs_Otu00347_4_1588_bp | 91.1 | Eukaryota;Opisthokonta;Nucletmycea;Fungi;Cryptomycota | 1;1;1;1;1 |
| m54088_170216_085445_59703894_ccs_Otu00264_6_1588_bp | 93 | Eukaryota;Opisthokonta;Nucletmycea;Fungi;Dikarya;Basidiomycota;Pucciniomycotina;Agaricostilbomycetes;Agaricostilbales;Chionosphaeraceae | 1;1;1;1;1;1;1;1;1;0.59842 |
| m54088_170216_085445_60228110_ccs_Otu00399_3_1588_bp | 90.5 | Eukaryota;Opisthokonta;Nucletmycea;Fungi;Cryptomycota | 1;1;1;1;1 |
| m54088_170216_085445_60293256_ccs_Otu00349_4_1588_bp | 98.2 | Eukaryota;Opisthokonta;Holozoa;Metazoa;Animalia;Eumetazoa;Bilateria;Arthropoda;Chelicerata;Arachnida;Acari;Tectocepheus;sarekensis | 1;1;1;1;1;1;1;1;1;1;1;0.998175;0.998175 |
| m54088_170216_085445_60817557_ccs_Otu00610_2_1588_bp | 81.4 | Eukaryota;Opisthokonta;Holozoa;Metazoa;Animalia;Eumetazoa;Bilateria;Nematoda;Chromadorea | 0.974036;0.974036;0.974036;0.974036;0.974036;0.974036;0.974036;0.974036;0.974036 |
| m54088_170216_085445_60883529_ccs_Otu00630_2_1588_bp | 96.2 | Eukaryota;Opisthokonta;Holozoa;Metazoa;Animalia;Eumetazoa;Bilateria;Nematoda;Enoplea;Enoplia;Enoplida | 0.99997;0.99997;0.99997;0.99997;0.99997;0.99997;0.99997;0.99997;0.99997;0.99997;0.99997 |
| m54088_170216_085445_61472932_ccs_Otu00546_2_1588_bp | 90.7 | Eukaryota;SAR;Rhizaria;Cercozoa;Vampyrellidae | 0.999997;0.999997;0.999997;0.999997;0.999997 |
| m54088_170216_085445_61800849_ccs_Otu00075_25_1588_bp | 98.7 | Eukaryota;SAR;Stramenopiles;Peronosporomycetes;Pythium | 0.999983;0.999983;0.999983;0.999983;0.998893 |
| m54088_170216_085445_61932130_ccs_Otu00211_8_1588_bp | 99.1 | Eukaryota;SAR;Rhizaria;Cercozoa;Cercomonadidae | 1;1;1;1;1 |
| m54088_170216_085445_62390461_ccs_Otu00509_2_1588_bp | 95 | Eukaryota;Opisthokonta;Nucletmycea;Fungi;Dikarya;Basidiomycota;Agaricomycotina;Agaricomycetes;Agaricales;Tricholomataceae;Camarophyllopsis | 0.999756;0.999756;0.999756;0.999756;0.999756;0.999756;0.999756;0.999756;0.999756;0.999756;0.999756 |
| m54088_170216_085445_62390477_ccs_Otu00263_6_1588_bp | 98.1 | Eukaryota;Opisthokonta;Nucletmycea;Fungi;Dikarya;Ascomycota;Pezizomycotina;Sordariomycetes | 1;1;1;1;1;1;1;1 |
| m54088_170216_085445_62653370_ccs_Otu00552_2_1588_bp | 83.1 | Eukaryota;Opisthokonta;Nucletmycea | 0.999681;0.999681;0.999681 |
| m54088_170216_085445_62784202_ccs_Otu00274_6_1588_bp | 97.2 | Eukaryota;Opisthokonta;Holozoa;Metazoa;Animalia;Eumetazoa;Bilateria;Annelida;Clitellata;Oligochaeta;Haplotaxida;Achaeta;affinis | 0.999252;0.999252;0.999252;0.999252;0.999252;0.999252;0.999252;0.999252;0.999252;0.999252;0.999252;0.999252;0.6659 |
| m54088_170216_085445_62849868_ccs_Otu00419_3_1588_bp | 97.8 | Eukaryota;Opisthokonta;Nucletmycea;Fungi;Dikarya;Basidiomycota;Agaricomycotina;Agaricomycetes | 1;1;1;1;1;1;1;1 |
| m54088_170216_085445_6291739_ccs_Otu00494_2_1588_bp | 87.2 | Eukaryota;SAR;Alveolata;Apicomplexa;Conoidasida;Gregarinasina;Archigregarinorida;Selenidium | 1;1;1;1;1;1;1;1 |
| m54088_170216_085445_63046112_ccs_Otu00625_2_1588_bp | 91.5 | Eukaryota;SAR;Rhizaria;Cercozoa;Vampyrellidae | 0.999996;0.999996;0.999996;0.999996;0.999996 |
| m54088_170216_085445_63373468_ccs_Otu00503_2_1588_bp | 89.7 | Eukaryota;Opisthokonta;Holozoa;Metazoa;Animalia;Eumetazoa;Bilateria;Arthropoda;Chelicerata;Arachnida;Acari | 0.993436;0.993436;0.993436;0.993436;0.993436;0.993436;0.993436;0.993436;0.993436;0.993436;0.993436 |
| m54088_170216_085445_64094469_ccs_Otu00623_2_1588_bp | 84.1 | Eukaryota;Opisthokonta;Holozoa;Metazoa;Animalia;Eumetazoa;Bilateria;Nematoda;Enoplea;Dorylaimia | 1;1;1;1;1;1;1;1;1;1 |
| m54088_170216_085445_64095031_ccs_Otu00392_3_1588_bp | 90.7 | Eukaryota;SAR;Rhizaria;Cercozoa | 0.996715;0.996715;0.996715;0.996715 |
| m54088_170216_085445_64160450_ccs_Otu00636_2_1588_bp | 94.7 | Eukaryota;Opisthokonta;Holozoa;Metazoa;Animalia;Eumetazoa;Bilateria;Nematoda;Enoplea;Enoplia;Enoplida | 0.999999;0.999999;0.999999;0.999999;0.999999;0.999999;0.999999;0.999999;0.999999;0.999999;0.999999 |

| query | % similarity | annotation | confidence for each rank |
|---|---|---|---|
| m54088_170216_085445_64618868_ccs_Otu00599_2_1588_bp | 83.5 | Eukaryota;Opisthokonta;Nucletmycea;Fungi;Dikarya;Ascomycota;Pezizomycotina;Sordariomycetes;Hypocreales;Bionectriaceae;Clonostachys;rosea | 0.985696;0.985696;0.985696;0.985696;0.985696;0.985696;0.985696;0.985696;0.985696;0.985696;0.985696;0.965313 |
| m54088_170216_085445_64946254_ccs_Otu00279_5_1588_bp | 98.6 | Eukaryota;Opisthokonta;Nucletmycea;Fungi;Dikarya;Ascomycota;Pezizomycotina;Sordariomycetes;Hypocreales;Bionectriaceae;Clonostachys | 0.992078;0.992078;0.992078;0.992078;0.992078;0.992078;0.992078;0.992078;0.992078;0.786682;0.786682 |
| m54088_170216_085445_65274564_ccs_Otu00138_13_1588_bp | 97.4 | Eukaryota;Opisthokonta;Nucletmycea;Fungi;Dikarya;Ascomycota;Pezizomycotina;Dothideomycetes;Pleosporales | 1;1;1;1;1;1;1;1;1 |
| m54088_170216_085445_65340055_ccs_Otu00490_2_1588_bp | 93.3 | Eukaryota;Opisthokonta;Aphelidea;Paraphelidium | 1;1;1;0.956161 |
| m54088_170216_085445_65470808_ccs_Otu00400_3_1588_bp | 95.6 | Eukaryota;SAR;Rhizaria;Cercozoa;Glissomonadida;Heteromita | 0.998587;0.998587;0.998587;0.998587;0.998587;0.998587 |
| m54088_170216_085445_66322750_ccs_Otu00064_31_1588_bp | 91 | Eukaryota;Opisthokonta;Nucletmycea;Fungi;Chytridiomycota;Incertae_Sedis;Chytridiomycetes;Rhizophydiales;Rhizophydiaceae;Rhizophydium;patellarium | 1;1;1;1;1;1;1;1;1;0.828318;0.828318 |
| m54088_170216_085445_66585512_ccs_Otu00289_5_1588_bp | 99 | Eukaryota;Opisthokonta;Nucletmycea;Fungi;Dikarya;Basidiomycota;Agaricomycotina;Agaricomycetes;Cantharellales;Ceratobasidiaceae;Ceratobasidium | 0.999998;0.999998;0.999998;0.999998;0.999998;0.999998;0.999998;0.999998;0.884242;0.884242;0.884242 |
| m54088_170216_085445_66913096_ccs_Otu00160_11_1588_bp | 95.2 | Eukaryota;Opisthokonta;Nucletmycea;Fungi;Zoopagomycota;Zoopagomycotina;Incertae_Sedis;Zoopagales;Piptocephalidaceae;Piptocephalis;xenophila | 1;1;1;1;1;1;1;1;0.711049;0.711049;0.711049 |
| m54088_170216_085445_67043611_ccs_Otu00413_3_1588_bp | 72 | Eukaryota;Opisthokonta;Nucletmycea;Fungi;Dikarya;Basidiomycota;Agaricomycotina;Agaricomycetes;Agaricales;Amanitaceae;Amanita;bisporigera | 0.951459;0.951459;0.951459;0.951459;0.951459;0.951459;0.72917;0.708705;0.708705;0.53234;0.53234;0.53234 |
| m54088_170216_085445_67305804_ccs_Otu00050_45_1588_bp | 97.1 | Eukaryota;SAR;Alveolata;Ciliophora;Intramacronucleata;Conthreep;Colpodea;Colpodida | 0.999751;0.999751;0.999751;0.999751;0.999751;0.999751;0.999751;0.999751 |
| m54088_170216_085445_67503046_ccs_Otu00501_2_1588_bp | 98.7 | Eukaryota;Archaeplastida;Chloroplastida;Chlorophyta;Trebouxiophyceae | 0.999999;0.999999;0.999999;0.999999;0.999999 |
| m54088_170216_085445_67503068_ccs_Otu00411_3_1588_bp | 81.5 | Eukaryota;Opisthokonta;Nucletmycea;Fungi;Cryptomycota | 0.998118;0.998118;0.998118;0.998118;0.998118 |
| m54088_170216_085445_67896206_ccs_Otu00457_3_1588_bp | 89.8 | Eukaryota;Opisthokonta;Nucletmycea;Fungi;LKM15 | 1;1;1;1;1 |
| m54088_170216_085445_68747772_ccs_Otu00230_7_1588_bp | 76.7 | Eukaryota;SAR;Alveolata;Apicomplexa | |
| m54088_170216_085445_68813311_ccs_Otu00522_2_1588_bp | 95.9 | Eukaryota;Opisthokonta;Nucletmycea;Fungi;Dikarya;Ascomycota;Saccharomycotina;Saccharomycetes;Saccharomycetales;Incertae_Sedis;Middelhovenomyces | 0.985092;0.985092;0.985092;0.985092;0.985092;0.985092;0.985092;0.985092;0.985092;0.885645;0.885645 |
| m54088_170216_085445_69009586_ccs_Otu00363_3_1588_bp | 89.7 | Eukaryota;Opisthokonta;Nucletmycea;Fungi;Zoopagomycota;Zoopagomycotina;Incertae_Sedis;Zoopagales | 1;1;1;1;1;1;1;1 |
| m54088_170216_085445_69206648_ccs_Otu00366_3_1588_bp | 81.9 | Eukaryota;SAR;Alveolata | 0.745146;0.745146;0.745146 |
| m54088_170216_085445_69206793_ccs_Otu00245_6_1588_bp | 98.7 | Eukaryota;Opisthokonta;Nucletmycea;Fungi;Dikarya;Ascomycota;Pezizomycotina;Dothideomycetes;Pleosporales | 1;1;1;1;1;1;1;1;1 |
| m54088_170216_085445_69337843_ccs_Otu00277_5_1588_bp | 98.4 | Eukaryota;Opisthokonta;Holozoa;Metazoa;Animalia;Eumetazoa;Bilateria;Nematoda;Chromadorea;Monhysterida | 1;1;1;1;1;1;1;1;1;1 |
| m54088_170216_085445_70320537_ccs_Otu00568_2_1588_bp | 89.8 | Eukaryota;SAR;Stramenopiles;Peronosporomycetes | 1;1;1;0.989333 |
| m54088_170216_085445_70386622_ccs_Otu00427_3_1588_bp | 89.3 | Eukaryota;Opisthokonta;Nucletmycea;Fungi;Chytridiomycota;Incertae_Sedis;Chytridiomycetes;Spizellomycetales;Olpidiaceae;Olpidium | 1;1;1;1;0.539971;0.539971;0.539971;0.539971;0.539971;0.539971 |
| m54088_170216_085445_70583078_ccs_Otu00215_8_1588_bp | 99 | Eukaryota;Opisthokonta;Nucletmycea;Fungi;Dikarya;Basidiomycota;Agaricomycotina;Agaricomycetes;Thelephorales | 1;1;1;1;1;1;1;1;0.596462 |
| m54088_170216_085445_70975772_ccs_Otu00485_2_1588_bp | 88 | Eukaryota;SAR;Alveolata;Apicomplexa;Conoidasida;Gregarinasina | 1;1;1;1;1;1 |
| m54088_170216_085445_71631554_ccs_Otu00603_2_1588_bp | 97.2 | Eukaryota;SAR;Alveolata;Ciliophora;Intramacronucleata;Spirotrichea;Hypotrichia | 0.989454;0.989454;0.989454;0.989454;0.989454;0.989454;0.989454 |
| m54088_170216_085445_71893500_ccs_Otu00356_4_1588_bp | 88.9 | Eukaryota;Opisthokonta;Holozoa;Metazoa;Animalia;Eumetazoa;Bilateria;Nematoda;Chromadorea;Rhabditida;Oscheius | 0.997614;0.997614;0.997614;0.997614;0.997614;0.997614;0.997614;0.997614;0.997614;0.997614;0.606565 |
| m54088_170216_085445_71959084_ccs_Otu00576_2_1588_bp | 95.1 | Eukaryota;SAR;Alveolata;Dinoflagellata;Dinophyceae;Gymnodiniphycidae_Gymnodinium_clade | |
| m54088_170216_085445_72024439_ccs_Otu00526_2_1588_bp | 86 | Eukaryota;SAR;Alveolata;Apicomplexa;Conoidasida;Gregarinasina;Eugregarinorida;Syncystis;mirabilis | 1;1;1;1;1;1;0.724335;0.724335;0.724335 |
| m54088_170216_085445_72024988_ccs_Otu00326_4_1588_bp | 95.1 | Eukaryota;Amoebozoa;Tubulinea;Euamoebida;BOLA868 | 0.999994;0.999994;0.999994;0.999994;0.999994 |
| m54088_170216_085445_72089952_ccs_Otu00394_3_1588_bp | 95.7 | Eukaryota;Opisthokonta;Nucletmycea;Fungi;Dikarya;Basidiomycota;Agaricomycotina;Tremellomycetes;Tremellales | 0.985274;0.985274;0.985274;0.985274;0.985274;0.985274;0.985274;0.985274;0.985274 |
| m54088_170216_085445_72155277_ccs_Otu00180_10_1588_bp | 95.8 | Eukaryota;Opisthokonta;Nucletmycea;Fungi;Dikarya;Basidiomycota;Agaricomycotina;Agaricomycetes;Agaricales;Tricholomataceae;Camarophyllopsis | 1;1;1;1;1;1;1;1;1;1;0.843715 |
| m54088_170216_085445_72548476_ccs_Otu00053_43_1588_bp | 97.8 | Eukaryota;SAR;Rhizaria;Cercozoa;Cercomonadidae;Cercomonas;longicauda | 1;1;1;1;1;0.554665;0.554665 |
| m54088_170216_085445_73007363_ccs_Otu00567_2_1588_bp | 93.3 | Eukaryota;SAR;Stramenopiles;Ochrophyta;Xanthophyceae;Tribonematales | 1;1;1;1;0.549815;0.549815 |
| m54088_170216_085445_73007549_ccs_Otu00557_2_1588_bp | 81 | Eukaryota;Opisthokonta;Nucletmycea;Fungi;Cryptomycota | 0.971924;0.971924;0.971924;0.953354;0.915605 |
| m54088_170216_085445_73138646_ccs_Otu00172_10_1588_bp | 99.1 | Eukaryota;Archaeplastida;Chloroplastida;Chlorophyta;Chlorophyceae;Chlorosarcinopsis;A_KF2011 | 1;1;1;1;1;0.604649;0.604649 |
| m54088_170216_085445_73269546_ccs_Otu00587_2_1588_bp | 92.4 | Eukaryota;SAR;Rhizaria;Cercozoa;Vampyrellidae | 0.999964;0.999964;0.999964;0.999964;0.999964 |
| m54088_170216_085445_73663423_ccs_Otu00537_2_1588_bp | 87.8 | Eukaryota;SAR;Alveolata;Apicomplexa;Conoidasida;Gregarinasina;Archigregarinorida;Selenidium | 1;1;1;1;1;1;1;1 |
| m54088_170216_085445_74187639_ccs_Otu00591_2_1588_bp | 99.1 | Eukaryota;SAR;Alveolata;Ciliophora;Intramacronucleata;Spirotrichea;Hypotrichia;Anteholosticha;monilata | 0.999893;0.999893;0.999893;0.999893;0.999893;0.999893;0.999893;0.9915;0.9915 |
| m54088_170216_085445_8061165_ccs_Otu00258_6_1588_bp | 78.8 | Eukaryota;Opisthokonta;Nucletmycea;Fungi;Zoopagomycota | 0.987461;0.987397;0.987397;0.987397;0.987397 |
| m54088_170216_085445_8192769_ccs_Otu00166_11_1588_bp | 97.3 | Eukaryota;SAR;Rhizaria;Cercozoa;Cercomonadidae | 0.999974;0.999974;0.999974;0.999974;0.999974 |
| m54088_170216_085445_8913735_ccs_Otu00156_11_1588_bp | 98.6 | Eukaryota;SAR;Stramenopiles;Ochrophyta;Diatomea;Bacillariophytina;Bacillariophyceae;Fistulifera | 0.999998;0.999998;0.999998;0.999998;0.999998;0.999998;0.999998;0.999998 |
| m54088_170216_085445_9372216_ccs_Otu00621_2_1588_bp | 94 | Eukaryota;SAR;Alveolata;Ciliophora;Intramacronucleata;Conthreep;Prostomatea;Prorodon | 1;1;1;1;1;1;1;1 |
| m54088_170216_085445_9437773_ccs_Otu00348_4_1588_bp | 86.6 | Eukaryota;SAR;Alveolata;Apicomplexa;Conoidasida;Gregarinasina;Eugregarinorida;Syncystis;mirabilis | 1;1;1;1;1;1;0.661653;0.661653;0.661653 |
